# Precise Transcript Targeting by CRISPR-Csm Complexes

**DOI:** 10.1101/2022.06.20.496908

**Authors:** David Colognori, Marena Trinidad, Jennifer A. Doudna

**Affiliations:** Department of Molecular and Cell Biology, University of California, Berkeley, CA, USA; Innovative Genomics Institute, University of California, Berkeley, Berkeley, CA, USA; Howard Hughes Medical Institute, University of California, Berkeley, Berkeley, CA, USA; Department of Chemistry, University of California, Berkeley, Berkeley, CA, USA; California Institute for Quantitative Biosciences (QB3), University of California, Berkeley, Berkeley, CA, USA; Molecular Biophysics and Integrated Bioimaging Division, Lawrence Berkeley National Laboratory, Berkeley, CA, USA; Gladstone Institutes, San Francisco, CA, USA

**Keywords:** CRISPR-Cas, Type III, Csm, Cas13, RNA knockdown, RNA imaging

## Abstract

Robust and precise transcript targeting in mammalian cells remains a difficult challenge using existing approaches due to inefficiency, imprecision, and subcellular compartmentalization. Here, we show that the CRISPR-Csm complex, a multi-protein effector from type III CRISPR immune systems in prokaryotes, provides surgical RNA ablation of both nuclear and cytoplasmic transcripts. As part of the most widely occurring CRISPR adaptive immunity pathway, CRISPR-Csm uses a programmable RNA-guided mechanism to find and degrade target RNA molecules without inducing indiscriminate *trans*-cleavage of cellular RNAs, giving it an important advantage over the CRISPR-Cas13-family enzymes. Using single-vector delivery of the *S. thermophilus* Csm complex, we observe high-efficiency RNA knockdown (90-99%) and minimal off-target effects in human cells, outperforming existing technologies including shRNA- and Cas13-mediated knockdown. We also find that catalytically inactivated Csm achieves specific and durable RNA binding, a property we harness for live-cell RNA imaging. These results establish the feasibility and efficacy of multi-protein CRISPR-Cas effector complexes as RNA-targeting tools in eukaryotes.

## INTRODUCTION

The ability to alter RNA and protein levels in cells and organisms without making permanent changes to DNA has proven invaluable for both basic research and therapeutics. For the past two decades, targeted RNA knockdown in eukaryotes has been accomplished by RNA interference (RNAi), an approach whereby small interfering RNAs (siRNAs) direct endogenous Argonaute nucleases to cleave complementary target RNAs (Wilson and Doudna 2013; Carthew and Sontheimer 2009). However, RNAi can cause unintended cleavage of targets carrying partial sequence complementarity, especially when this complementarity occurs within the nucleating “seed” region (nucleotides 2-7) of the siRNA (Jackson et al. 2003, 2006; Birmingham et al. 2006). Furthermore, siRNAs are inefficient at targeting nuclear RNAs since the RNAi machinery localizes primarily to the cytoplasm (Behlke 2016; Zeng and Cullen 2002). Finally, RNAi is incompatible with certain eukaryotic model systems, including budding yeast which lack RNAi machinery (Aravind et al. 2000; Nakayashiki, Kadotani, and Mayama 2006) and zebrafish embryos which suffer nonspecific developmental defects (Zhao et al. 2001; Oates, Bruce, and Ho 2000). Thus, there has been ongoing interest in the development of new RNA knockdown tools with higher specificity and broader targeting capability.

CRISPR (clustered regularly interspaced short palindromic repeats)-Cas proteins, which comprise adaptive defense systems against infectious agents in prokaryotes (Barrangou et al. 2007; Brouns et al. 2008), operate as programmable DNA or RNA nucleases (Jinek et al. 2012; Cong et al. 2013; Mali et al. 2013). Similar to RNAi, Cas nucleases utilize small RNAs (CRISPR RNAs [crRNAs]) to recognize nucleic acid targets via base-pairing complementarity. One such nuclease, Cas13, has gained attention as a new RNA-cleavage tool for use in eukaryotes (Abudayyeh et al. 2017, 2016; Konermann et al. 2018). However, unlike Argonaute proteins that cut only complementary RNAs in *cis* (Song et al. 2004), Cas13 also degrades nearby non-complementary RNAs in *trans* (East-Seletsky et al. 2016; Abudayyeh et al. 2016) (Fig. 1A). This is because the nuclease domains of Cas13 are located away from the crRNA:target binding pocket on an exposed surface of the protein (Zhang et al. 2018; Slaymaker et al. 2019; L. Liu, Li, Ma, et al. 2017; L. Liu, Li, Wang, et al. 2017). Cas13’s *trans*-cleavage activity is readily detectable in *vitro*, where it has been exploited for viral RNA detection tools (Gootenberg et al. 2017; East-Seletsky et al. 2016; Gootenberg et al. 2018; T. Y. Liu et al. 2021; Fozouni et al. 2021). In bacteria, *trans*-cleavage leads to stalled cell growth or cell death (abortive infection) (Meeske, Nakandakari-Higa, and Marraffini 2019), which is now believed to be Cas13’s primary mode of defensive action against viral infection. Only recently, however, has evidence mounted that Cas13 exhibits *trans*-cleavage activity in eukaryotic cells, causing cytotoxicity and/or cell death (Q. Wang et al. 2019; Özcan et al. 2021; Ai, Liang, and Wilusz 2022; Tong et al. 2021; Shi et al. 2021). The convolution of cis- and trans-RNA cutting effects have made it difficult to interpret results obtained using Cas13, and call into question its utility as a tool for specific RNA knockdown.

**Figure 1.**
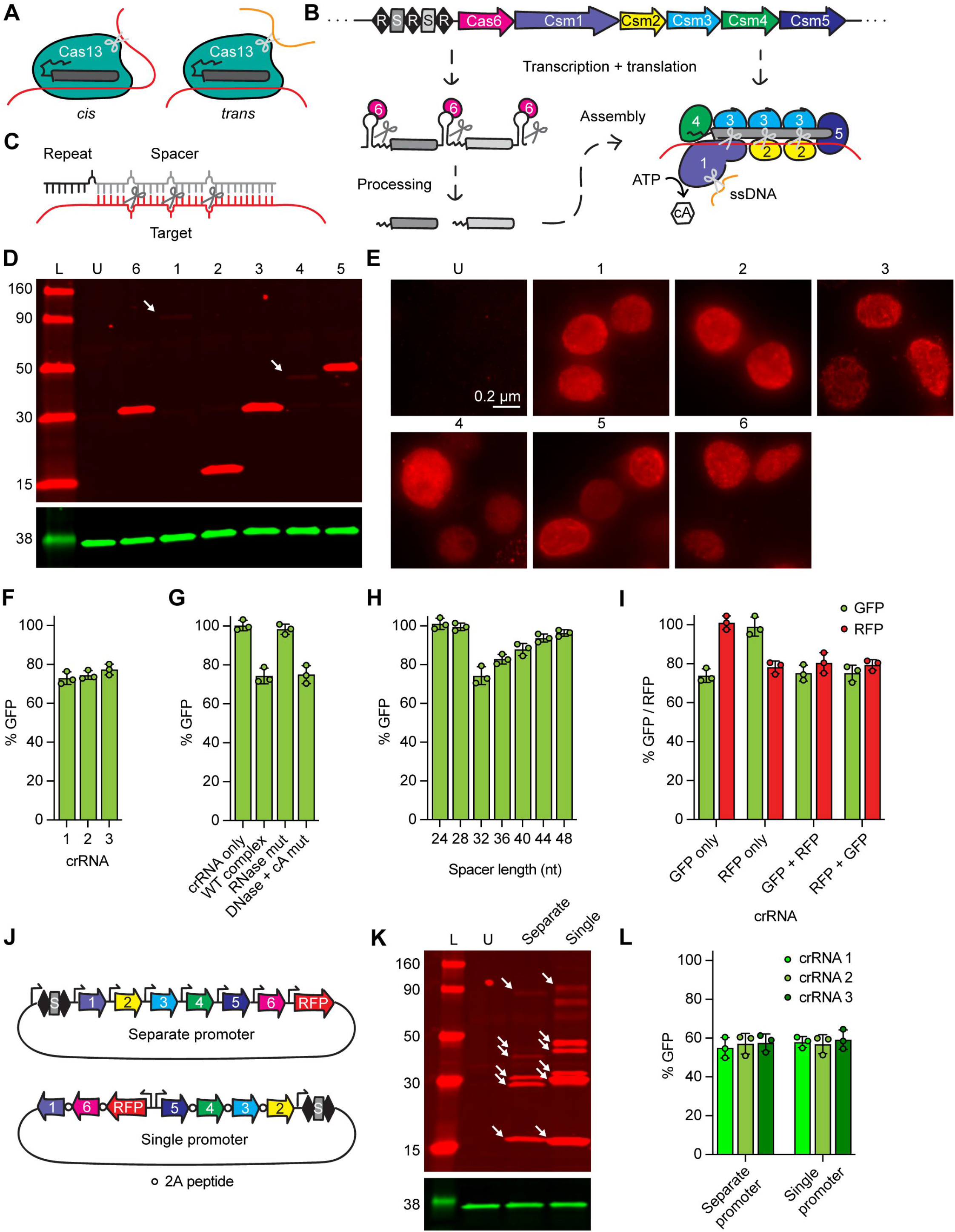
Establishing an all-in-one Type III CRISPR-Cas system in mammalian cells. A. Diagram showing cis- and trans-cleavage of Cas13. B. Diagram showing Type III-A CRISPR-Cas locus. The CRISPR array is transcribed and processed into mature crRNAs by Cas6, which assemble with Csm proteins. cA, cyclic oligoadenylate. C. Close-up of crRNA:target binding and cleavage, showing the 6-nt spacing pattern. D. Western blot showing proper size and expression of Csm proteins (red) in HEK293T cells. GAPDH shown as loading control (green). Arrows indicate faint bands. L, ladder; U, untransfected; 1-6, Csm1-5 and Cas6. E. Immunofluorescence showing expression and nuclear localization of Csm proteins in HEK293T cells. Labeling same as in (D). F. Relative GFP fluorescence (= MFI targeting crRNA / MFI non-targeting crRNA) of HEK293T-GFP cells transfected with the indicated crRNAs, measured by flow cytometry. Error bars indicate mean ± standard deviation of 3 biological replicates. G. Relative GFP fluorescence of HEK293T-GFP cells transfected with the indicated protein complexes (or crRNA and Cas6 only), measured by flow cytometry. H. Relative GFP fluorescence of HEK293T-GFP cells transfected with crRNAs of indicated spacer length, measured by flow cytometry. I. Relative GFP and RFP fluorescence of HEK293T-GFP/RFP cells transfected with the indicated crRNAs (individual or multiplexed), measured by flow cytometry. J. Diagram showing all-in-one delivery vector designs. K. Western blot showing proper size and expression of Csm proteins (red) in HEK293T cells. GAPDH shown as loading control (green). Arrows indicate faint bands. Labeling same as in (D). L. Relative GFP fluorescence of HEK293T-GFP cells transfected with the indicated delivery vectors and crRNAs, measured by flow cytometry.

Despite their higher prevalence in nature (Makarova et al. 2015), multi-subunit Cas effectors have been harnessed only rarely as tools in eukaryotes (with few exceptions (Pickar-Oliver et al. 2019; Cameron et al. 2019; Chen et al. 2020)) due to their component complexity. Nonetheless, their biochemical activities and well-studied structural properties make the type III RNA-targeting CRISPR-Csm complexes of particular interest as potential transcript targeting tools. The multi-protein Csm complex comprises five subunits (Csm1-5) in varying stoichiometries and relies on an additional protein, Cas6, for processing the precursor crRNA (Staals et al. 2014; Zhu et al. 2018; You et al. 2019; Tamulaitis et al. 2014; Jia et al. 2019; Guo et al. 2019; T. Y. Liu, Iavarone, and Doudna 2017; Mogila et al. 2019) (Fig. 1B). The crRNA lies at the core of the complex, with Csm1 and Csm4 binding the 5’ end, Csm5 binding the 3’ end, and multiple copies of Csm2 and Csm3 wrapping around the center. The complex contains a groove along its length into which target RNAs can enter and hybridize to the variable spacer region of the crRNA. Csm1 and Csm4 specifically recognize the 5’ region of the crRNA derived from the CRISPR repeat. Each Csm3 subunit has ribonuclease (RNase) activity, leading to multiple cleavage sites within the target RNA spaced six nucleotides apart (Fig. 1C). Csm1 functions as a non-specific single-stranded DNase (Kazlauskiene et al. 2016; Samai et al. 2015) and a cyclic oligoadenylate (cA) synthase (Niewoehner et al. 2017; Kazlauskiene et al. 2017) (Fig. 1B). The ssDNase activity is thought to defend against ssDNA or actively transcribed (R-looped) foreign genomes (Kazlauskiene et al. 2016; Samai et al. 2015), while the latter acts as a second messenger that activates downstream defense effectors in *trans*, such as the RNase Csm6 (Niewoehner et al. 2017; Kazlauskiene et al. 2017). Importantly, all three catalytic activities are carried out by independent domains of the Csm complex and can be individually ablated.

Csm is an attractive RNA knockdown tool over current methods. A self-contained system found only in prokaryotes, it can be orthogonally introduced into eukaryotes without intersecting host RNA regulatory pathways like RNAi. Furthermore, unlike RNAi, it can be localized to the nucleus and used to target nuclear noncoding RNAs as well pre-mRNAs. Compared to Cas13, Csm cleaves only in *cis* within the crRNA:target complementary region and thus does not suffer *trans*-cleavage events. Additionally, unlike Cas13, Csm-mediated RNA cleavage does not preferentially occur at a particular nucleotide base (e.g. U) (Abudayyeh et al. 2016; Gootenberg et al. 2018), nor is directly influenced by sequence flanking the target (e.g. tag:anti-tag complementarity) (B. Wang et al. 2021; Tamulaitis et al. 2014). In this work, we demonstrate the utility of the Csm system as a highly efficient, specific, and versatile RNA knockdown tool in eukaryotes.

## RESULTS

### Establishing an all-in-one Type III CRISPR-Cas system in mammalian cells

We chose the Type III-A Csm complex from *Streptococcus thermophilus* for several reasons: 1. it has been extensively characterized biochemically, structurally, and in bacteria (Staals et al. 2014; Zhu et al. 2018; You et al. 2019; Tamulaitis et al. 2014; Jia et al. 2019; Guo et al. 2019; T. Y. Liu, Iavarone, and Doudna 2017; Mogila et al. 2019), 2. functions optimally at 37C, 3. has been demonstrated to work in zebrafish upon ribonucleoprotein (RNP) microinjection (Fricke et al. 2020), and 4. has fewer components than the analogous Type III-B Cmr complex (Staals et al. 2013). We began by verifying proper expression of each individual protein component (Csm1-5 and Cas6) in immortalized human embryonic kidney (HEK293T) cells. Proteins were human-codon-optimized, N-terminally FLAG-tagged for detection, and expressed from a CMV promoter. While RNAi operates in the cytoplasm where mRNAs mainly reside, we chose to localize each Csm component to the nucleus through the addition of an N-terminal SV40 nuclear localization signal (NLS) so as to target nuclear RNAs as well as pre-mRNAs prior to export. Following transient transfection, Western blot (Fig. 1D) and immunofluorescence staining (Fig. 1E) verified proper size, expression, and nuclear localization of each protein.

To test our system, we targeted eGFP (henceforth “GFP”) mRNA in a GFP-expressing HEK293T cell line. Seven plasmids individually expressing Csm1-5, Cas6, and either a GFP-targeting or non-targeting crRNA from a U6 promoter were co-transfected into cells, and GFP fluorescence assayed by flow cytometry 48 hr post-transfection (Fig. S1A). Note that this strategy does not allow for any means to select cells into which all plasmids were successfully delivered, and will thus under-report knockdown (KD) efficiency. % GFP KD was calculated by dividing the mean fluorescence intensity (MFI) of cells transfected with the GFP-targeting crRNA by that of cells transfected with the non-targeting crRNA (Fig. S1B). ∼25% KD was observed using any of three crRNAs targeting different regions of the GFP ORF (Fig. 1F). Importantly, no KD was seen after transfecting the GFP-targeting crRNA and its processing factor (Cas6) alone (Fig. 1G), indicating that KD was not due to an antisense RNA effect. Furthermore, whereas ablating DNase (H15A, D15A) and cA synthase (D577A, D578A) activities in Csm1 did not affect GFP KD, ablating RNase activity (D33A) in Csm3 completely abolished GFP KD (Fig. 1G), indicating that RNase activity is necessary and sufficient for KD.

Next, we examined crRNA parameters. Naturally occurring spacers for *Sth*Csm crRNAs range from ∼30-45 nucleotides (nt) in length, although *in vitro*, spacers as short as 27 nt are sufficient to trigger all three catalytic activities (You et al. 2019). We varied the GFP-targeting spacer length from 24-48 nt in increments of 4 and assayed GFP KD. A length of 32 nt yielded the highest KD for the crRNA tested (Fig. 1H), with no KD seen for lengths ≤28 nt, and diminishing KD seen for lengths ≥32 nt. A more large-scale analysis must be performed to determine whether optimal spacer length differs from sequence to sequence. Next, we examined the potential to multiplex crRNAs against multiple targets. We encoded two crRNAs within a single array--one targeting GFP and the other targeting mCherry (henceforth “RFP”)--and examined KD of GFP and RFP in a HEK293T cell line expressing both (Fig 1I). ∼25% KD was achieved for both GFP and RFP regardless of the order of crRNAs in the array (GFP-RFP or RFP-GFP), comparable to KD efficiency when targeting GFP or RFP alone. Together, these results demonstrate broad multiplexing capability for the Csm system.

With the Csm system up and running, we sought to simplify its delivery by consolidating all components into a single vector. For this, we pursued two approaches concurrently: 1. expression of each protein from separate promoters, or 2. expression of all proteins from a single bidirectional promoter separated by 2A peptides (Fig. 1J). We also included RFP in the plasmid backbone to allow identification of transfected cells and thus more accurate measurement of KD efficiency (Fig. S1C). After re-confirming proper expression of all protein components by Western blot for both plasmids (Fig. 1K), we found both strategies (after optimizing the order of proteins in the single-promoter arrangement) led to ∼50% GFP KD in transfected cells (Fig. 1L). In summary, the single-promoter design is well-equipped for promoter-swapping and thus use in specific cell types or other eukaryotic systems, while the modular design of the separate-promoter vector allows for easy swapping or modification of individual Csm components.

### Robust knockdown of endogenous nuclear and cytoplasmic RNAs

Thus far, we have only used Csm to KD highly overexpressed, heterologous GFP/RFP transgenes and assayed KD at the protein level (half-life >24 hours (Corish and Tyler-Smith 1999)), which may not accurately reflect abundance at the RNA level. We thus sought to target endogenous transcripts and assay RNA KD directly. We chose to target a panel of three nuclear noncoding RNAs (XIST, MALAT1, NEAT1) and eight cytoplasmic mRNAs (BRCA1, TARDBP, SMARCA1, CKB, ENO1, MECP2, UBE3A, SMAD4) (Fig. 2A) of varying abundances (Fig. 2B), testing three individual crRNAs for each. HEK293T cells were transfected, transfected (RFP-positive) cells were isolated by FACS after 48 hr, total cell RNA extracted, and RNA KD assayed by RT-qPCR (Fig. S1C,S2A). To our surprise, we achieved >90% KD for all eleven RNAs with at least one crRNA, compared to non-targeting crRNA control (Fig. 2A). These results demonstrate the Csm system to be a highly robust and efficient RNA KD tool for not only cytoplasmic but also nuclear RNAs, which are typically recalcitrant to KD by conventional RNAi methods (Behlke 2016).

**Figure 2.**
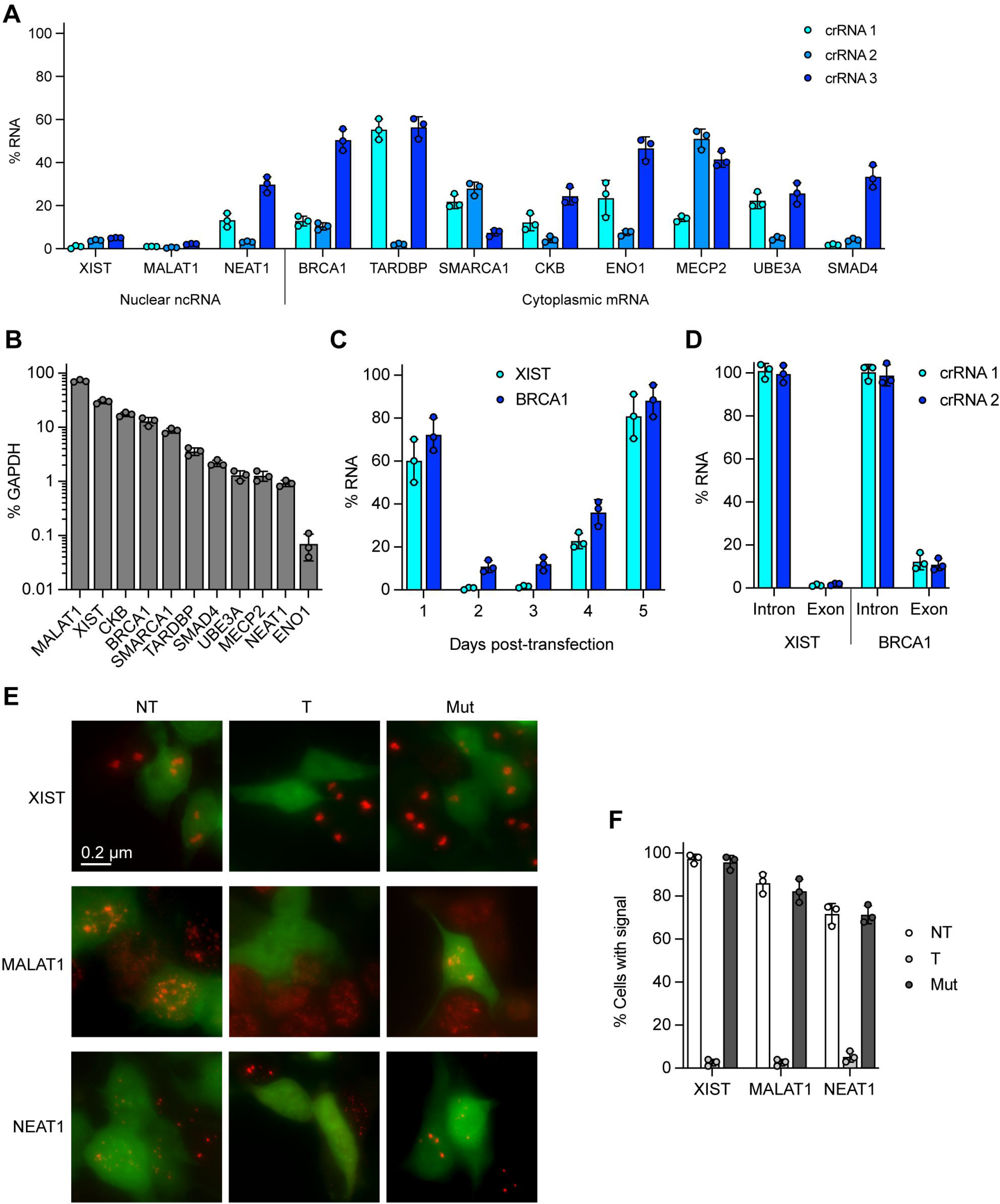
Robust knockdown of endogenous nuclear and cytoplasmic RNAs. A. Relative RNA abundance (normalized to non-targeting crRNA) of the indicated targets in HEK293T cells transfected with the indicated crRNAs, measured by RT-qPCR. Error bars indicate mean ± standard deviation of 3 biological replicates. B. Relative RNA abundance (normalized to GAPDH) of the indicated targets in untransfected HEK293T cells, measured by RT-qPCR. C. Relative RNA abundance (normalized to non-targeting crRNA) of XIST and BRCA1 in HEK293T cells at the indicated times post crRNA transfection, measured by RT-qPCR. D. Relative RNA abundance (normalized to non-targeting crRNA) of XIST and BRCA1 in HEK293T cells transfected with intron- or exon-targeting crRNAs, measured by RT-qPCR. E. RNA FISH (red) for the indicated targets in HEK293T cells transfected with targeting (T) or non-targeting (NT) crRNA, and RNase-active or -inactive (Mut) protein complex. Untransfected cells serve as internal control for transfected (green) cells. F. Quantification of (E). 100 transfected cells were counted for each condition.

To examine KD kinetics, we repeated the above RT-qPCR experiment for two of the RNA targets (XIST, BRCA1) across a 5-day time-course. KD peaked 2-3 days post-transfection and waned thereafter (Fig. 2C), as might be expected from the transient transfection method used to deliver the Csm into cells. We also compared KD efficiency of crRNAs targeting intronic versus exonic regions for the same two RNAs (Fig. 2D). Targeting introns did not lead to any noticeable reduction in mature RNA levels, possibly because their excision from pre-mRNA occurs more rapidly than their binding and cleavage by Csm.

To corroborate RNA KD with an orthogonal method, we performed RNA FISH for all three nuclear noncoding RNAs, which are easily visualized and display characteristic morphologies. HEK293T cells were transfected with Csm plasmid carrying a GFP reporter (to identify transfected cells) and either a targeting or non-targeting crRNA, and assayed by RNA FISH after 48 hr (Fig. S2B). XIST, MALAT1, and NEAT1 were all readily detected when delivering a non-targeting crRNA control (Fig. 2E,F). By contrast, use of a single targeting crRNA abolished all visible signal for each target RNA in transfected cells (GFP-positive cells), whereas signal was detected in untransfected (GFP-negative) cells. For further validation, delivery of targeting crRNA with RNase-inactivate Csm fully restored detection of each target RNA. Thus, we demonstrate near-complete target RNA KD with active Csm complexes by both molecular and microscopy-based techniques.

### RNA knockdown with minimal off-targets or cytotoxicity

We performed RNA-sequencing to examine potential off-target effects of Csm-mediated KD in cells. For comparison with other established KD technologies, RNA-seq was also performed for Cas13 (RfxCas13d) and RNAi (shRNA) -mediated KD (Wei et al. 2021; Bofill-De Ros and Gu 2016; Wessels et al. 2020). XIST, MALAT1, CKB, or SMAD4 was depleted for 48 hr using Csm, Cas13, or shRNA using crRNAs/shRNAs targeting the same complementary sequence for each transcript (Fig. S3A). Scatterplots comparing differential transcript levels between Csm-treated and untreated samples showed significant KD of the target transcript (CKB, MALAT1 shown) with few other differentially expressed genes (≥2-fold change, indicated in red) (Fig. 3A,B). Cas13-treated samples showed significant KD of the target but with thousands of differentially expressed off-target genes. shRNA-treated samples showed variable KD depending on whether the target was cytoplasmic (CKB) or nuclear (MALAT1), with few other differentially expressed genes. Similar trends were seen for all four target transcripts and both crRNAs/shRNAs per transcript (Fig. 3C). Examination of RNA-seq read coverage confirmed that target KD was transcript-wide and not only localized near the Csm cleavage sites--unsurprising given exonucleotic RNA degradation pathways in mammalian cells (Houseley and Tollervey 2009) (Fig. 3D,E). Hence, unlike Cas13, Csm- and shRNA-mediated RNA KD has few off-target effects in human cells.

**Figure 3.**
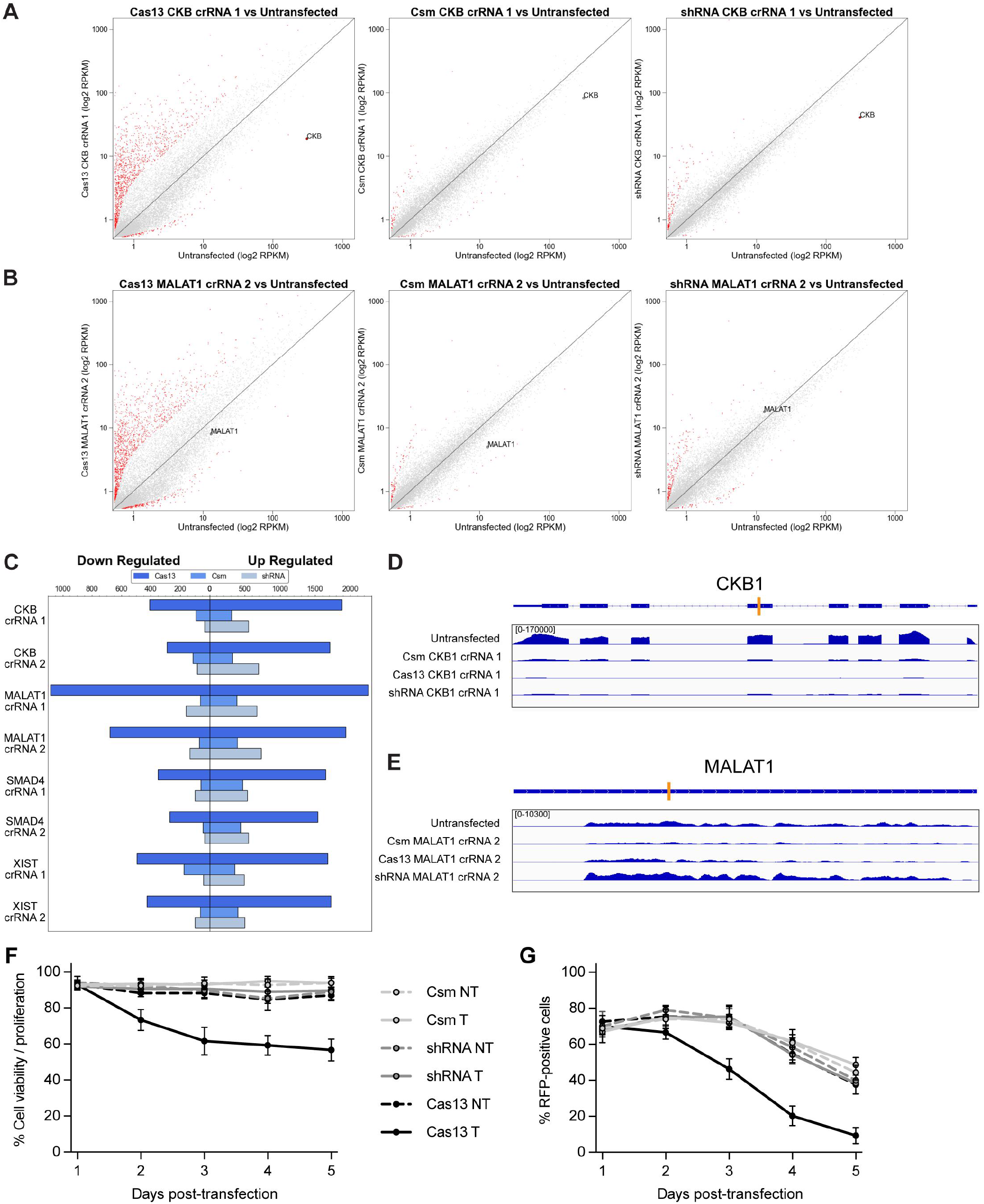
RNA knockdown with minimal off-targets or cytotoxicity. A,B. Scatterplots showing differential transcript levels between Csm, Cas13, or shRNA-treated cells targeting CKB (A) or MALAT1 (B) versus untreated cells. Target transcript is indicated; red dots indicate differentially regulated off-targets (≥2-fold change). C. Quantification of significantly up- or down-regulated genes (≥2-fold change) for each sample. D,E. RNA-seq read coverage across target transcripts, CKB (D) or MALAT1 (E), in Csm, Cas13, or shRNA-treated cells. Orange bar indicates location of crRNA/shRNA target sequence. F. Relative cell viability and proliferation (normalized to untransfected cells) of HEK293T cells at the indicated times post transfection with the indicated targeting (T) or non-targeting (NT) plasmids, measured by WST-1 assay. G. Relative abundance of RFP-positive HEK293T cells (normalized to untransfected cells) at the indicated times post transfection with the indicated targeting (T) or non-targeting (NT) plasmids, measured by flow cytometry.

Other RNA-targeting CRISPR-Cas systems such as Cas13 suffer from severe cytotoxic effects due to inherent *trans*-cleavage activity of the Cas effector (Q. Wang et al. 2019; Özcan et al. 2021; Ai, Liang, and Wilusz 2022; Tong et al. 2021; Shi et al. 2021). Type III systems do not exhibit such *trans*-activity and are thus poised to offer robust RNA KD without such toxicity. To check this, we tracked cell proliferation/viability using the WST-1 assay across a time-course after transfecting cells with Csm, Cas13, or shRNA constructs (Fig. 3F). Whereas Cas13-treated cells exhibited a significant decrease in proliferation/viability, Csm- or shRNA-treated cells were unaffected. This decrease in proliferation/viability was accompanied by a more rapid decrease over time in the proportion of RFP-positive (transfected) cells for the Cas13-treated population compared to the Csm- or shRNA-treated population (Fig. 3G). Taken together, these results suggest that, unlike Cas13, Csm and shRNAs do not cause pronounced toxicity in cells. In summary, Csm-mediated KD has the benefits of both Cas13 and shRNA strategies without the accompanying drawbacks: the robust KD and nuclear-targeting capability of Cas13, along with the minimal off-targets and cytotoxicity of shRNAs.

### Live-cell RNA imaging without genetic manipulation

Tracking RNA in live cells remains a difficult task, often requiring genetic insertion of aptamer sequences into the RNA target, which is both laborious and potentially disruptive to RNA function and/or regulation (George et al. 2018). Fluorescently tagged programmable RNA-binding proteins such as catalytically inactivated Cas13 have recently been adopted for such purposes (Abudayyeh et al. 2017; H. Wang et al. 2019; Yang et al. 2019). We asked whether the Csm complex could similarly be used to track RNA targets in live cells. To test this, we fused GFP to the C-terminus of catalytically inactivated Csm3 in our vector (Fig. 4A). We chose this super-stoichiometric subunit (≥3 per complex) in order to increase the signal-to-noise ratio of complexed, target-bound Csm over unbound, unassembled subunits (Fig. 4B). To visualize XIST RNA, we targeted its “Repeat A” region with a single crRNA predicted to bind 8 times per transcript, further increasing signal. Whereas a non-targeting control crRNA led to only background nuclear fluorescence, the XIST-targeting crRNA led to a strong cloud-like signal in most cells (Fig. 4C,D), characteristic of XIST RNA and phenocopying what we previously observed by XIST RNA FISH (Fig. 2E). Similar results were obtained for MALAT1 and NEAT1 RNAs, even with crRNAs predicted to bind only once per target transcript (Fig. 4C,D). Multiplexing several crRNAs against the same target may further improve signal-to-noise, especially for targets of low abundance. Thus, fluorescently-tagged Csm can be used for easy visualization of RNA in living cells.

**Figure 4.**
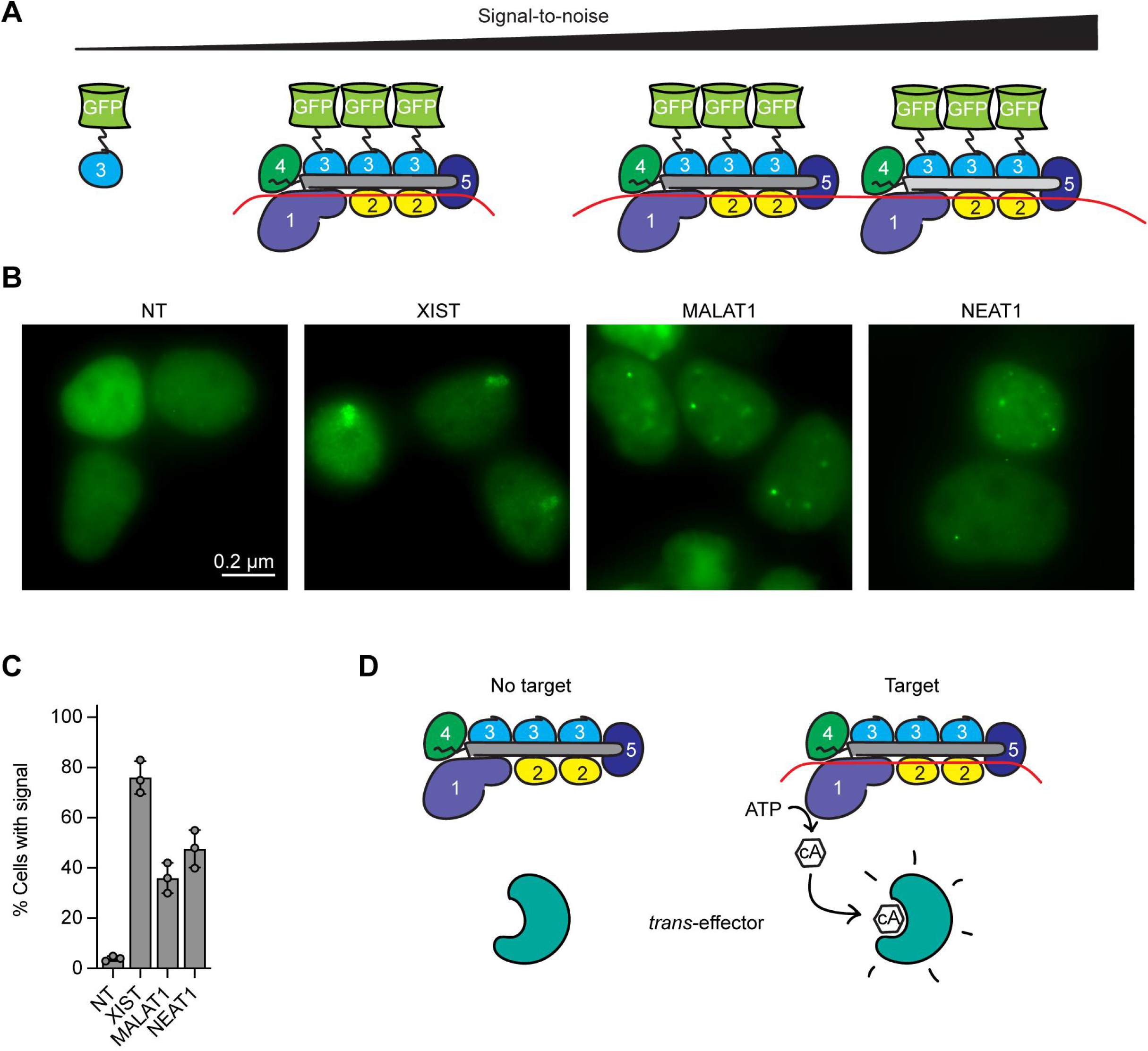
Live-cell RNA imaging without genetic manipulation. A. Diagram showing Csm-GFP fusion. Signal-to-noise increases from left to right, from unassembled Csm3, to target-bound Csm complexes, to multiplexed target-bound complexes. B. Live-cell fluorescent imaging of HEK293T cells transfected with Csm-GFP protein complex and the indicated crRNAs. C. Quantification of (B). 100 transfected cells were counted for each condition. D. Diagram showing RNA sequence-dependent activation of downstream effectors by the Csm complex.

## DISCUSSION

Here, we have shown that the Type III-A Csm complex from *Streptococcus thermophilus* is a powerful tool for eukaryotic RNA knockdown. Significantly, knockdown of both nuclear noncoding RNAs as well as cytoplasmic mRNAs was achieved with high efficiency (90-99%) and specificity (10-fold fewer off-targets than Cas13), outperforming currently used RNA KD technologies. More importantly, RNA KD was not accompanied by detectable cytotoxicity, unlike Cas13-based KD which suffers from inherent *trans*-cleavage activity (Q. Wang et al. 2019; Özcan et al. 2021; Ai, Liang, and Wilusz 2022; Tong et al. 2021).

Recently, *St*Csm was shown to be effective at depleting GFP mRNA upon microinjection of bacterially purified RNP into transgenic zebrafish embryos (Fricke et al. 2020). While demonstrating proof of principle, RNP delivery of multi-subunit CRISPR-Cas effectors is not ideal for several reasons: 1. delivery is often difficult and short-lived compared to DNA-delivery methods, 2. the RNP may be unstable and prone to disassembly, and 3. for every new crRNA, the entire RNP must be re-purified from bacteria or reconstituted from individually purified subunits in the proper ratio. We have overcome these hurdles of multi-component CRISPR-Cas systems by encoding all necessary parts in a single deliverable plasmid.

Recently, a single-protein Type III effector, Cas7-11, was characterized as a potential tool for RNA KD in eukaryotes (Özcan et al. 2021). This effector is interesting from an evolutionary and structural standpoint in that it appears to have arisen from fusion of the canonical Type III subunits into a single large polypeptide. While simpler introduce into eukaryotes, Cas7-11’s demonstrated RNA KD efficiency was 50-75% for most targets, making it less practical as a tool. Meanwhile, a recent pre-print of engineered high-fidelity (hf)Cas13 variants reports to have mitigated *trans-*cleavage activity (and thus cytotoxicity) while preserving on-target activity (Tong et al. 2021)--though a mechanistic explanation for this remains unclear. Both Cas7-11 and hfCas13 await further characterization before widespread use.

A key advantage of our approach over existing technologies is the ability to target RNAs in the nucleus. We were able to achieve ∼99% KD for 3 biologically significant nuclear ncRNAs (XIST, MALAT1, NEAT1). Nuclear RNAs are notoriously difficult to KD, often requiring expensive chemically modified antisense oligos to direct RNase H-mediated cleavage (Behlke 2016). However, the increased stability of these oligos often leads to unexpected off-target hybridization as well as cytotoxic effects. Aside from long ncRNAs, nuclear targeting may prove useful for the study of other ncRNA species such as eRNAs, tRNAs, rRNAs, circRNAs, miRNAs, and snoRNAs. For instance, it will be interesting to see whether targeting introns with miRNA or snoRNA clusters causes their degradation prior to processing, or if targeting particular exons alters the abundance of mRNA splice isoforms.

Another advantage of our system is its ease of multiplexing. Multiple spacers can be cloned into the CRISPR array and processed into individual mature crRNAs by Cas6. This allows for pooled screening, either by encoding crRNAs against multiple targets at once or encoding multiple crRNAs against the same target. The latter may enable robust KD on the first try without the need to individually screen multiple crRNAs against a target. An unexpected observation was the titratable nature of KD with increasing spacer length. This may allow for easy tunability of KD, rather than all-or-none, when studying concentration-dependent effects of gene products. It will be interesting to see what accounts for this phenomenon, though the heterogenous nature of Csm complexes (varying stoichiometries of Csm2 and Csm3 depending on crRNA length) is likely involved. Perhaps cleavage activity decreases with larger complexes, or longer crRNA spacers leads to increased off-target binding, thus diluting on-target cleavage.

Csm-mediated RNA KD appears particularly robust. We achieved >50% KD for every target and every crRNA tested, with at least one of three crRNAs per target yielding >90% KD. Because, like other RNA-targeting CRISPR-Cas systems, Csm does not have any PAM requirement for target site selection, the only criteria we used were that the target be a unique sequence in the human transcriptome and the spacer avoid stretches of ≥5 consecutive Ts, which might cause premature Pol III transcriptional termination of the crRNA (Gao, Herrera-Carrillo, and Berkhout 2018). A more large-scale analysis must be performed to determine optimal spacer design criteria, and to test how different factors (melting temperature, GC-content, target site availability) influence KD efficiency.

We showed that a fluorescently tagged, catalytically inactivated Csm complex can be used for live-cell RNA visualization. By fusing GFP to the most abundant subunit, Csm3, we were able to ensure ≥3 copies of GFP per assembled complex at the target (versus 1 copy on soluble protein), yielding higher signal-to-noise achievable by single-subunit effectors like Cas13. Beyond GFP, other proteins of interest may be fused to different subunits of the Csm complex to achieve assembly at a desired stoichiometric ratio. Thus, as a multi-subunit complex, Csm offers the benefits of split-protein systems without the engineering effort. Catalytically inactivated Csm could also be used to bind specific RNA sequences, disrupting structural motifs or RNA-protein interactions of interest without manipulation at the DNA level.

Finally, this work utilized only the RNase activity of Csm while ablating its DNase and cA synthase activities. In prokaryotes, cA signaling appears to be the main defensive strategy employed by Type III systems (Rostøl and Marraffini 2019), leading to activation of various downstream effectors (Niewoehner et al. 2017; Kazlauskiene et al. 2017). These effectors range from RNases, to DNases, to transcription factors, to membrane pores (Huang and Zhu 2020; Makarova et al. 2020; Athukoralage and White 2021). cA species and reliant pathways are currently not known to exist in eukaryotes, and thus could be introduced in an orthogonal manner. By bringing Type III systems to eukaryotes, we have paved the way for co-introduction of CARF domain-containing *trans*-effectors that can be activated in an RNA sequence-dependent manner (Fig. 4E). This has important implications for development of synthetic circuits, RNA diagnostics, and reporter assays/screens *in vivo*.

## FIGURE LEGENDS

**Figure S1**

A. Diagram showing workflow for flow cytometry experiments. Delivery plasmids and recipient cell lines are indicated in each experiment.

B. Diagram showing gating procedure for flow cytometry.

C. Diagram showing gating procedure for flow cytometry and FACS in which transfected (RFP-positive) cells were enriched.

**Figure S2**

A. Diagram showing workflow for RT-qPCR experiments.

B. Diagram showing workflow for RNA FISH experiments.

**Figure S3**

A. Diagram showing workflow for RNA-seq experiments.

**Figure S4**

A. Diagram showing workflow for live-cell imaging experiments.

**Table S1**

qPCR primer, crRNA, shRNA, and FISH probe sequences used in this work.

**Table S2**

Plasmid sequences used in this work.

## METHODS

### Cell lines and culture conditions

HEK293T, HEK293T-GFP, and HEK293T-GFP/RFP cells (UC Berkeley Cell Culture Facility) were grown in medium containing DMEM, high glucose, GlutaMAX supplement, sodium pyruvate (Thermo Fisher Scientific), 10% FBS (Sigma), 25 mM HEPES pH 7.2-7.5 (Thermo Fisher Scientific), 1x MEM non-essential amino acids (Thermo Fisher Scientific), 1x Pen/Strep (Thermo Fisher Scientific), and 0.1 mM BME (Thermo Fisher Scientific) at 37C with 5% CO2. All cell lines were verified to be mycoplasma-free (abm, PCR mycoplasma detection kit).

### Plasmid construction and cloning

Csm CRISPR-Cas sequences were derived from *Streptococcus thermophilus* strain ND03 (NCBI ###). Protein sequences were human codon-optimized using online tools (GenScript), synthesized as gene blocks (IDT), modified using PCR, and cloned into custom eukaryotic expression vectors (derived from pUC19) by Golden Gate assembly, Gibson assembly (NEB), or Gibson assembly Ultra (Synthetic Genomics). Plasmids were verified by Sanger or whole-plasmid sequencing. All cloning was performed in Stbl3 *E. coli* (Thermo Fisher Scientific) to prevent recombination between repetitive sequences. crRNA/shRNA sequences are provided in Table S1. Plasmid sequences are provided in Table S2.

### DNA Transfections

1×10^6 HEK293T cells were transfected with 2.5-5 ug plasmid DNA using 7.5-15 ul FuGENE HD transfection reagent in 6-well plates as per manufacturer’s instructions. Cells were grown for 48 hr post-transfection to allow protein expression and RNA KD to occur, unless otherwise stated.

### Flow cytometry

Cell fluorescence was assayed on an Attune NxT acoustic focusing cytometer (Thermo Fisher Scientific) equipped with 488 nm excitation laser and 530/30 emission filter (GFP), and 561 nm excitation laser and 620/15 emission filter (mCherry). Data were analyzed using Attune Cytometric Software v5.1.1 and FlowJo v10.7.1.

### FACS

Cells were sorted by fluorescence on a Sony Cell Sorter SH800Z (100 um sorting chip) equipped with 488 nm excitation laser and 525/50 emission filter (GFP), and 561 nm excitation laser and 600/60 emission filter (mCherry). Data were analyzed using Sony Cell Sorter Software v2.1.5.

### RT-qPCR

Total cell RNA was extracted using TRIzol Reagent (Thermo Fisher Scientific) as per manufacturer’s instructions. Genomic DNA was removed using TURBO DNase (Thermo Fisher Scientific). After inactivating TURBO DNase with DNase Inactivating Reagent, 1 ug DNase-free RNA was reverse transcribed using SuperScript III Reverse Transcriptase (Thermo Fisher Scientific) with random primers (Promega) as per manufacturer’s instructions. qPCR was performed using iTaq Universal SYBR Green Supermix (Bio-Rad) in a CFX96 Real-Time PCR Detection System (Bio-Rad). Gene-specific primer pairs are listed in Table S1.

### Cell viability and proliferation assay

The WST-1 assay was used to quantify cell viability and proliferation. Cells transfected with Csm, Cas13, or shRNA constructs were grown in 96-well plates until the indicated timepoints, incubated with WST-1 reagent (Sigma) at 37C for 1 hr as per manufacturer’s instructions, and absorbance measured using a Cytation 5 microplate reader (BioTek Instruments) at 450 nm with 600 nm reference.

### Microscopy

For wide-field fluorescent imaging, cells were observed on a Zeiss Axio Observer Z1 inverted fluorescence microscope, equipped with 63x/1.4 NA oil DIC and 100x/1.4 NA oil Ph3 Plan Apochromat objective lenses, ORCA-Flash4.0 camera (Hamamatsu), and ZEN 2012 software. Images were generated using ZEN 2012 (Zeiss) and FIJI (ImageJ) software. For live-cell imaging, cells were grown on chambered #1.5 coverglasses (Nunc Lab-Tek II) in medium lacking phenol red (Thermo Fisher Scientific) and imaged directly on the inverted fluorescent microscope.

### RNA FISH

Cells were grown on glass coverslips and rinsed in PBS. They were permeabilized in PBS/0.5% Triton X-100 for 10 min and then fixed in 4% paraformaldehyde for 10 min at room temp. Cells were dehydrated in a series of 70%, 80%, 90%, and 100% ethanol for 5 min each. Labeled oligo probe pool (10 nM final) was added to hybridization buffer containing 25% formamide, 2x SSC, 10% dextran sulfate, and nonspecific competitor (0.1 mg/mL human Cot-1 DNA [Thermo Fisher Scientific]). Hybridization was performed in a humidified chamber at 37C overnight. After being washed 1x in 25% formamide/2x SSC at 37C for 20 min and 3x in 2x SSC at 37C for 5 min each, cells were mounted for wide-field fluorescent imaging. Nuclei were counter-stained with Hoechst 33342 (Life Technologies).

### FISH probes

XIST oligo FISH probes were designed against the “Repeat D” region of human XIST RNA and synthesized by IDT carrying a 5’ Cy3 dye modification (see Table S1 for sequences). MALAT1 and NEAT1 oligo FISH probes were ordered from LGC Biosearch Technologies (SMF-2035-1, SMF-2036-1) carrying a Quasar 570 dye modification.

### Immunofluorescence

Cells were grown on glass coverslips and rinsed in PBS. They were fixed in 4% paraformaldehyde for 10 min and then permeabilized in PBS/0.5% Triton X-100 for 10 min at room temp. Cells were blocked with blocking buffer (PBS/0.05% Tween-20 containing 1% BSA) for 1 hr, incubated with primary antibody in blocking buffer for 1 hr, washed 3x with PBS/0.05% Tween-20 for 5 min each, incubated with dye-conjugated secondary antibody in blocking buffer for 1 hr at room temp, and washed 3x again with PBS/0.05% Tween-20 for 5 min each. Cells were mounted for wide-field fluorescent imaging and nuclei were counter-stained with Hoechst 33342 (Life Technologies).

### Western blot

Cells were washed once with PBS and lysed in cold RIPA lysis buffer (50 mM Tris pH 7.5, 150 mM NaCl, 1% NP-40, 1% sodium deoxycholate, 0.1% SDS, 1x protease inhibitor cocktail [Sigma]). Lysate was sonicated (Qsonica Q800 Sonicator) in polystyrene tubes at 50% power setting, 30 sec on/30 sec off for a total sonication time of 5 min at 4C. After removing debris by centrifugation at 16,000 g for 10 min, protein concentration in the supernatant was measured (Pierce BCA Assay Kit). 20-50 ug protein lysate was denatured in 1x Laemmli buffer at 95C for 10 min and resolved by SDS-PAGE. Protein was transferred to Immun-Blot LF PVDF membrane (Bio-Rad). The membrane was blocked with blocking buffer (PBS/0.05% Tween-20 containing 5% milk) for 1 hr at room temp, incubated with primary antibody in blocking buffer overnight at 4C, washed 3x with PBS/0.05% Tween-20 for 5 min each, incubated with dye-conjugated secondary antibody in blocking buffer for 1 hr at room temp, and washed 3x again with PBS/0.05% Tween-20 for 5 min each. Protein bands were visualized on a LI-COR Odyssey CLx with Image Studio v5.2 software using 700 nm and 800 nm channels.

### Antibodies

The following primary antibodies were used for Western blot: mouse anti-FLAG (Sigma, F1804), rabbit anti-GAPDH (Cell Signaling Technology, 14C10); for immunofluorescence: mouse anti-FLAG (Sigma, F1804). The following secondary antibodies were used for Western blot: IRDye 680RD goat anti-mouse (LI-COR, 926-68070), IRDye 800CW goat anti-rabbit (LI-COR, 926-32211); for immunofluorescence: Alexa Fluor 555 goat anti-mouse (Invitrogen, A21424).

### RNA-seq

Total cell RNA was extracted using TRIzol Reagent (Thermo Fisher Scientific). Strand-specific cDNA libraries were prepared from polyA mRNA and sequenced using the Illumina NovaSeq paired-end 150 bp platform by Novogene. Libraries were sequenced to a depth of 30 million reads each, with 3 biological replicates per sample.

### RNA-seq analysis

Custom scripts were used for transcriptomic analysis. Briefly, reads were assessed for sequencing quality with FastQC, then adapters and low-quality bases were trimmed with CutAdapt. Samples were aligned to the GRCh38 reference genome (GENCODE Release 39) with STAR and uniquely mapped reads were used to generate a count matrix with FeatureCounts. EdgeR was used to normalize read counts and identify differentially expressed genes. Genes with a fold-change ≥2 relative to the untransfected sample were considered differentially expressed. Off-target editing was interrogated through sequence-similarity, by aligning crRNA sequences to the human transcriptome (GRCh38 cDNA, ENSEMBL release 105) with blastn and lenient parameters (E-value = 10000, word_size = 5, perc_identity 0.6). Potential off-targets were limited to BLAST results with 7 or fewer mismatched bases.

### Statistical analysis

All graphs display the mean and standard deviation of 3 biological replicates.

### Data and materials availability

RNA-seq datasets have been deposited at GEO (GSE###). Essential plasmids have been deposited at Addgene (plasmid ###). Original data and unprocessed images have been deposited at ###.

## ACKNOWLEDGMENTS

We thank all members of the Doudna lab for helpful advice and discussion. This project was supported by XXX (grant no. ###). D.C. is supported by the Jane Coffin Childs Memorial Fund for Medical Research. J.A.D. is an investigator of the Howard Hughes Medical Institute.

## Author contributions

D.C. and J.A.D. conceived of the project. J.A.D. supervised the project. D.C. designed and performed all experiments. M.T. performed all bioinformatics analyses. D.C., M.T., and J.A.D. wrote the manuscript.

