## Supplement for "Precise Transcript Targeting by CRISPR-Csm Complexes"

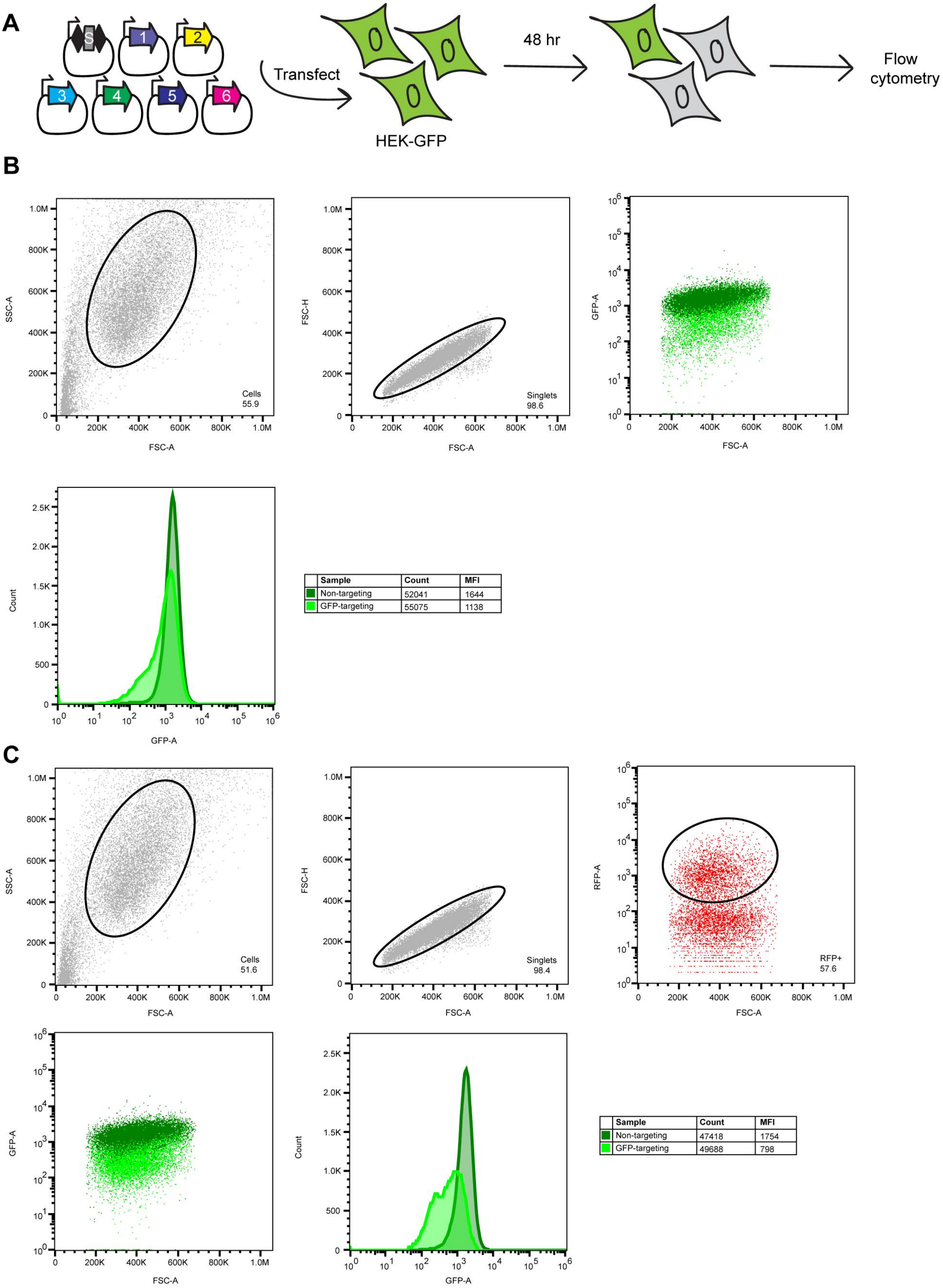
A
S
1
2
3
4
5
6
Transfect
HEK-GFP
48 hr
Flow cytometry
B
SSC-A
FSC-A
Cells
55.9
FSC-H
Singlets
98.6
GFP-A
Count
Sample | Count | MFI
Non-targeting | 52041 | 1644
GFP-targeting | 55075 | 1138
C
Cells
51.6
Singlets
98.4
RFP-A
RFP+
57.6
Sample | Count | MFI
Non-targeting | 47418 | 1754
GFP-targeting | 49688 | 798

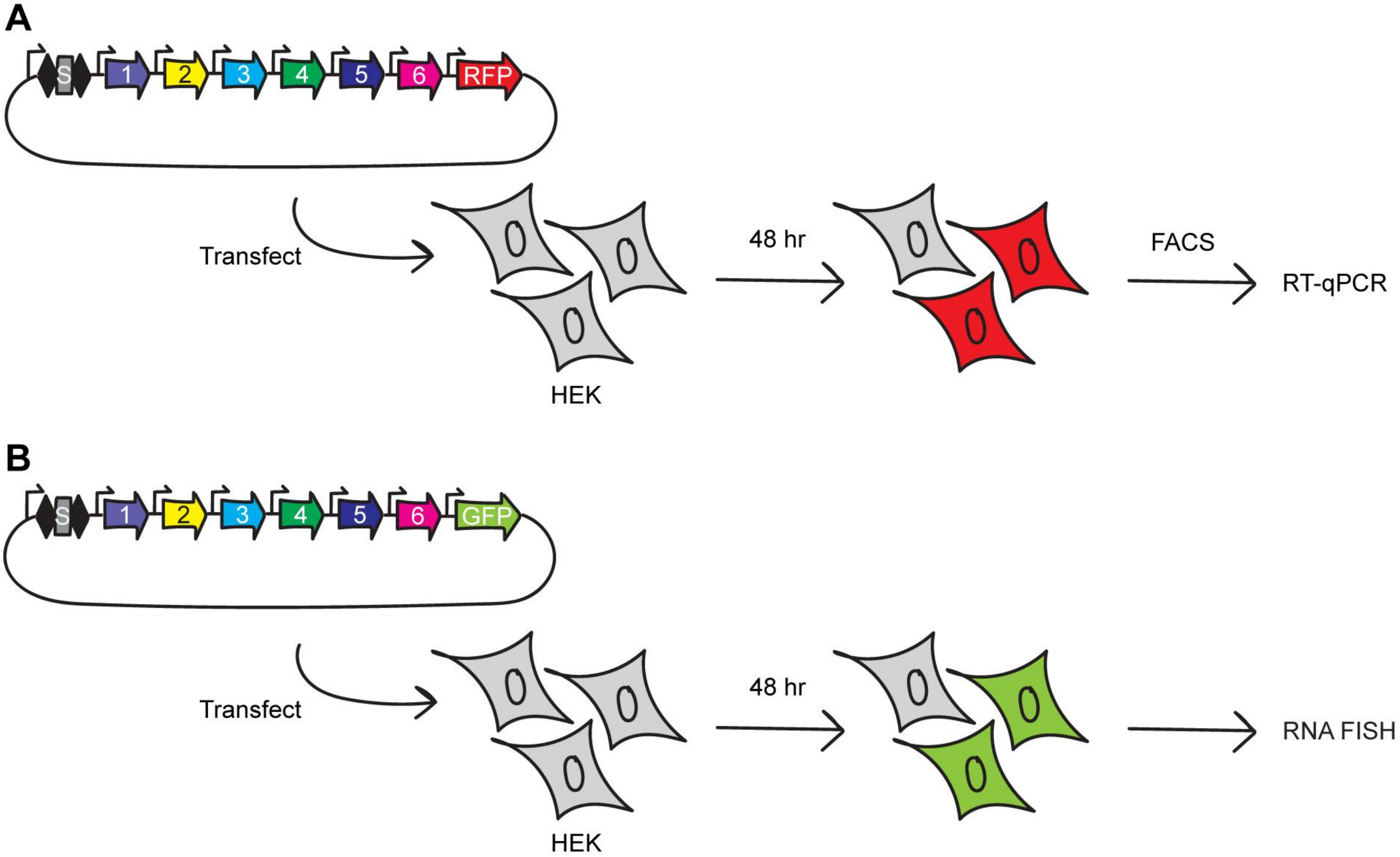
A
S
1
2
3
4
5
6
RFP
Transfect
HEK
48 hr
FACS
RT-qPCR
B
S
1
2
3
4
5
6
GFP
Transfect
HEK
48 hr
RNA FISH

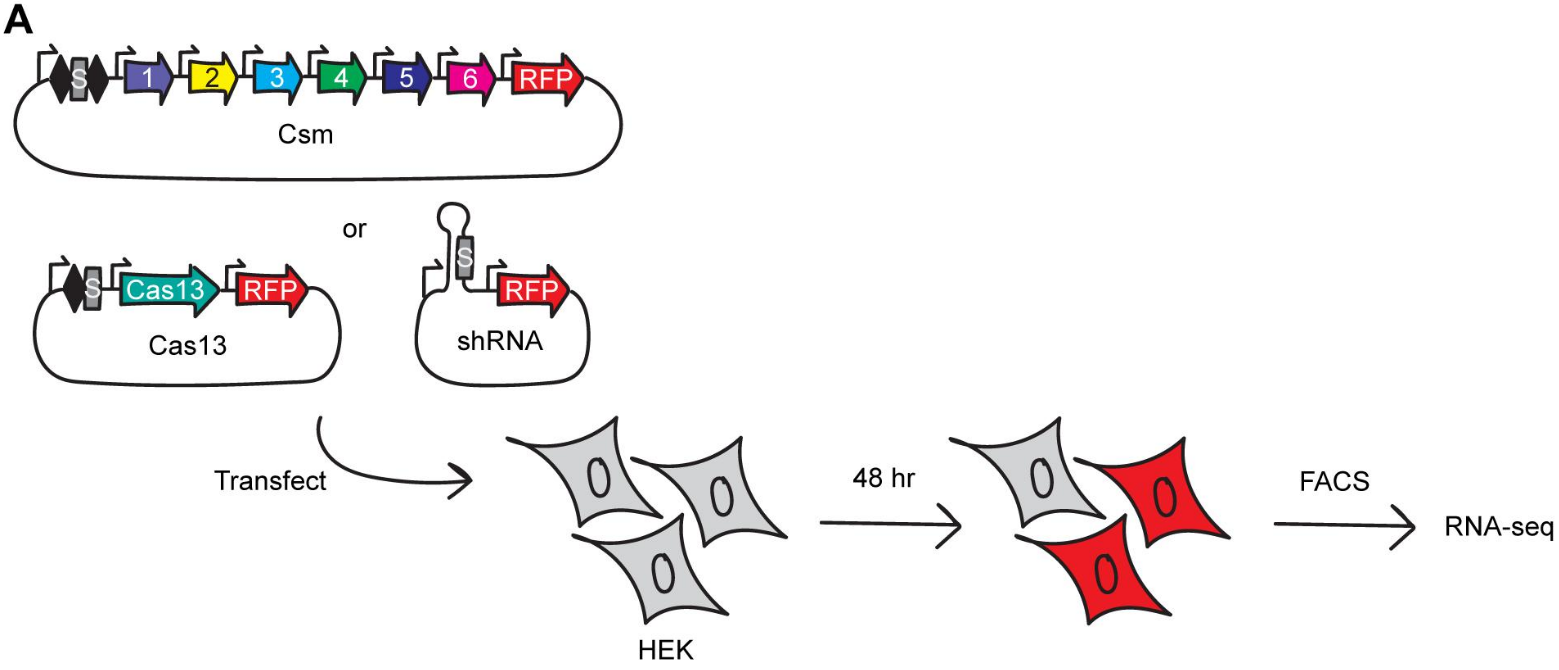
A
S
1
2
3
4
5
6
RFP
Csm
or
S
Cas13
RFP
Cas13
S
RFP
shRNA
Transfect
HEK
48 hr
FACS
RNA-seq

**A**

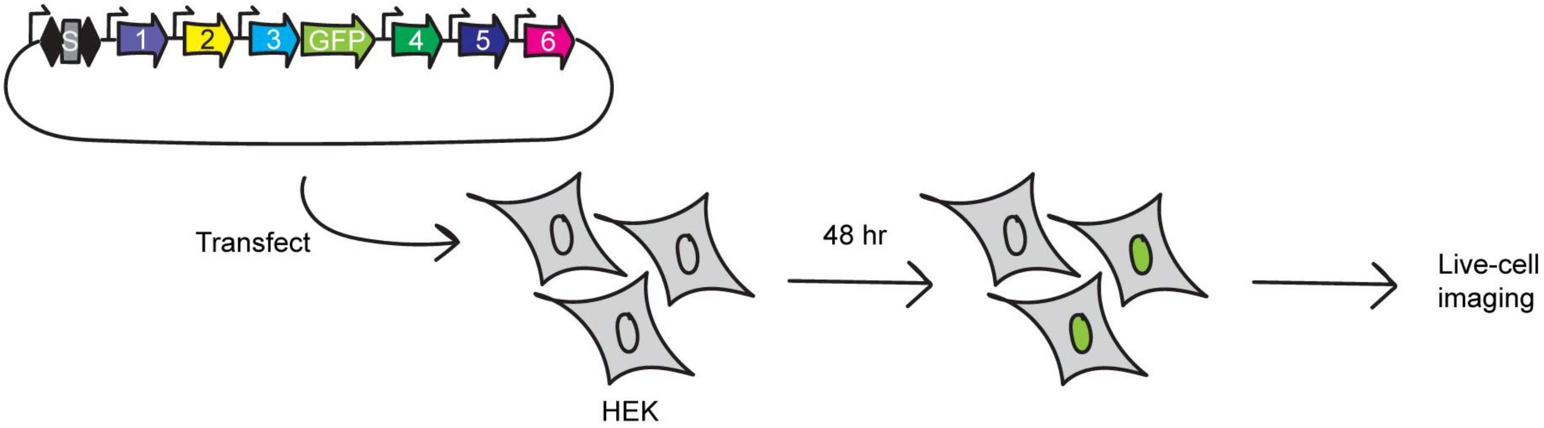

Table S1

| Target | qPCR primer F | qPCR primer R |
|---|---|---|
| **XIST** | GTTGTATCGGGAGGCAGTAAGAATCATCTTT | GAAAAGCACACAGCAAAGACAAAGAGGC |
| **MALAT1** | ACTAGCATTAATTGACAGCTGACCCAGG | GCTACCTTCATCACCAAATTGCACTCG |
| **NEAT1** | GCTTAGGAGGAGGAAGTTCTCCAATGT | CTCCATCTGCAAGCTCCATCTACAAG |
| **BRCA1** | TACATCAGGCCTTCATCCTGAGGATTTTATC | ACAATTAGGTGGGCTTAGATTTCTACTGACTACTA |
| **TARDBP** | GTCAAGAAAGATCTTAAGACTGGTCATTCAAAGGG | CTTAGAATTAGGAAGTTTGCAGTCACACCATC |
| **SMARCA1** | GATAAACCAGTCAAATCTAAACTGGGGAGCA | GATACAAGGCTCCATTTCATCAGTTGCG |
| **CKB** | TTAAGCACCTCCGAGAACTTCTCATGC | TTGAAACTCTCTTCAAGTCTAAGGACTATGAGTTCA |
| **ENO1** | GGAACTCATTAATATACTTAATGGGTCTGGAGACG | CTTCAACTGGTATCTATGAGGCCCTAGAG |
| **MECP2** | GGAAGAAAAGTCAGAAGACCAGGACC | TTGATCAAATACACATCATACTTCCCAGCAGAG |
| **UBE3A** | ATATTGATGCCATTAGAAGGGTCTACACCAGAT | CTTTGCAAAATAATGGCAAAGCCATTTCCAG |
| **SMAD4** | ATGGACAATATGTCTATTACGAATACACCAACAAG TAATG | CTGAAGCCTCCCATCCAATGTTCTCT |
| **GAPDH** | CCAGAACATCATCCCTGCCTCTACTG | GGAAATGAGCTTGACAAAGTGGTCGTTG |

| Target | Csm crRNA 1 | Csm crRNA 2 | Csm crRNA 3 |
|---|---|---|---|
| **XIST** | GCCACTTGAACACTGCGACAGA ACTGGATCCG | TTGGACAACCTAACAAAGCACAGC CCGCCATG | CGCACATGTCCACCACCATGCTAA CCACTTAA |
| **MALAT1** | AGCTTCCTTCACCAAATCGCAC TGGCTCCTGG | GCCGCCTGCTACCTTCATCACCAA ATTGCACT | CCTAGCTTCACCACCAAATCGTTA GCGCTCCT |
| **NEAT1** | CCGGATGCATCTGCTGTGGACT TTTTAAGATT | CACCATTACCAACAATACCGACTC CAACAGCC | GAAGATGCAGCATCTGAAAACCTT TACCCCAG |
| **BRCA1** | GGTTAGGATTTTTCTCATTCTG AATAGAATCA | ATTGTGGATATTTAATTCGAGTTC CATATTGC | GAGCAGAGGGTGAAGGCCTCCTGA GCGCAGGG |
| **TARDBP** | CTTGTGTTTCATATTCCGTAAA ACGAACAAAG | GTCCATCTATCATATGTCGCTGTG ACATTACT | CTTTGAATGACCAGTCTTAAGATC TTTCTTGA |
| **SMARCA1** | GTCCATCCAGTCGACAATACTC ATAACCACGC | CCGACAAAACAAATGACACGGAGA GATGGGAC | AAAAGTTGAGTAAGGCCCACAGTT CATGCAGG |
| **CKB** | GCTTGTCGAAGAGGAAGTGGTC GTCGATGAGC | TTGATATGCACACCTGCCCGCAGC CCGGTGCC | CGAGAACTTCTCATGCTTGCCCAG GTTGGGCA |
| **ENO1** | CATCCATCTCGATCATCAGTTT GTCAATCTTC | TTCCCGCGAGAGTCAAAGATCTCC CTGGCATG | GACCCCCTTCTCAACGGCACCAGC TTTGCAGA |
| **MECP2** | CAGAGTGGTGGGCTGATGGCTG CACGGGCTCA | GGCAGAAGCTTCCGGCACAGCCGG GGCGGAGC | CAAATACACATCATACTTCCCAGC AGAGCGGC |
| **UBE3A** | CTCGAGAGTATACATTGTGATA CGTCAAGTCA | GAGATTTCTATTCTCCATTACGAT AATGAACA | AAATTCCACATACAACTGCTTCTT CAAGTCTG |
| **SMAD4** | TCCACCTTGTCTATGGCACATC AAACTATGCA | AGCTTCTTTACCAAACTTTCAATT GCTCTTTT | TCCAATGTTCTCTGTATGGTAACA CATTTACT |
| **GFP** | CAGCTTGCCGGTGGTGCAGATG AACTTCAGGG | TGAAGCACTGCACGCCGTAGGTCA GGGTGGTC | AGGATGTTGCCGTCCTCCTTGAAG TCGATGCC |
| **RFP** | CTTGAAGCCCTCGGGGAAGGAC AGCTTCAAGT | | |
| **NT** | TCTCCGAACGTGTCACGTCTTT AGCGACTAAA | | |

| Target | Cas13 crRNA 1 | Cas13 crRNA 2 |
|---|---|---|
| **XIST** | GCCACTTGAACACTGCGACAGAACTGGATC | TTGGACAACCTAACAAAGCACAGCCCGCCA |
| **MALAT1** | AGCTTCCTTCACCAAATCGCACTGGCTCCT | GCCGCCTGCTACCTTCATCACCAAATTGCA |
| **CKB** | GCTTGTCGAAGAGGAAGTGGTCGTCGATGA | TTGATATGCACACCTGCCCGCAGCCCGGTG |
| **SMAD4** | TCCACCTTGTCTATGGCACATCAAACTATG | AGCTTCTTTACCAAACTTTCAATTGCTCTT |
| **NT** | TCTCCGAACGTGTCACGTCTTTAGCGACTA | |

| Target | shRNA 1 | shRNA 2 |
|---|---|---|
| **XIST** | GTTCTGTCGCAGTGTTCAAGTcctgacccaACTT GAACACTGCGACAGAAC | GTGCTTTGTTAGGTTGTCCAAcctgacccaTTGGACAA CCTAACAAAGCAC |
| **MALAT1** | GTGCGATTTGGTGAAGGAAGCcctgacccaGCTT CCTTCACCAAATCGCAC | GTGATGAAGGTAGCAGGCGGCcctgacccaGCCGCCTG CTACCTTCATCAC |
| **CKB** | GACCACTTCCTCTTCGACAAGcctgacccaCTTG TCGAAGAGGAAGTGGTC | GCGGGCAGGTGTGCATATCAAcctgacccaTTGATATG CACACCTGCCCGC |
| **SMAD4** | GATGTGCCATAGACAAGGTGGcctgacccaCCAC CTTGTCTATGGCACATC | GCAATTGAAAGTTTGGTAAAGcctgacccaCTTTACCA AACTTTCAATTGC |

Table S1

| | |
|---|---|
| **NT** | GCTAAAGACGTGACACGTTCGcctgacccaCGAA CGTGTCACGTCTTTAGC |

| **spacer length** | **GFP** |
|---|---|
| **24 nt** | CAGCTTGCCGGTGGTGCAGATGAA |
| **28 nt** | CAGCTTGCCGGTGGTGCAGATGAACTTC |
| **32 nt** | CAGCTTGCCGGTGGTGCAGATGAACTTCAGGG |
| **36 nt** | CAGCTTGCCGGTGGTGCAGATGAACTTCAGGGTCAG |
| **40 nt** | CAGCTTGCCGGTGGTGCAGATGAACTTCAGGGTCAGCTTG |
| **44 nt** | CAGCTTGCCGGTGGTGCAGATGAACTTCAGGGTCAGCTTGCCGT |
| **48 nt** | CAGCTTGCCGGTGGTGCAGATGAACTTCAGGGTCAGCTTGCCGTAGGT |

| **Target** | **Live-cell imaging Csm crRNA** |
|---|---|
| **XIST** | AAAAGCAGGTATCCGCGGCCCCGATGGGCAAA |
| **MALAT1** | AGCTTCCTTCACCAAATCGCACTGGCTCCTGG |
| **NEAT1** | CCGGATGCATCTGCTGTGGACTTTTTAAGATT |

| **Target** | **RNA FISH probe** |
|---|---|
| **XIST** | /5Cy3/GGGCACTCCCTGCTGGAAGGGAA |
| | /5Cy3/AATTGTGCACCTTGACTGTCCAAA |
| | /5Cy3/TCTGAGAGTAGGACCTTATTCA |
| | /5Cy3/TCAGCACCCCTGCTGTACTGCAAA |

Table S2

| | |
|---|---|
| **Plasmid** | **pDAC338** |
| **Description** | Expression of Csm1 |
| **Utility** | |
| **Features** | Pcmv-FLAG-NLS-Csm1-pA |
| **Sequence** | ACATGTGAGCAAAAGGCCAGCAAAAGGCCAGGAACCGTAAAAAGGCCGCGTTGCTGGCGTTTTTCCATAGG<br>CTCCGCCCCCCTGACGAGCATCACAAAAATCGACGCTCAAGTCAGAGGTGGCGAAACCCGACAGGACTATA<br>AAGATACCAGGCGTTTCCCCCTGGAAGCTCCCTCGTGCGCTCTCCTGTTCCGACCCTGCCGCTTACCGGAT<br>ACCTGTCCGCCTTTCTCCCTTCGGGAAGCGTGGCGCTTTCTCATAGCTCACGCTGTAGGTATCTCAGTTCG<br>GTGTAGGTCGTTCGCTCCAAGCTGGGCTGTGTGCACGAACCCCCCGTTCAGCCCGACCGCTGCGCCTTATC<br>CGGTAACTATCGTCTTGAGTCCAACCCGGTAAGACACGACTTATCGCCACTGGCAGCAGCCACTGGTAACA<br>GGATTAGCAGAGCGAGGTATGTAGGCGGTGCTACAGAGTTCTTGAAGTGGTGGCCTAACTACGGCTACACT<br>AGAAGAACAGTATTTGGTATCTGCGCTCTGCTGAAGCCAGTTACCTTCGGAAAAAGAGTTGGTAGCTCTTG<br>ATCCGGCAAACAAACCACCGCTGGTAGCGGTGGTTTTTTTGTTTGCAAGCAGCAGATTACGCGCAGAAAAA<br>AAGGATCTCAAGAAGATCCTTTGATCTTTTCTACGGGGTCTGACGCTCAGTGGAACGAAAACTCACGTTAA<br>GGGATTTTGGTCATGAGATTATCAAAAAGGATCTTCACCTAGATCCTTTTAAATTAAAAATGAAGTTTTAA<br>ATCAATCTAAAGTATATATGAGTAAACTTGGTCTGACAGTTACCAATGCTTAATCAGTGAGGCACCTATCT<br>CAGCGATCTGTCTATTTCGTTCATCCATAGTTGCCTGACTCCCCGTCGTGTAGATAACTACGATACGGGAG<br>GGCTTACCATCTGGCCCCAGTGCTGCAATGATACCGCGAGACCCACGCTCACCGGCTCCAGATTTATCAGC<br>AATAAACCAGCCAGCCGGAAGGGCCGAGCGCAGAAGTGGTCCTGCAACTTTATCCGCCTCCATCCAGTCTA<br>TTAATTGTTGCCGGGAAGCTAGAGTAAGTAGTTCGCCAGTTAATAGTTTGCGCAACGTTGTTGCCATTGCT<br>ACAGGCATCGTGGTGTCACGCTCGTCGTTTGGTATGGCTTCATTCAGCTCCGGTTCCCAACGATCAAGGCG<br>AGTTACATGATCCCCCATGTTGTGCAAAAAAGCGGTTAGCTCCTTCGGTCCTCCGATCGTTGTCAGAAGTA<br>AGTTGGCCGCAGTGTTATCACTCATGGTTATGGCAGCACTGCATAATTCTCTTACTGTCATGCCATCCGTA<br>AGATGCTTTTCTGTGACTGGTGAGTACTCAACCAAGTCATTCTGAGAATAGTGTATGCGGCGACCGAGTTG<br>CTCTTGCCCGGCGTCAATACGGGATAATACCGCGCCACATAGCAGAACTTTAAAAGTGCTCATCATTGGAA<br>AACGTTCTTCGGGGCGAAAACTCTCAAGGATCTTACCGCTGTTGAGATCCAGTTCGATGTAACCCACTCGT<br>GCACCCAACTGATCTTCAGCATCTTTTACTTTCACCAGCGTTTCTGGGTGAGCAAAAACAGGAAGGCAAAA<br>TGCCGCAAAAAAGGGAATAAGGGCGACACGGAAATGTTGAATACTCATACTCTTCCTTTTTCAATATTATT<br>GAAGCATTTATCAGGGTTATTGTCTCATGAGCGGATACATATTTGAATGTATTTAGAAAAATAAACAAATA<br>GGGGTTCCGCGCACATTTCCCCGAAAAGTGCCACCTGACGTCGGATCCgacattgattattgactagttat<br>taatagtaatcaattacggggtcattagttcatagcccatatatggagttccgCGTTACATAACTTACGGT<br>AAATGGCCCGCCTGGCTGACCGCCCAACGACCCCCGCCCATTGACGTCAATAATGACGTATGTTCCCATAG<br>TAACGCCAATAGGGACTTTCCATTGACGTCAATGGGTGGAGTATTTACGGTAAACTGCCCACTTGGCAGTA<br>CATCAAGTGTATCATATGCCAAGTACGCCCCCTATTGACGTCAATGACGGTAAATGGCCCGCCTGGCATTA<br>TGCCCAGTACATGACCTTATGGGACTTTCCTACTTGGCAGTACATCTACGTATTAGTCATCGCTATTACCA<br>TGGTGATGCGGTTTTGGCAGTACATCAATGGGCGTGGATAGCGGTTTGACTCACGGGGATTTCCAAGTCTC<br>CACCCCATTGACGTCAATGGGAGTTTGTTTTGGCACCAAAATCAACGGGACTTTCCAAAATGTCGTAACAA<br>CTCCGCCCCATTGACGCAAATGGGCGGTAGGCGTGTACGGTGGGAGGTCTATATAAGCAGAGCTCGTTTAG<br>TGAACCGTCAGATCTCTAGAgccgccaccATGGATTACAAAGACGATGACGATAAGATGGCGCCTAAGAAG<br>AAACGCAAAGTGCGGGCATGAAGAAAGAAAAGATTGATCTGTTTTACGGAGCCCTGCTGCACGACATCGG<br>AAAGGTCATCCAGCGAGCAACCGGAGAGCGGAAGAAACACGCACTTGTGGGCGCCGACTGGTTCGACGAGA<br>TCGCCGACAACCAAGTCATCTCGGATCAGATCCGGTACCATATGGCCAACTACCAGTCTGATAAGCTCGGC<br>AACGATCACCTGGCTTACATCACCTACATTGCCGACAACATCGCCTCCGGTGTCGACCGCCGGCAATCCAA<br>CGAAGAGTCAGACGAAGATACCTCCGCAAAGATCTGGGACACCTACACGAACCAGGCCGACATCTTTAACG<br>TGTTCGGAGCGCAGACCGATAAGCGGTACTTCAAGCCTACCGTGCTGAATCTCAAGTCGAAGCCCAACTTC<br>GCGTCCGCCACTTACGAACCCTTTAGCAAGGGCGATTACGCTGCCATCGCCACCCGGATTAAGAACGAACT<br>GGCCGAGTTCGAGTTCAACCAAGTCCAGATTGACTCCCTGCTCAACCTTTTCGAGGCTACTCTCTCCTTCG<br>TGCCGTCAAGCACCAACACTAAGGAAATCGCCGACATCTCCCTGGCCGACCATTCCCGCTTGACTGCTGCC<br>TTCGCTCTGGCGATCTACGACTACCTGGAGGACAAGGGTCGGCACAACTACAAAGAGGACCTGTTCACCAA<br>AGTGTCAGCGTTCTATGAAGAAGAAGCCTTCCTGCTGGCCTCCTTCGACCTGTCGGGAATCCAGGACTTTA<br>TCTACAACATTAACATCGCAACTAACGGCGCGGCGAAGCAGCTGAAGGCCCGGAGCCTCTACCTGGACTTT<br>ATGTCCGAGTACATCGCCGATAGCCTGCTGGACAAGCTGGGACTGAACAGGGCTAACATGCTTTACGTCGG<br>CGGCGGACACGCCTACTTCGTCCTGGCCAACACCGAAAAGACTGTGGAAACCCTGGTGCAGTTTGAGAAGG<br>ATTTCAACCAGTTCCTGTTGGCAAACTTCCAGACCCGCCTCTATGTGGCCTTTGGCTGGGGTTCCTTCGCG<br>GCCAAGGACATCATGTCCGAGCTGAATAGCCCCGAGTCCTACCGCCAAGTGTACCAAAAGGCTTCGCGCAT<br>GATCTCCAAAAAGAAAATCTCCAGATACGACTACCAGACACTGATGCTCCTGAATCGCGGTGGAAAGTCCT<br>CAGAGAGAGAGTGCGAGATTTGCCACTCCGTGGAGAACCTGGTGTCCTACCACGACCAGAAAGTCTGTGAC<br>ATTTGCCGGGGACTGTACCAGTTCTCGAAAGAAATTGCCCATGACCACTTCATCATTACCGAAAATGAGGG<br>GCTGCCGATTGGACCAAACGCGTGCTTAAAGGGCGTGGCATTCGAAAAGCTGTCCCAAGAAGCGTTCAGCC<br>GGGTCTACGTGAAGAATGACTATAAGGCCGGTACCGTGAAGGCTACGCATGTGTTCGTGGGGGATTACCAG<br>TGCGACGAGATCTACAACTACGCCGCCCTGAGCAAGAACGAGAACGGCCTAGGCATCAAGAGACTGGCCGT<br>GGTCCGGCTCGACGTGGATGACTTGGGCGCCGCCTTCATGGCCGGTTTCAGCCAGCAGGGAAACGGACAAT<br>ACTCCACTCTGTCAAGATCGGCCACATTCTCCCGGAGCATGTCGCTGTTCTTCAAAGTGTACATTAACCAG |

Table S2

| | |
|---|---|
| | TTCGCCTCCGACAAGAAGCTGAGCATTATCTACGCGGGCGGCGATGACGTGTTCGCCATTGGATCGTGGCA<br>GGATATCATCGCGTTCACTGTGGAACTTCGCGAAAACTTCATCAAGTGGACCAACGGGAAGCTCACCCTCT<br>CCGCGGGGATAGGGTTGTTCGCCGACAAGACTCCTATTAGCCTGATGGCTCACCAGACCGGGGAACTGGAA<br>GAGGCCGCCAAGGGCAACGAAAAGGACTCCATCTCGCTGTTCTCAAGCGACTACACTTTCAAGTTTGATAG<br>GTTCATCACTAACGTGTACGACGACAAACTGGAACAGATTAGATACTTCTTCAACCATCAAGACGAGAGGG<br>GAAAGAACTTCATCTATAAGCTTATTGAGCTTTTGAGGAACCACGACCGCATGAATATGGCACGCCTCGCC<br>TATTACCTCACTCGCCTGGAAGAACTGACCCGGGAGACTGACAGGGACAAGTTCAAGACCTTCAAGAACCT<br>GTTCTACTCCTGGTACACCAACAAGAACGATAAGGACCGGAAGGAAGCCGAGCTCGCGCTCCTGCTGTACA<br>TCTACGAAATCAGAAAGGATTAAcggcaataaaaagacagaataaaacgcacggtgttgggtcgtttgttc<br>AAGCTC |

| | |
|---|---|
| **Plasmid** | **pDAC758** |
| **Description** | Expression of Csm1(DNase/cA mut) |
| **Utility** | |
| **Features** | Pcmv-FLAG-NLS-Csm1(DNase/cA mut)-pA |
| **Sequence** | ACATGTGAGCAAAAGGCCAGCAAAAGGCCAGGAACCGTAAAAAGGCCGCGTTGCTGGCGTTTTTCCATAGG<br>CTCCGCCCCCCTGACGAGCATCACAAAAATCGACGCTCAAGTCAGAGGTGGCGAAACCCGACAGGACTATA<br>AAGATACCAGGCGTTTCCCCCTGGAAGCTCCCTCGTGCGCTCTCCTGTTCCGACCCTGCCGCTTACCGGAT<br>ACCTGTCCGCCTTTCTCCCTTCGGGAAGCGTGGCGCTTTCTCATAGCTCACGCTGTAGGTATCTCAGTTCG<br>GTGTAGGTCGTTCGCTCCAAGCTGGGCTGTGTGCACGAACCCCCCGTTCAGCCCGACCGCTGCGCCTTATC<br>CGGTAACTATCGTCTTGAGTCCAACCCGGTAAGACACGACTTATCGCCACTGGCAGCAGCCACTGGTAACA<br>GGATTAGCAGAGCGAGGTATGTAGGCGGTGCTACAGAGTTCTTGAAGTGGTGGCCTAACTACGGCTACACT<br>AGAAGAACAGTATTTGGTATCTGCGCTCTGCTGAAGCCAGTTACCTTCGGAAAAAGAGTTGGTAGCTCTTG<br>ATCCGGCAAACAAACCACCGCTGGTAGCGGTGGTTTTTTTGTTTGCAAGCAGCAGATTACGCGCAGAAAAA<br>AAGGATCTCAAGAAGATCCTTTGATCTTTTCTACGGGGTCTGACGCTCAGTGGAACGAAAACTCACGTTAA<br>GGGATTTTGGTCATGAGATTATCAAAAAGGATCTTCACCTAGATCCTTTTAAATTAAAAATGAAGTTTTAA<br>ATCAATCTAAAGTATATATGAGTAAACTTGGTCTGACAGTTACCAATGCTTAATCAGTGAGGCACCTATCT<br>CAGCGATCTGTCTATTTCGTTCATCCATAGTTGCCTGACTCCCCGTCGTGTAGATAACTACGATACGGGAG<br>GGCTTACCATCTGGCCCCAGTGCTGCAATGATACCGCGAGACCCACGCTCACCGGCTCCAGATTTATCAGC<br>AATAAACCAGCCAGCCGGAAGGGCCGAGCGCAGAAGTGGTCCTGCAACTTTATCCGCCTCCATCCAGTCTA<br>TTAATTGTTGCCGGGAAGCTAGAGTAAGTAGTTCGCCAGTTAATAGTTTGCGCAACGTTGTTGCCATTGCT<br>ACAGGCATCGTGGTGTCACGCTCGTCGTTTGGTATGGCTTCATTCAGCTCCGGTTCCCAACGATCAAGGCG<br>AGTTACATGATCCCCCATGTTGTGCAAAAAAGCGGTTAGCTCCTTCGGTCCTCCGATCGTTGTCAGAAGTA<br>AGTTGGCCGCAGTGTTATCACTCATGGTTATGGCAGCACTGCATAATTCTCTTACTGTCATGCCATCCGTA<br>AGATGCTTTTCTGTGACTGGTGAGTACTCAACCAAGTCATTCTGAGAATAGTGTATGCGGCGACCGAGTTG<br>CTCTTGCCCGGCGTCAATACGGGATAATACCGCGCCACATAGCAGAACTTTAAAAGTGCTCATCATTGGAA<br>AACGTTCTTCGGGGCGAAAACTCTCAAGGATCTTACCGCTGTTGAGATCCAGTTCGATGTAACCCACTCGT<br>GCACCCAACTGATCTTCAGCATCTTTTACTTTCACCAGCGTTTCTGGGTGAGCAAAAACAGGAAGGCAAAA<br>TGCCGCAAAAAAGGGAATAAGGGCGACACGGAAATGTTGAATACTCATACTCTTCCTTTTTCAATATTATT<br>GAAGCATTTATCAGGGTTATTGTCTCATGAGCGGATACATATTTGAATGTATTTAGAAAAATAAACAAATA<br>GGGGTTCCGCGCACATTTCCCCGAAAAGTGCCACCTGACGTCGGATCCgacattgattattgactagttat<br>taatagtaatcaattacggggtcattagttcatagcccatatatggagttccgCGTTACATAACTTACGGT<br>AAATGGCCCGCCTGGCTGACCGCCCAACGACCCCCGCCCATTGACGTCAATAATGACGTATGTTCCCATAG<br>TAACGCCAATAGGGACTTTCCATTGACGTCAATGGGTGGAGTATTTACGGTAAACTGCCCACTTGGCAGTA<br>CATCAAGTGTATCATATGCCAAGTACGCCCCCTATTGACGTCAATGACGGTAAATGGCCCGCCTGGCATTA<br>TGCCCAGTACATGACCTTATGGGACTTTCCTACTTGGCAGTACATCTACGTATTAGTCATCGCTATTACCA<br>TGGTGATGCGGTTTTGGCAGTACATCAATGGGCGTGGATAGCGGTTTGACTCACGGGGATTTCCAAGTCTC<br>CACCCCATTGACGTCAATGGGAGTTTGTTTTGGCACCAAAATCAACGGGACTTTCCAAAATGTCGTAACAA<br>CTCCGCCCCATTGACGCAAATGGGCGGTAGGCGTGTACGGTGGGAGGTCTATATAAGCAGAGCTCGTTTAG<br>TGAACCGTCAGATCTCTAGAgccgccaccATGGATTACAAAGACGATGACGATAAGATGGCGCCTAAGAAG<br>AAACGCAAAGTGCGGGGCATGAAGAAAGAAAAGATTGATCTGTTTTACGGAGCCCTGCTGGCCGCCATCGG<br>AAAGGTCATCCAGCGAGCAACCGGAGAGCGGAAGAAACACGCACTTGTGGGCGCCGACTGGTTCGACGAGA<br>TCGCCGACAACCAAGTCATCTCGGATCAGATCCGGTACCATATGGCCAACTACCAGTCTGATAAGCTCGGC<br>AACGATCACCTGGCTTACATCACCTACATTGCCGACAACATCGCCTCCGGTGTCGACCGCCGGCAATCCAA<br>CGAAGAGTCAGACGAAGATACCTCCGCAAAGATCTGGGACACCTACACGAACCAGGCCGACATCTTTAACG<br>TGTTCGGAGCGCAGACCGATAAGCGGTACTTCAAGCCTACCGTGCTGAATCTCAAGTCGAAGCCCAACTTC<br>GCGTCCGCCACTTACGAACCCTTTAGCAAGGGCGATTACGCTGCCATCGCCACCCGGATTAAGAACGAACT<br>GGCCGAGTTCGAGTTCAACCAAGTCCAGATTGACTCCCTGCTCAACCTTTTCGAGGCTACTCTCTCCTTCG<br>TGCCGTCAAGCACCAACACTAAGGAAATCGCCGACATCTCCCTGGCCGACCATTCCCGCTTGACTGCTGCC<br>TTCGCTCTGGCGATCTACGACTACCTGGAGGACAAGGGTCGGCACAACTACAAAGAGGACCTGTTCACCAA<br>AGTGTCAGCGTTCTATGAAGAAGAAGCCTTCCTGCTGGCCTCCTTCGACCTGTCGGGAATCCAGGACTTTA<br>TCTACAACATTAACATCGCAACTAACGGCGCGGCGAAGCAGCTGAAGGCCCGGAGCCTCTACCTGGACTTT<br>ATGTCCGAGTACATCGCCGATAGCCTGCTGGACAAGCTGGGACTGAACAGGGCTAACATGCTTTACGTCGG |

Table S2

| | |
|---|---|
| | CGGCGGACACGCCTACTTCGTCCTGGCCAACACCGAAAAGACTGTGGAAACCCTGGTGCAGTTTGAGAAGG<br>ATTTCAACCAGTTCCTGTTGGCAAACTTCCAGACCCGCCTCTATGTGGCCTTTGGCTGGGGTTCCTTCGCG<br>GCCAAGGACATCATGTCCGAGCTGAATAGCCCCGAGTCCTACCGCCAAGTGTACCAAAAGGCTTCGCGCAT<br>GATCTCCAAAAAGAAAATCTCCAGATACGACTACCAGACACTGATGCTCCTGAATCGCGGTGGAAAGTCCT<br>CAGAGAGAGAGTGCGAGATTTGCCACTCCGTGGAGAACCTGGTGTCCTACCACGACCAGAAAGTCTGTGAC<br>ATTTGCCGGGGACTGTACCAGTTCTCGAAAGAAATTGCCCATGACCACTTCATCATTACCGAAAATGAGGG<br>GCTGCCGATTGGACCAAACGCGTGCTTAAAGGGCGTGGCATTCGAAAAGCTGTCCCAAGAAGCGTTCAGCC<br>GGGTCTACGTGAAGAATGACTATAAGGCCGGTACCGTGAAGGCTACGCATGTGTTCGTGGGGGATTACCAG<br>TGCGACGAGATCTACAACTACGCCGCCCTGAGCAAGAACGAGAACGGCCTAGGCATCAAGAGACTGGCCGT<br>GGTCCGGCTCGACGTGGATGACTTGGGCGCCGCCTTCATGGCCGGTTTCAGCCAGCAGGGAAACGGACAAT<br>ACTCCACTCTGTCAAGATCGGCCACATTCTCCCGGAGCATGTCGCTGTTCTTCAAAGTGTACATTAACCAG<br>TTCGCCTCCGACAAGAAGCTGAGCATTATCTACGCGGGCGGCGCCGCCGTGTTCGCCATTGGATCGTGGCA<br>GGATATCATCGCGTTCACTGTGGAACTTCGCGAAAACTTCATCAAGTGGACCAACGGGAAGCTCACCCTCT<br>CCGCGGGGATAGGGTTGTTCGCCGACAAGACTCCTATTAGCCTGATGGCTCACCAGACCGGGGAACTGGAA<br>GAGGCCGCCAAGGGCAACGAAAAGGACTCCATCTCGCTGTTCTCAAGCGACTACACTTTCAAGTTTGATAG<br>GTTCATCACTAACGTGTACGACGACAAACTGGAACAGATTAGATACTTCTTCAACCATCAAGACGAGAGGG<br>GAAAGAACTTCATCTATAAGCTTATTGAGCTTTTGAGGAACCACGACCGCATGAATATGGCACGCCTCGCC<br>TATTACCTCACTCGCCTGGAAGAACTGACCCGGGAGACTGACAGGGACAAGTTCAAGACCTTCAAGAACCT<br>GTTCTACTCCTGGTACACCAACAAGAACGATAAGGACCGGAAGGAAGCCGAGCTCGCGCTCCTGCTGTACA<br>TCTACGAAATCAGAAAGGATTAAcggcaataaaaagacagaataaaacgcacggtgttgggtcgtttgttc<br>AAGCTC |

| **Plasmid** | **pDAC309** |
|---|---|
| **Description** | Expression of Csm2 |
| **Utility** | |
| **Features** | Pcmv-FLAG-NLS-Csm2-pA |
| **Sequence** | ACATGTGAGCAAAAGGCCAGCAAAAGGCCAGGAACCGTAAAAAGGCCGCGTTGCTGGCGTTTTTCCATAGG<br>CTCCGCCCCCCTGACGAGCATCACAAAAATCGACGCTCAAGTCAGAGGTGGCGAAACCCGACAGGACTATA<br>AAGATACCAGGCGTTTCCCCCTGGAAGCTCCCTCGTGCGCTCTCCTGTTCCGACCCTGCCGCTTACCGGAT<br>ACCTGTCCGCCTTTCTCCCTTCGGGAAGCGTGGCGCTTTCTCATAGCTCACGCTGTAGGTATCTCAGTTCG<br>GTGTAGGTCGTTCGCTCCAAGCTGGGCTGTGTGCACGAACCCCCCGTTCAGCCCGACCGCTGCGCCTTATC<br>CGGTAACTATCGTCTTGAGTCCAACCCGGTAAGACACGACTTATCGCCACTGGCAGCAGCCACTGGTAACA<br>GGATTAGCAGAGCGAGGTATGTAGGCGGTGCTACAGAGTTCTTGAAGTGGTGGCCTAACTACGGCTACACT<br>AGAAGAACAGTATTTGGTATCTGCGCTCTGCTGAAGCCAGTTACCTTCGGAAAAAGAGTTGGTAGCTCTTG<br>ATCCGGCAAACAAACCACCGCTGGTAGCGGTGGTTTTTTTGTTTGCAAGCAGCAGATTACGCGCAGAAAAA<br>AAGGATCTCAAGAAGATCCTTTGATCTTTTCTACGGGGTCTGACGCTCAGTGGAACGAAAACTCACGTTAA<br>GGGATTTTGGTCATGAGATTATCAAAAAGGATCTTCACCTAGATCCTTTTAAATTAAAAATGAAGTTTTAA<br>ATCAATCTAAAGTATATATGAGTAAACTTGGTCTGACAGTTACCAATGCTTAATCAGTGAGGCACCTATCT<br>CAGCGATCTGTCTATTTCGTTCATCCATAGTTGCCTGACTCCCCGTCGTGTAGATAACTACGATACGGGAG<br>GGCTTACCATCTGGCCCCAGTGCTGCAATGATACCGCGAGACCCACGCTCACCGGCTCCAGATTTATCAGC<br>AATAAACCAGCCAGCCGGAAGGGCCGAGCGCAGAAGTGGTCCTGCAACTTTATCCGCCTCCATCCAGTCTA<br>TTAATTGTTGCCGGGAAGCTAGAGTAAGTAGTTCGCCAGTTAATAGTTTGCGCAACGTTGTTGCCATTGCT<br>ACAGGCATCGTGGTGTCACGCTCGTCGTTTGGTATGGCTTCATTCAGCTCCGGTTCCCAACGATCAAGGCG<br>AGTTACATGATCCCCCATGTTGTGCAAAAAAGCGGTTAGCTCCTTCGGTCCTCCGATCGTTGTCAGAAGTA<br>AGTTGGCCGCAGTGTTATCACTCATGGTTATGGCAGCACTGCATAATTCTCTTACTGTCATGCCATCCGTA<br>AGATGCTTTTCTGTGACTGGTGAGTACTCAACCAAGTCATTCTGAGAATAGTGTATGCGGCGACCGAGTTG<br>CTCTTGCCCGGCGTCAATACGGGATAATACCGCGCCACATAGCAGAACTTTAAAAGTGCTCATCATTGGAA<br>AACGTTCTTCGGGGCGAAAACTCTCAAGGATCTTACCGCTGTTGAGATCCAGTTCGATGTAACCCACTCGT<br>GCACCCAACTGATCTTCAGCATCTTTTACTTTCACCAGCGTTTCTGGGTGAGCAAAAACAGGAAGGCAAAA<br>TGCCGCAAAAAAGGGAATAAGGGCGACACGGAAATGTTGAATACTCATACTCTTCCTTTTTCAATATTATT<br>GAAGCATTTATCAGGGTTATTGTCTCATGAGCGGATACATATTTGAATGTATTTAGAAAAATAAACAAATA<br>GGGGTTCCGCGCACATTTCCCCGAAAAGTGCCACCTGACGTCGGATCCgacattgattattgactagttat<br>taatagtaatcaattacggggtcattagttcatagcccatatatggagttccgCGTTACATAACTTACGGT<br>AAATGGCCCGCCTGGCTGACCGCCCAACGACCCCCGCCCATTGACGTCAATAATGACGTATGTTCCCATAG<br>TAACGCCAATAGGGACTTTCCATTGACGTCAATGGGTGGAGTATTTACGGTAAACTGCCCACTTGGCAGTA<br>CATCAAGTGTATCATATGCCAAGTACGCCCCCTATTGACGTCAATGACGGTAAATGGCCCGCCTGGCATTA<br>TGCCCAGTACATGACCTTATGGGACTTTCCTACTTGGCAGTACATCTACGTATTAGTCATCGCTATTACCA<br>TGGTGATGCGGTTTTGGCAGTACATCAATGGGCGTGGATAGCGGTTTGACTCACGGGGATTTCCAAGTCTC<br>CACCCCATTGACGTCAATGGGAGTTTGTTTTGGCACCAAAATCAACGGGACTTTCCAAAATGTCGTAACAA<br>CTCCGCCCCATTGACGCAAATGGGCGGTAGGCGTGTACGGTGGGAGGTCTATATAAGCAGAGCTCGTTTAG<br>TGAACCGTCAGATCTCTAGAgccgccaccATGGATTACAAAGACGATGACGATAAGATGGCGCCTAAGAAG<br>AAACGCAAAGTGCGGGGCATGACCATCCTGACCGACGAGAACTACGTGGACATCGCCGAGAAAGCCATCCT<br>GAAGCTGGAAAGAAACACCAGAAATAGAAAGAACCCTGATGCCTTCTTCCTGACCACATCTAAGCTGCGGA |

Table S2

| | |
|---|---|
| | ACCTGCTGAGCCTGACAAGCACCCTGTTCGACGAGAGCAAGGTGAAGGAATACGACGCCCTGCTGGACAGA<br>ATCGCTTATCTGAGAGTGCAGTTCGTGTACCAGGCCGGCAGAGAGATCGCCGTGAAAGATCTGATCGAGAA<br>GGCCCAGATCCTGGAAGCTCTGAAAGAGATCAAGGACCGGGAAACCCTGCAGAGATTCTGCAGATACATGG<br>AAGCCCTGGTGGCCTACTTCAAGTTCTACGGCGGCAAGGACTGAcggcaataaaaagacagaataaaacgc<br>acggtgttgggtcgtttgttcAAGCTC |

| | |
|---|---|
| **Plasmid** | **pDAC310** |
| **Description** | Expression of Csm3 |
| **Utility** | |
| **Features** | Pcmv-FLAG-NLS-Csm3-pA |
| **Sequence** | ACATGTGAGCAAAAGGCCAGCAAAAGGCCAGGAACCGTAAAAAGGCCGCGTTGCTGGCGTTTTTCCATAGG<br>CTCCGCCCCCCTGACGAGCATCACAAAAATCGACGCTCAAGTCAGAGGTGGCGAAACCCGACAGGACTATA<br>AAGATACCAGGCGTTTCCCCCTGGAAGCTCCCTCGTGCGCTCTCCTGTTCCGACCCTGCCGCTTACCGGAT<br>ACCTGTCCGCCTTTCTCCCTTCGGGAAGCGTGGCGCTTTCTCATAGCTCACGCTGTAGGTATCTCAGTTCG<br>GTGTAGGTCGTTCGCTCCAAGCTGGGCTGTGTGCACGAACCCCCCGTTCAGCCCGACCGCTGCGCCTTATC<br>CGGTAACTATCGTCTTGAGTCCAACCCGGTAAGACACGACTTATCGCCACTGGCAGCAGCCACTGGTAACA<br>GGATTAGCAGAGCGAGGTATGTAGGCGGTGCTACAGAGTTCTTGAAGTGGTGGCCTAACTACGGCTACACT<br>AGAAGAACAGTATTTGGTATCTGCGCTCTGCTGAAGCCAGTTACCTTCGGAAAAAGAGTTGGTAGCTCTTG<br>ATCCGGCAAACAAACCACCGCTGGTAGCGGTGGTTTTTTTGTTTGCAAGCAGCAGATTACGCGCAGAAAAA<br>AAGGATCTCAAGAAGATCCTTTGATCTTTTCTACGGGGTCTGACGCTCAGTGGAACGAAAACTCACGTTAA<br>GGGATTTTGGTCATGAGATTATCAAAAAGGATCTTCACCTAGATCCTTTTAAATTAAAAATGAAGTTTTAA<br>ATCAATCTAAAGTATATATGAGTAAACTTGGTCTGACAGTTACCAATGCTTAATCAGTGAGGCACCTATCT<br>CAGCGATCTGTCTATTTCGTTCATCCATAGTTGCCTGACTCCCCGTCGTGTAGATAACTACGATACGGGAG<br>GGCTTACCATCTGGCCCCAGTGCTGCAATGATACCGCGAGACCCACGCTCACCGGCTCCAGATTTATCAGC<br>AATAAACCAGCCAGCCGGAAGGGCCGAGCGCAGAAGTGGTCCTGCAACTTTATCCGCCTCCATCCAGTCTA<br>TTAATTGTTGCCGGGAAGCTAGAGTAAGTAGTTCGCCAGTTAATAGTTTGCGCAACGTTGTTGCCATTGCT<br>ACAGGCATCGTGGTGTCACGCTCGTCGTTTGGTATGGCTTCATTCAGCTCCGGTTCCCAACGATCAAGGCG<br>AGTTACATGATCCCCCATGTTGTGCAAAAAAGCGGTTAGCTCCTTCGGTCCTCCGATCGTTGTCAGAAGTA<br>AGTTGGCCGCAGTGTTATCACTCATGGTTATGGCAGCACTGCATAATTCTCTTACTGTCATGCCATCCGTA<br>AGATGCTTTTCTGTGACTGGTGAGTACTCAACCAAGTCATTCTGAGAATAGTGTATGCGGCGACCGAGTTG<br>CTCTTGCCCGGCGTCAATACGGGATAATACCGCGCCACATAGCAGAACTTTAAAAGTGCTCATCATTGGAA<br>AACGTTCTTCGGGGCGAAAACTCTCAAGGATCTTACCGCTGTTGAGATCCAGTTCGATGTAACCCACTCGT<br>GCACCCAACTGATCTTCAGCATCTTTTACTTTCACCAGCGTTTCTGGGTGAGCAAAAACAGGAAGGCAAAA<br>TGCCGCAAAAAAGGGAATAAGGGCGACACGGAAATGTTGAATACTCATACTCTTCCTTTTTCAATATTATT<br>GAAGCATTTATCAGGGTTATTGTCTCATGAGCGGATACATATTTGAATGTATTTAGAAAAATAAACAAATA<br>GGGGTTCCGCGCACATTTCCCCGAAAAGTGCCACCTGACGTCGGATCCgacattgattattgactagttat<br>taatagtaatcaattacggggtcattagttcatagcccatatatggagttccgCGTTACATAACTTACGGT<br>AAATGGCCCGCCTGGCTGACCGCCCAACGACCCCCGCCCATTGACGTCAATAATGACGTATGTTCCCATAG<br>TAACGCCAATAGGGACTTTCCATTGACGTCAATGGGTGGAGTATTTACGGTAAACTGCCCACTTGGCAGTA<br>CATCAAGTGTATCATATGCCAAGTACGCCCCCTATTGACGTCAATGACGGTAAATGGCCCGCCTGGCATTA<br>TGCCCAGTACATGACCTTATGGGACTTTCCTACTTGGCAGTACATCTACGTATTAGTCATCGCTATTACCA<br>TGGTGATGCGGTTTTGGCAGTACATCAATGGGCGTGGATAGCGGTTTGACTCACGGGGATTTCCAAGTCTC<br>CACCCCATTGACGTCAATGGGAGTTTGTTTTGGCACCAAAATCAACGGGACTTTCCAAAATGTCGTAACAA<br>CTCCGCCCCATTGACGCAAATGGGCGGTAGGCGTGTACGGTGGGAGGTCTATATAAGCAGAGCTCGTTTAG<br>TGAACCGTCAGATCTCTAGAgccgccaccATGGATTACAAAGACGATGACGATAAGATGGCGCCTAAGAAG<br>AAACGCAAAGTGCGGGGCATGACCTTCGCCAAGATCAAATTCAGCGCCCAGATCCGGCTGGAAACCGGCCT<br>GCACATCGGAGGATCTGATGCCTTTGCCGCTATCGGCGCCATCGACAGCCCTGTGATCAAGGACCCCATCA<br>CCAACCTGCCTATCATCCCCGGCTCTAGCCTGAAGGGCAAGATGAGAACACTGCTGGCCAAGGTGTACAAC<br>GAAAAGGTGGCCGAGAAGCCTAGCGACGACAGCGACATCCTGAGCAGACTGTTCGGAAATAGCAAGGATAA<br>GCGGTTCAAGATGGGCAGACTGATCTTCCGGGACGCCTTCCTGAGCAACGCCGACGAGCTGGATTCTCTGG<br>GCGTGCGGAGCTACACCGAGGTGAAGTTCGAGAACACCATCGATAGAATCACCGCCGAGGCCAATCCTAGA<br>CAGATCGAGAGAGCCATTCGGAACTCAACATTCGACTTCGAGCTGATCTACGAGATCACTGATGAGAATGA<br>GAACCAGGTCGAGGAAGATTTCAAGGTGATCAGAGACGGCCTGAAGCTGCTGGAACTGGACTACCTGGGCG<br>GAAGCGGCTCCAGAGGCTACGGCAAAGTGGCTTTTGAGAACCTGAAAGCCACCACAGTGTTCGGCAACTAC<br>GACGTGAAAACCCTGAACGAGCTGCTGACCGCCGAAGTGTGAcggcaataaaaagacagaataaaacgcac<br>ggtgttgggtcgtttgttcAAGCTC |

| | |
|---|---|
| **Plasmid** | **pDAC327** |
| **Description** | Expression of Csm3(RNase mut) |
| **Utility** | |
| **Features** | Pcmv-FLAG-NLS-Csm3(RNase mut)-pA |

Table S2

| | |
|---|---|
| **Sequence** | ACATGTGAGCAAAAGGCCAGCAAAAGGCCAGGAACCGTAAAAAGGCCGCGTTGCTGGCGTTTTTCCATAGG<br>CTCCGCCCCCCTGACGAGCATCACAAAAATCGACGCTCAAGTCAGAGGTGGCGAAACCCGACAGGACTATA<br>AAGATACCAGGCGTTTCCCCCTGGAAGCTCCCTCGTGCGCTCTCCTGTTCCGACCCTGCCGCTTACCGGAT<br>ACCTGTCCGCCTTTCTCCCTTCGGGAAGCGTGGCGCTTTCTCATAGCTCACGCTGTAGGTATCTCAGTTCG<br>GTGTAGGTCGTTCGCTCCAAGCTGGGCTGTGTGCACGAACCCCCCGTTCAGCCCGACCGCTGCGCCTTATC<br>CGGTAACTATCGTCTTGAGTCCAACCCGGTAAGACACGACTTATCGCCACTGGCAGCAGCCACTGGTAACA<br>GGATTAGCAGAGCGAGGTATGTAGGCGGTGCTACAGAGTTCTTGAAGTGGTGGCCTAACTACGGCTACACT<br>AGAAGAACAGTATTTGGTATCTGCGCTCTGCTGAAGCCAGTTACCTTCGGAAAAAGAGTTGGTAGCTCTTG<br>ATCCGGCAAACAAACCACCGCTGGTAGCGGTGGTTTTTTTGTTTGCAAGCAGCAGATTACGCGCAGAAAAA<br>AAGGATCTCAAGAAGATCCTTTGATCTTTTCTACGGGGTCTGACGCTCAGTGGAACGAAAACTCACGTTAA<br>GGGATTTTGGTCATGAGATTATCAAAAAGGATCTTCACCTAGATCCTTTTAAATTAAAAATGAAGTTTTAA<br>ATCAATCTAAAGTATATATGAGTAAACTTGGTCTGACAGTTACCAATGCTTAATCAGTGAGGCACCTATCT<br>CAGCGATCTGTCTATTTCGTTCATCCATAGTTGCCTGACTCCCCGTCGTGTAGATAACTACGATACGGGAG<br>GGCTTACCATCTGGCCCCAGTGCTGCAATGATACCGCGAGACCCACGCTCACCGGCTCCAGATTTATCAGC<br>AATAAACCAGCCAGCCGGAAGGGCCGAGCGCAGAAGTGGTCCTGCAACTTTATCCGCCTCCATCCAGTCTA<br>TTAATTGTTGCCGGGAAGCTAGAGTAAGTAGTTCGCCAGTTAATAGTTTGCGCAACGTTGTTGCCATTGCT<br>ACAGGCATCGTGGTGTCACGCTCGTCGTTTGGTATGGCTTCATTCAGCTCCGGTTCCCAACGATCAAGGCG<br>AGTTACATGATCCCCCATGTTGTGCAAAAAAGCGGTTAGCTCCTTCGGTCCTCCGATCGTTGTCAGAAGTA<br>AGTTGGCCGCAGTGTTATCACTCATGGTTATGGCAGCACTGCATAATTCTCTTACTGTCATGCCATCCGTA<br>AGATGCTTTTCTGTGACTGGTGAGTACTCAACCAAGTCATTCTGAGAATAGTGTATGCGGCGACCGAGTTG<br>CTCTTGCCCGGCGTCAATACGGGATAATACCGCGCCACATAGCAGAACTTTAAAAGTGCTCATCATTGGAA<br>AACGTTCTTCGGGGCGAAAACTCTCAAGGATCTTACCGCTGTTGAGATCCAGTTCGATGTAACCCACTCGT<br>GCACCCAACTGATCTTCAGCATCTTTTACTTTCACCAGCGTTTCTGGGTGAGCAAAAACAGGAAGGCAAAA<br>TGCCGCAAAAAAGGGAATAAGGGCGACACGGAAATGTTGAATACTCATACTCTTCCTTTTTCAATATTATT<br>GAAGCATTTATCAGGGTTATTGTCTCATGAGCGGATACATATTTGAATGTATTTAGAAAAATAAACAAATA<br>GGGGTTCCGCGCACATTTCCCCGAAAAGTGCCACCTGACGTCGGATCCgacattgattattgactagttat<br>taatagtaatcaattacggggtcattagttcatagcccatatatggagttccgCGTTACATAACTTACGGT<br>AAATGGCCCGCCTGGCTGACCGCCCAACGACCCCCGCCCATTGACGTCAATAATGACGTATGTTCCCATAG<br>TAACGCCAATAGGGACTTTCCATTGACGTCAATGGGTGGAGTATTTACGGTAAACTGCCCACTTGGCAGTA<br>CATCAAGTGTATCATATGCCAAGTACGCCCCCTATTGACGTCAATGACGGTAAATGGCCCGCCTGGCATTA<br>TGCCCAGTACATGACCTTATGGGACTTTCCTACTTGGCAGTACATCTACGTATTAGTCATCGCTATTACCA<br>TGGTGATGCGGTTTTGGCAGTACATCAATGGGCGTGGATAGCGGTTTGACTCACGGGGATTTCCAAGTCTC<br>CACCCCATTGACGTCAATGGGAGTTTGTTTTGGCACCAAAATCAACGGGACTTTCCAAAATGTCGTAACAA<br>CTCCGCCCCATTGACGCAAATGGGCGGTAGGCGTGTACGGTGGGAGGTCTATATAAGCAGAGCTCGTTTAG<br>TGAACCGTCAGATCTCTAGAgccgccaccATGGATTACAAAGACGATGACGATAAGATGGCGCCTAAGAAG<br>AAACGCAAAGTGCGGGGCATGACCTTCGCCAAGATCAAATTCAGCGCCCAGATCCGGCTGGAAACCGGCCT<br>GCACATCGGAGGATCTGATGCCTTTGCCGCTATCGGCGCCATCGCCAGCCCTGTGATCAAGGACCCCATCA<br>CCAACCTGCCTATCATCCCCGGCTCTAGCCTGAAGGGCAAGATGAGAACACTGCTGGCCAAGGTGTACAAC<br>GAAAAGGTGGCCGAGAAGCCTAGCGACGACAGCGACATCCTGAGCAGACTGTTCGGAAATAGCAAGGATAA<br>GCGGTTCAAGATGGGCAGACTGATCTTCCGGGACGCCTTCCTGAGCAACGCCGACGAGCTGGATTCTCTGG<br>GCGTGCGGAGCTACACCGAGGTGAAGTTCGAGAACACCATCGATAGAATCACCGCCGAGGCCAATCCTAGA<br>CAGATCGAGAGAGCCATTCGGAACTCAACATTCGACTTCGAGCTGATCTACGAGATCACTGATGAGAATGA<br>GAACCAGGTCGAGGAAGATTTCAAGGTGATCAGAGACGGCCTGAAGCTGCTGGAACTGGACTACCTGGGCG<br>GAAGCGGCTCCAGAGGCTACGGCAAAGTGGCTTTTGAGAACCTGAAAGCCACCACAGTGTTCGGCAACTAC<br>GACGTGAAAACCCTGAACGAGCTGCTGACCGCCGAAGTGTGAcggcaataaaaagacagaataaaacgcac<br>ggtgttgggtcgtttgttcAAGCTC |

| | |
|---|---|
| **Plasmid** | **pDAC339** |
| **Description** | Expression of Csm4 |
| **Utility** | |
| **Features** | Pcmv-FLAG-NLS-Csm4-pA |
| **Sequence** | ACATGTGAGCAAAAGGCCAGCAAAAGGCCAGGAACCGTAAAAAGGCCGCGTTGCTGGCGTTTTTCCATAGG<br>CTCCGCCCCCCTGACGAGCATCACAAAAATCGACGCTCAAGTCAGAGGTGGCGAAACCCGACAGGACTATA<br>AAGATACCAGGCGTTTCCCCCTGGAAGCTCCCTCGTGCGCTCTCCTGTTCCGACCCTGCCGCTTACCGGAT<br>ACCTGTCCGCCTTTCTCCCTTCGGGAAGCGTGGCGCTTTCTCATAGCTCACGCTGTAGGTATCTCAGTTCG<br>GTGTAGGTCGTTCGCTCCAAGCTGGGCTGTGTGCACGAACCCCCCGTTCAGCCCGACCGCTGCGCCTTATC<br>CGGTAACTATCGTCTTGAGTCCAACCCGGTAAGACACGACTTATCGCCACTGGCAGCAGCCACTGGTAACA<br>GGATTAGCAGAGCGAGGTATGTAGGCGGTGCTACAGAGTTCTTGAAGTGGTGGCCTAACTACGGCTACACT<br>AGAAGAACAGTATTTGGTATCTGCGCTCTGCTGAAGCCAGTTACCTTCGGAAAAAGAGTTGGTAGCTCTTG<br>ATCCGGCAAACAAACCACCGCTGGTAGCGGTGGTTTTTTTGTTTGCAAGCAGCAGATTACGCGCAGAAAAA<br>AAGGATCTCAAGAAGATCCTTTGATCTTTTCTACGGGGTCTGACGCTCAGTGGAACGAAAACTCACGTTAA<br>GGGATTTTGGTCATGAGATTATCAAAAAGGATCTTCACCTAGATCCTTTTAAATTAAAAATGAAGTTTTAA<br>ATCAATCTAAAGTATATATGAGTAAACTTGGTCTGACAGTTACCAATGCTTAATCAGTGAGGCACCTATCT |

Table S2

| | |
|---|---|
| | CAGCGATCTGTCTATTTCGTTCATCCATAGTTGCCTGACTCCCCGTCGTGTAGATAACTACGATACGGGAG<br>GGCTTACCATCTGGCCCCAGTGCTGCAATGATACCGCGAGACCCACGCTCACCGGCTCCAGATTTATCAGC<br>AATAAACCAGCCAGCCGGAAGGGCCGAGCGCAGAAGTGGTCCTGCAACTTTATCCGCCTCCATCCAGTCTA<br>TTAATTGTTGCCGGGAAGCTAGAGTAAGTAGTTCGCCAGTTAATAGTTTGCGCAACGTTGTTGCCATTGCT<br>ACAGGCATCGTGGTGTCACGCTCGTCGTTTGGTATGGCTTCATTCAGCTCCGGTTCCCAACGATCAAGGCG<br>AGTTACATGATCCCCCATGTTGTGCAAAAAAGCGGTTAGCTCCTTCGGTCCTCCGATCGTTGTCAGAAGTA<br>AGTTGGCCGCAGTGTTATCACTCATGGTTATGGCAGCACTGCATAATTCTCTTACTGTCATGCCATCCGTA<br>AGATGCTTTTCTGTGACTGGTGAGTACTCAACCAAGTCATTCTGAGAATAGTGTATGCGGCGACCGAGTTG<br>CTCTTGCCCGGCGTCAATACGGGATAATACCGCGCCACATAGCAGAACTTTAAAAGTGCTCATCATTGGAA<br>AACGTTCTTCGGGGCGAAAACTCTCAAGGATCTTACCGCTGTTGAGATCCAGTTCGATGTAACCCACTCGT<br>GCACCCAACTGATCTTCAGCATCTTTTACTTTCACCAGCGTTTCTGGGTGAGCAAAAACAGGAAGGCAAAA<br>TGCCGCAAAAAAGGGAATAAGGGCGACACGGAAATGTTGAATACTCATACTCTTCCTTTTTCAATATTATT<br>GAAGCATTTATCAGGGTTATTGTCTCATGAGCGGATACATATTTGAATGTATTTAGAAAAATAAACAAATA<br>GGGGTTCCGCGCACATTTCCCCGAAAAGTGCCACCTGACGTCGGATCCgacattgattattgactagttat<br>taatagtaatcaattacggggtcattagttcatagcccatatatggagttccgCGTTACATAACTTACGGT<br>AAATGGCCCGCCTGGCTGACCGCCCAACGACCCCCGCCCATTGACGTCAATAATGACGTATGTTCCCATAG<br>TAACGCCAATAGGGACTTTCCATTGACGTCAATGGGTGGAGTATTTACGGTAAACTGCCCACTTGGCAGTA<br>CATCAAGTGTATCATATGCCAAGTACGCCCCCTATTGACGTCAATGACGGTAAATGGCCCGCCTGGCATTA<br>TGCCCAGTACATGACCTTATGGGACTTTCCTACTTGGCAGTACATCTACGTATTAGTCATCGCTATTACCA<br>TGGTGATGCGGTTTTGGCAGTACATCAATGGGCGTGGATAGCGGTTTGACTCACGGGGATTTCCAAGTCTC<br>CACCCCATTGACGTCAATGGGAGTTTGTTTTGGCACCAAAATCAACGGGACTTTCCAAAATGTCGTAACAA<br>CTCCGCCCCATTGACGCAAATGGGCGGTAGGCGTGTACGGTGGGAGGTCTATATAAGCAGAGCTCGTTTAG<br>TGAACCGTCAGATCTCTAGAgccgccaccATGGATTACAAAGACGATGACGATAAGATGGCGCCTAAGAAG<br>AAACGCAAAGTGCGGGGCATGACTTACAAGCTCTACATTATGACCTTTCAAAACGCCCACTTCGGTTCCGG<br>CACTCTGGACTCATCGAAGCTGACCTTCTCCGCGGATAGAATCTTCTCGGCACTCGTGCTCGAGGCTCTGA<br>AGATGGGAAAGCTCGACGCCTTCTTGGCCGAGGCCAACCAGGATAAGTTCACTCTGACCGACGCGTTCCCA<br>TTCCAATTCGGTCCTTTCCTGCCGAAACCGATTGGTTACCCCAAGCACGACCAGATCGACCAGTCTGTGGA<br>CGTGAAGGAAGTCCGCCGCCAAGCGAAGCTGTCCAAAAAGCTCCAGTTCCTGGCTCTGGAAAACGTCGACG<br>ACTACCTGAACGGAGAGCTGTTTGAGAATGAGGAACACGCCGTGATCGACACAGTGACCAAGAACCAGCCC<br>CATAAAGATGATAATCTGTACCAAGTGGCCACCACTCGGTTCTCGAACGACACCTCCCTTTACGTGATCGC<br>CAACGAATCCGATCTGCTGAACGAACTGATGAGCAGCCTTCAGTACTCCGGGCTGGGCGGCAAAAGGTCCT<br>CAGGATTCGGCAGATTTGAGCTGGACATCCAGAACATTCCCTTGGAACTGTCCGACCGGCTGACGAAGAAC<br>CACAGCGACAAGGTCATGTCACTTACCACCGCCCTCCCGGTGGACGCTGATCTCGAGGAAGCGATGGAAGA<br>TGGCCATTACCTGTTGACCAAGTCGTCCGGATTCGCATTCTCCCACGCCACCAACGAAAACTATCGGAAGC<br>AGGACCTGTACAAGTTCGCCTCCGGGAGCACCTTCAGCAAGACTTTCGAGGGACAGATCGTGGACGTGCGC<br>CCTCTCGATTTCCCTCACGCCGTGCTGAACTACGCCAAGCCGCTGTTCTTTAAGCTCGAAGTCTAAcggca<br>ataaaaagacagaataaaacgcacggtgttgggtcgtttgttcAAGCTC |

| **Plasmid** | **pDAC312** |
|---|---|
| **Description** | Expression of Csm5 |
| **Utility** | |
| **Features** | Pcmv-FLAG-NLS-Csm5-pA |
| **Sequence** | ACATGTGAGCAAAAGGCCAGCAAAAGGCCAGGAACCGTAAAAAGGCCGCGTTGCTGGCGTTTTTCCATAGG<br>CTCCGCCCCCCTGACGAGCATCACAAAAATCGACGCTCAAGTCAGAGGTGGCGAAACCCGACAGGACTATA<br>AAGATACCAGGCGTTTCCCCCTGGAAGCTCCCTCGTGCGCTCTCCTGTTCCGACCCTGCCGCTTACCGGAT<br>ACCTGTCCGCCTTTCTCCCTTCGGGAAGCGTGGCGCTTTCTCATAGCTCACGCTGTAGGTATCTCAGTTCG<br>GTGTAGGTCGTTCGCTCCAAGCTGGGCTGTGTGCACGAACCCCCCGTTCAGCCCGACCGCTGCGCCTTATC<br>CGGTAACTATCGTCTTGAGTCCAACCCGGTAAGACACGACTTATCGCCACTGGCAGCAGCCACTGGTAACA<br>GGATTAGCAGAGCGAGGTATGTAGGCGGTGCTACAGAGTTCTTGAAGTGGTGGCCTAACTACGGCTACACT<br>AGAAGAACAGTATTTGGTATCTGCGCTCTGCTGAAGCCAGTTACCTTCGGAAAAAGAGTTGGTAGCTCTTG<br>ATCCGGCAAACAAACCACCGCTGGTAGCGGTGGTTTTTTTGTTTGCAAGCAGCAGATTACGCGCAGAAAAA<br>AAGGATCTCAAGAAGATCCTTTGATCTTTTCTACGGGGTCTGACGCTCAGTGGAACGAAAACTCACGTTAA<br>GGGATTTTGGTCATGAGATTATCAAAAAGGATCTTCACCTAGATCCTTTTAAATTAAAAATGAAGTTTTAA<br>ATCAATCTAAAGTATATATGAGTAAACTTGGTCTGACAGTTACCAATGCTTAATCAGTGAGGCACCTATCT<br>CAGCGATCTGTCTATTTCGTTCATCCATAGTTGCCTGACTCCCCGTCGTGTAGATAACTACGATACGGGAG<br>GGCTTACCATCTGGCCCCAGTGCTGCAATGATACCGCGAGACCCACGCTCACCGGCTCCAGATTTATCAGC<br>AATAAACCAGCCAGCCGGAAGGGCCGAGCGCAGAAGTGGTCCTGCAACTTTATCCGCCTCCATCCAGTCTA<br>TTAATTGTTGCCGGGAAGCTAGAGTAAGTAGTTCGCCAGTTAATAGTTTGCGCAACGTTGTTGCCATTGCT<br>ACAGGCATCGTGGTGTCACGCTCGTCGTTTGGTATGGCTTCATTCAGCTCCGGTTCCCAACGATCAAGGCG<br>AGTTACATGATCCCCCATGTTGTGCAAAAAAGCGGTTAGCTCCTTCGGTCCTCCGATCGTTGTCAGAAGTA<br>AGTTGGCCGCAGTGTTATCACTCATGGTTATGGCAGCACTGCATAATTCTCTTACTGTCATGCCATCCGTA<br>AGATGCTTTTCTGTGACTGGTGAGTACTCAACCAAGTCATTCTGAGAATAGTGTATGCGGCGACCGAGTTG<br>CTCTTGCCCGGCGTCAATACGGGATAATACCGCGCCACATAGCAGAACTTTAAAAGTGCTCATCATTGGAA |

Table S2

| | |
|---|---|
| | AACGTTCTTCGGGGCGAAAACTCTCAAGGATCTTACCGCTGTTGAGATCCAGTTCGATGTAACCCACTCGT<br>GCACCCAACTGATCTTCAGCATCTTTTACTTTCACCAGCGTTTCTGGGTGAGCAAAAACAGGAAGGCAAAA<br>TGCCGCAAAAAAGGGAATAAGGGCGACACGGAAATGTTGAATACTCATACTCTTCCTTTTTCAATATTATT<br>GAAGCATTTATCAGGGTTATTGTCTCATGAGCGGATACATATTTGAATGTATTTAGAAAAATAAACAAATA<br>GGGGTTCCGCGCACATTTCCCCGAAAAGTGCCACCTGACGTCGGATCCgacattgattattgactagttat<br>taatagtaatcaattacggggtcattagttcatagcccatatatggagttccgCGTTACATAACTTACGGT<br>AAATGGCCCGCCTGGCTGACCGCCCAACGACCCCCGCCCATTGACGTCAATAATGACGTATGTTCCCATAG<br>TAACGCCAATAGGGACTTTCCATTGACGTCAATGGGTGGAGTATTTACGGTAAACTGCCCACTTGGCAGTA<br>CATCAAGTGTATCATATGCCAAGTACGCCCCCTATTGACGTCAATGACGGTAAATGGCCCGCCTGGCATTA<br>TGCCCAGTACATGACCTTATGGGACTTTCCTACTTGGCAGTACATCTACGTATTAGTCATCGCTATTACCA<br>TGGTGATGCGGTTTTGGCAGTACATCAATGGGCGTGGATAGCGGTTTGACTCACGGGGATTTCCAAGTCTC<br>CACCCCATTGACGTCAATGGGAGTTTGTTTTGGCACCAAAATCAACGGGACTTTCCAAAATGTCGTAACAA<br>CTCCGCCCCATTGACGCAAATGGGCGGTAGGCGTGTACGGTGGGAGGTCTATATAAGCAGAGCTCGTTTAG<br>TGAACCGTCAGATCTCTAGAgccgccaccATGGATTACAAAGACGATGACGATAAGATGGCGCCTAAGAAG<br>AAACGCAAAGTGCGGGGCATGAAAAATGACTACCGGACCTTCAAGCTGAGCCTGCTGACCCTGGCTCCTAT<br>CCACATCGGCAACGGCGAGAAGTACACCAGCAGAGAATTCATCTACGAGAACAAGAAGTTCTACTTCCCCG<br>ACATGGGCAAGTTCTACAACAAGATGGTGGAAAAGAGACTGGCCGAGAAGTTCGAGGCCTTCCTGATCCAG<br>ACCAGACCCAACGCCAGAAACAACCGGCTGATTTCTTTTCTGAACGACAACAGAATCGCCGAAAGATCTTT<br>TGGCGGCTACAGCATCAGTGAAACCGGCCTGGAATCTGATAAGAACCCTAACAGCGCCGGAGCTATCAACG<br>AGGTGAACAAATTCATCCGGGACGCCTTCGGAAATCCTTACATCCCAGGCAGCAGCCTGAAGGGCGCCATC<br>CGCACCATCCTGATGAACACCACACCTAAGTGGAACAACGAGAACGCCGTGAACGACTTCGGCAGATTCCC<br>AAAGGAAAACAAGAACCTGATCCCTTGGGGACCTAAGAAAGGCAAGGAATACGACGACCTGTTCAACGCCA<br>TCAGAGTGTCCGACAGCAAGCCCTTCGACAACAAAAGCCTGATCCTCGTGCAGAAGTGGGACTACAGCGCC<br>AAAACCAACAAGGCCAAGCCTCTGCCTCTGTACAGAGAGTCTATCAGCCCTCTGACCAAGATCGAGTTCGA<br>GATAACAACAACCACTGATGAGGCCGGCAGACTGATCGAGGAACTGGGAAAGCGGGCCCAGGCCTTTTATA<br>AGGACTACAAGGCCTTTTTCCTGTCTGAATTCCCTGATGATAAGATCCAGGCTAATCTGCAATACCCCATC<br>TACCTGGGCGCCGGCAGCGGCGCTTGGACAAAGACCCTGTTTAAGCAGGCCGACGGCATCCTGCAGCGGAG<br>ATACTCCAGAATGAAAACCAAGATGGTCAAGAAGGGCGTGCTGAAGCTGACAAAGGCCCCTCTGAAAACAG<br>TGAAGATCCCCAGCGGCAACCACAGCCTGGTGAAGAATCACGAGAGCTTCTACGAGATGGGCAAAGCCAAC<br>TTCATGATCAAGGAAATCGACAAGTGAcggcaataaaaagacagaataaaacgcacggtgttgggtcgttt<br>gttcAAGCTC |

| **Plasmid** | **pDAC307** |
|---|---|
| **Description** | Expression of Cas6 |
| **Utility** | |
| **Features** | Pcmv-FLAG-NLS-Cas6-pA |
| **Sequence** | ACATGTGAGCAAAAGGCCAGCAAAAGGCCAGGAACCGTAAAAAGGCCGCGTTGCTGGCGTTTTTCCATAGG<br>CTCCGCCCCCCTGACGAGCATCACAAAAATCGACGCTCAAGTCAGAGGTGGCGAAACCCGACAGGACTATA<br>AAGATACCAGGCGTTTCCCCCTGGAAGCTCCCTCGTGCGCTCTCCTGTTCCGACCCTGCCGCTTACCGGAT<br>ACCTGTCCGCCTTTCTCCCTTCGGGAAGCGTGGCGCTTTCTCATAGCTCACGCTGTAGGTATCTCAGTTCG<br>GTGTAGGTCGTTCGCTCCAAGCTGGGCTGTGTGCACGAACCCCCCGTTCAGCCCGACCGCTGCGCCTTATC<br>CGGTAACTATCGTCTTGAGTCCAACCCGGTAAGACACGACTTATCGCCACTGGCAGCAGCCACTGGTAACA<br>GGATTAGCAGAGCGAGGTATGTAGGCGGTGCTACAGAGTTCTTGAAGTGGTGGCCTAACTACGGCTACACT<br>AGAAGAACAGTATTTGGTATCTGCGCTCTGCTGAAGCCAGTTACCTTCGGAAAAAGAGTTGGTAGCTCTTG<br>ATCCGGCAAACAAACCACCGCTGGTAGCGGTGGTTTTTTTGTTTGCAAGCAGCAGATTACGCGCAGAAAAA<br>AAGGATCTCAAGAAGATCCTTTGATCTTTTCTACGGGGTCTGACGCTCAGTGGAACGAAAACTCACGTTAA<br>GGGATTTTGGTCATGAGATTATCAAAAAGGATCTTCACCTAGATCCTTTTAAATTAAAAATGAAGTTTTAA<br>ATCAATCTAAAGTATATATGAGTAAACTTGGTCTGACAGTTACCAATGCTTAATCAGTGAGGCACCTATCT<br>CAGCGATCTGTCTATTTCGTTCATCCATAGTTGCCTGACTCCCCGTCGTGTAGATAACTACGATACGGGAG<br>GGCTTACCATCTGGCCCCAGTGCTGCAATGATACCGCGAGACCCACGCTCACCGGCTCCAGATTTATCAGC<br>AATAAACCAGCCAGCCGGAAGGGCCGAGCGCAGAAGTGGTCCTGCAACTTTATCCGCCTCCATCCAGTCTA<br>TTAATTGTTGCCGGGAAGCTAGAGTAAGTAGTTCGCCAGTTAATAGTTTGCGCAACGTTGTTGCCATTGCT<br>ACAGGCATCGTGGTGTCACGCTCGTCGTTTGGTATGGCTTCATTCAGCTCCGGTTCCCAACGATCAAGGCG<br>AGTTACATGATCCCCCATGTTGTGCAAAAAAGCGGTTAGCTCCTTCGGTCCTCCGATCGTTGTCAGAAGTA<br>AGTTGGCCGCAGTGTTATCACTCATGGTTATGGCAGCACTGCATAATTCTCTTACTGTCATGCCATCCGTA<br>AGATGCTTTTCTGTGACTGGTGAGTACTCAACCAAGTCATTCTGAGAATAGTGTATGCGGCGACCGAGTTG<br>CTCTTGCCCGGCGTCAATACGGGATAATACCGCGCCACATAGCAGAACTTTAAAAGTGCTCATCATTGGAA<br>AACGTTCTTCGGGGCGAAAACTCTCAAGGATCTTACCGCTGTTGAGATCCAGTTCGATGTAACCCACTCGT<br>GCACCCAACTGATCTTCAGCATCTTTTACTTTCACCAGCGTTTCTGGGTGAGCAAAAACAGGAAGGCAAAA<br>TGCCGCAAAAAAGGGAATAAGGGCGACACGGAAATGTTGAATACTCATACTCTTCCTTTTTCAATATTATT<br>GAAGCATTTATCAGGGTTATTGTCTCATGAGCGGATACATATTTGAATGTATTTAGAAAAATAAACAAATA<br>GGGGTTCCGCGCACATTTCCCCGAAAAGTGCCACCTGACGTCGGATCCgacattgattattgactagttat<br>taatagtaatcaattacggggtcattagttcatagcccatatatggagttccgCGTTACATAACTTACGGT |

| | |
|---|---|
| | AAATGGCCCGCCTGGCTGACCGCCCAACGACCCCCGCCCATTGACGTCAATAATGACGTATGTTCCCATAG<br>TAACGCCAATAGGGACTTTCCATTGACGTCAATGGGTGGAGTATTTACGGTAAACTGCCCACTTGGCAGTA<br>CATCAAGTGTATCATATGCCAAGTACGCCCCCTATTGACGTCAATGACGGTAAATGGCCCGCCTGGCATTA<br>TGCCCAGTACATGACCTTATGGGACTTTCCTACTTGGCAGTACATCTACGTATTAGTCATCGCTATTACCA<br>TGGTGATGCGGTTTTGGCAGTACATCAATGGGCGTGGATAGCGGTTTGACTCACGGGGATTTCCAAGTCTC<br>CACCCCATTGACGTCAATGGGAGTTTGTTTTGGCACCAAAATCAACGGGACTTTCCAAAATGTCGTAACAA<br>CTCCGCCCCATTGACGCAAATGGGCGGTAGGCGTGTACGGTGGGAGGTCTATATAAGCAGAGCTCGTTTAG<br>TGAACCGTCAGATCTCTAGAgccgccaccATGGATTACAAAGACGATGACGATAAGATGGCGCCTAAGAAG<br>AAACGCAAAGTGCGGGGCATGAAAAAGCTCGTGTTCACCTTTAAGCGGATCGACCACCCTGCTCAGGACCT<br>GGCCGTGAAATTCCACGGCTTCCTGATGGAACAGCTGGATAGCGACTACGTGGACTACCTGCACCAGCAGC<br>AGACCAACCCCTACGCCACAAAGGTGATCCAGGGCAAAGAGAACACCCAGTGGGTCGTGCATCTGCTGACA<br>GACGACATCGAGGACAAGGTGTTCATGACCCTGCTGCAGATCAAGGAAGTGTCCCTGAACGACCTGCCTAA<br>GTTGTCTGTGGAAAAGGTGGAAATCCAGGAGCTGGGCGCTGATAAGCTGCTCGAGATCTTCAACAGCGAGG<br>AAAACCAGACCTACTTCAGCATCATCTTCGAGACACCTACAGGCTTTAAAAGCCAGGGCAGCTACGTGATC<br>TTCCCCAGCATGCGGCTGATCTTTCAGAGCCTGATGCAGAAGTACGGCAGACTGGTGGAAAACCAGCCTGA<br>GATCGAGGAAGATACCCTGGACTACCTGAGCGAGCACAGCACCATCACCAATTACAGACTGGAAACAAGCT<br>ACTTCAGAGTGCATAGACAGAGAATCCCCGCCTTCCGGGGCAAGCTGACCTTCAAGGTGCAGGGAGCCCAG<br>ACACTGAAGGCCTACGTGAAGATGCTGCTGACCTTCGGCGAGTACAGCGGCCTGGGCATGAAAACCAGCCT<br>GGGAATGGGCGGCATCAAGCTGGAAGAAAGAAAGGACTGAcggcaataaaaagacagaataaaacgcacgg<br>tgttgggtcgtttgttcAAGCTC |

| **Plasmid** | **pDAC324** |
|---|---|
| **Description** | Expression of crRNA |
| **Utility** | |
| **Features** | Pu6-crRNA-pT |
| **Sequence** | TCGCGCGTTTCGGTGATGACGGTGAAAACCTCTGACACATGCAGCTCCCGGAGACGGTCACAGCTTGTCTG<br>TAAGCGGATGCCGGGAGCAGACAAGCCCGTCAGGGCGCGTCAGCGGGTGTTGGCGGGTGTCGGGGCTGGCT<br>TAACTATGCGGCATCAGAGCAGATTGTACTGAGAGTGCACCATATGGAGGGCCTATTTCCCATGATTCCTT<br>CATATTTGCATATACGATACAAGGCTGTTAGAGAGATAATTAGAATTAATTTGACTGTAAACACAAAGATA<br>TTAGTACAAAATACGTGACGTAGAAAGTAATAATTTCTTGGGTAGTTTGCAGTTTTAAAATTATGTTTTAA<br>AATGGACTATCATATGCTTACCGTAACTTGAAAGTATTTCGATTTCTTGGCTTTATATATCTTGTGGAAAG<br>GACGAAACACCGATATAAACCTAATTACCTCGAGAGGGGACGGAAACCCGTCTTCGATGAAGCGATTCAGA<br>AGACTTGATATAAACCTAATTACCTCGAGAGGGGACTTTTTTACATGTGAGCAAAAGGCCAGCAAAAGGCC<br>AGGAACCGTAAAAAGGCCGCGTTGCTGGCGTTTTTCCATAGGCTCCGCCCCCCTGACGAGCATCACAAAAA<br>TCGACGCTCAAGTCAGAGGTGGCGAAACCCGACAGGACTATAAAGATACCAGGCGTTTCCCCCTGGAAGCT<br>CCCTCGTGCGCTCTCCTGTTCCGACCCTGCCGCTTACCGGATACCTGTCCGCCTTTCTCCCTTCGGGAAGC<br>GTGGCGCTTTCTCATAGCTCACGCTGTAGGTATCTCAGTTCGGTGTAGGTCGTTCGCTCCAAGCTGGGCTG<br>TGTGCACGAACCCCCCGTTCAGCCCGACCGCTGCGCCTTATCCGGTAACTATCGTCTTGAGTCCAACCCGG<br>TAAGACACGACTTATCGCCACTGGCAGCAGCCACTGGTAACAGGATTAGCAGAGCGAGGTATGTAGGCGGT<br>GCTACAGAGTTCTTGAAGTGGTGGCCTAACTACGGCTACACTAGAAGAACAGTATTTGGTATCTGCGCTCT<br>GCTGAAGCCAGTTACCTTCGGAAAAAGAGTTGGTAGCTCTTGATCCGGCAAACAAACCACCGCTGGTAGCG<br>GTGGTTTTTTTGTTTGCAAGCAGCAGATTACGCGCAGAAAAAAAGGATCTCAAGAAGATCCTTTGATCTTT<br>TCTACGGGGTCTGACGCTCAGTGGAACGAAAACTCACGTTAAGGGATTTTGGTCATGAGATTATCAAAAAG<br>GATCTTCACCTAGATCCTTTTAAATTAAAAATGAAGTTTTAAATCAATCTAAAGTATATATGAGTAAACTT<br>GGTCTGACAGTTACCAATGCTTAATCAGTGAGGCACCTATCTCAGCGATCTGTCTATTTCGTTCATCCATA<br>GTTGCCTGACTCCCCGTCGTGTAGATAACTACGATACGGGAGGGCTTACCATCTGGCCCCAGTGCTGCAAT<br>GATACCGCGAGACCCACGCTCACCGGCTCCAGATTTATCAGCAATAAACCAGCCAGCCGGAAGGGCCGAGC<br>GCAGAAGTGGTCCTGCAACTTTATCCGCCTCCATCCAGTCTATTAATTGTTGCCGGGAAGCTAGAGTAAGT<br>AGTTCGCCAGTTAATAGTTTGCGCAACGTTGTTGCCATTGCTACAGGCATCGTGGTGTCACGCTCGTCGTT<br>TGGTATGGCTTCATTCAGCTCCGGTTCCCAACGATCAAGGCGAGTTACATGATCCCCCATGTTGTGCAAAA<br>AAGCGGTTAGCTCCTTCGGTCCTCCGATCGTTGTCAGAAGTAAGTTGGCCGCAGTGTTATCACTCATGGTT<br>ATGGCAGCACTGCATAATTCTCTTACTGTCATGCCATCCGTAAGATGCTTTTCTGTGACTGGTGAGTACTC<br>AACCAAGTCATTCTGAGAATAGTGTATGCGGCGACCGAGTTGCTCTTGCCCGGCGTCAATACGGATAATA<br>CCGCGCCACATAGCAGAACTTTAAAAGTGCTCATCATTGGAAAACGTTCTTCGGGGCGAAAACTCTCAAGG<br>ATCTTACCGCTGTTGAGATCCAGTTCGATGTAACCCACTCGTGCACCCAACTGATCTTCAGCATCTTTTAC<br>TTTCACCAGCGTTTCTGGGTGAGCAAAAACAGGAAGGCAAAATGCCGCAAAAAAGGGAATAAGGGCGACAC<br>GGAAATGTTGAATACTCATACTCTTCCTTTTTCAATATTATTGAAGCATTTATCAGGGTTATTGTCTCATG<br>AGCGGATACATATTTGAATGTATTTAGAAAAATAAACAAATAGGGGTTCCGCGCACATTTCCCCGAAAAGT<br>GCCACCTGACGTCTAAGAAACCATTATTATCATGACATTAACCTATAAAAATAGGCGTATCACGAGGCCCT<br>TTCGTC |

| **Plasmid** | **pDAC759** |
|---|---|

Table S2

| | |
|---|---|
| **Description** | Expression of Csm complex (DNase/cA mut) from single promoter; RFP backbone |
| **Utility** | RNA KD |
| **Features** | Pcmv-FLAG-NLS-Csm5-2A-FLAG-NLS-Csm4-2A-FLAG-NLS-Csm3-2A-FLAG-NLS-Csm2-pA; Pcmv-RFP-2A-FLAG-NLS-Cas6-2A-FLAG-NLS-Csm1(Dnase/cA mut)-pA; Pu6-crRNA-pT |
| **Sequence** | CTCGAGTAGTTATTAATAGTAATCAATTACGGGGTCATTAGTTCATAGCCCATATATGGAGTTCCGCGTTA<br>CATAACTTACGGTAAATGGCCCGCCTGGCTGACCGCCCAACGACCCCCGCCCATTGACGTCAATAATGACG<br>TATGTTCCCATAGTAACGCCAATAGGGACTTTCCATTGACGTCAATGGGTGGAGTATTTACGGTAAACTGC<br>CCACTTGGCAGTACATCAAGTGTATCATATGCCAAGTACGCCCCCTATTGACGTCAATGACGGTAAATGGC<br>CCGCCTGGCATTATGCCCAGTACATGACCTTATGGGACTTTCCTACTTGGCAGTACATCTACGTATTAGTC<br>ATCGCTATTACCATGGTGATGCGGTTTTGGCAGTACATCAATGGGCGTGGATAGCGGTTTGACTCACGGGG<br>ATTTCCAAGTCTCCACCCCATTGACGTCAATGGGAGTTTGTTTTGGCACCAAAATCAACGGGACTTTCCAA<br>AATGTCGTAACAACTCCGCCCCATTGACGCAAATGGGCGGTAGGCGTGTACGGTGGGAGGTCTATATAAGC<br>AGAGCTGGTTTAGTGAACCGTCAGATCCGCTAGGGATCCgccgccaccATGGATTACAAAGACGATGACGA<br>TAAGATGGCGCCTAAGAAGAAACGCAAAGTGCGGGGCATGAAAAATGACTACCGGACCTTCAAGCTGAGCC<br>TGCTGACCCTGGCTCCTATCCACATCGGCAACGGCGAGAAGTACACCAGCAGAGAATTCATCTACGAGAAC<br>AAGAAGTTCTACTTCCCCGACATGGGCAAGTTCTACAACAAGATGGTGGAAAAGAGACTGGCCGAGAAGTT<br>CGAGGCCTTCCTGATCCAGACCAGACCCAACGCCAGAAACAACCGGCTGATTTCTTTTCTGAACGACAACA<br>GAATCGCCGAAAGATCTTTTGGCGGCTACAGCATCAGTGAAACCGGCCTGGAATCTGATAAGAACCCTAAC<br>AGCGCCGGAGCTATCAACGAGGTGAACAAATTCATCCGGGACGCCTTCGGAAATCCTTACATCCCAGGCAG<br>CAGCCTGAAGGGCGCCATCCGCACCATCCTGATGAACACCACACCTAAGTGGAACAACGAGAACGCCGTGA<br>ACGACTTCGGCAGATTCCCAAAGGAAAACAAGAACCTGATCCCTTGGGGACCTAAGAAAGGCAAGGAATAC<br>GACGACCTGTTCAACGCCATCAGAGTGTCCGACAGCAAGCCCTTCGACAACAAAAGCCTGATCCTCGTGCA<br>GAAGTGGGACTACAGCGCCAAAACCAACAAGGCCAAGCCTCTGCCTCTGTACAGAGAGTCTATCAGCCCTC<br>TGACCAAGATCGAGTTCGAGATAACAACAACCACTGATGAGGCCGGCAGACTGATCGAGGAACTGGGAAAG<br>CGGGCCCAGGCCTTTTATAAGGACTACAAGGCCTTTTTCCTGTCTGAATTCCCTGATGATAAGATCCAGGC<br>TAATCTGCAATACCCCATCTACCTGGGCGCCGGCAGCGGCGCTTGGACAAAGACCCTGTTTAAGCAGGCCG<br>ACGGCATCCTGCAGCGGAGATACTCCAGAATGAAAACCAAGATGGTCAAGAAGGGCGTGCTGAAGCTGACA<br>AAGGCCCCTCTGAAAACAGTGAAGATCCCCAGCGGCAACCACAGCCTGGTGAAGAATCACGAGAGCTTCTA<br>CGAGATGGGCAAAGCCAACTTCATGATCAAGGAAATCGACAAGgaaggaaggggctccctcctcacttgtg<br>gagatgtcgaagaaaatcctggacctGATTACAAAGACGATGACGATAAGATGGCGCCTAAGAAGAAACGC<br>AAAGTGCGGGGCATGACTTACAAGCTCTACATTATGACCTTTCAAAACGCCCACTTCGGTTCCGGCACTCT<br>GGACTCATCGAAGCTGACCTTCTCCGCGGATAGAATCTTCTCGGCACTCGTGCTCGAGGCTCTGAAGATGG<br>GAAAGCTCGACGCCTTCTTGGCCGAGGCCAACCAGGATAAGTTCACTCTGACCGACGCGTTCCCATTCCAA<br>TTCGGTCCTTTCCTGCCGAAACCGATTGGTTACCCCAAGCACGACCAGATCGACCAGTCTGTGGACGTGAA<br>GGAAGTCCGCCGCCAAGCGAAGCTGTCCAAAAAGCTCCAGTTCCTGGCTCTGGAAAACGTCGACGACTACC<br>TGAACGGAGAGCTGTTTGAGAATGAGGAACACGCCGTGATCGACACAGTGACCAAGAACCAGCCCCATAAA<br>GATGATAATCTGTACCAAGTGGCCACCACTCGGTTCTCGAACGACACCTCCCTTTACGTGATCGCCAACGA<br>ATCCGATCTGCTGAACGAACTGATGAGCAGCCTTCAGTACTCCGGGCTGGGCGGCAAAAGGTCCTCAGGAT<br>TCGGCAGATTTGAGCTGGACATCCAGAACATTCCCTTGGAACTGTCCGACCGGCTGACGAAGAACCACAGC<br>GACAAGGTCATGTCACTTACCACCGCCCTCCCGGTGGACGCTGATCTCGAGGAAGCGATGGAAGATGGCCA<br>TTACCTGTTGACCAAGTCGTCCGGATTCGCATTCTCCCACGCCACCAACGAAAACTATCGGAAGCAGGACC<br>TGTACAAGTTCGCCTCCGGGAGCACCTTCAGCAAGACTTTCGAGGGACAGATCGTGGACGTGCGCCCTCTC<br>GATTTCCCTCACGCCGTGCTGAACTACGCCAAGCCGCTGTTCTTTAAGCTCGAAGTCgagggcagaggaag<br>tctgctaacatgcggtgacgtcgaggagaatcctggacctGATTACAAAGACGATGACGATAAGATGGCGC<br>CTAAGAAGAAACGCAAAGTGCGGGGCATGACCTTCGCCAAGATCAAATTCAGCGCCCAGATCCGGCTGGAA<br>ACCGGCCTGCACATCGGAGGATCTGATGCCTTTGCCGCTATCGGCGCCATCGACAGCCCTGTGATCAAGGA<br>CCCCATCACCAACCTGCCTATCATCCCCGGCTCTAGCCTGAAGGGCAAGATGAGAACACTGCTGGCCAAGG<br>TGTACAACGAAAAGGTGGCCGAGAAGCCTAGCGACGACAGCGACATCCTGAGCAGACTGTTCGGAAATAGC<br>AAGGATAAGCGGTTCAAGATGGGCAGACTGATCTTCCGGGACGCCTTCCTGAGCAACGCCGACGAGCTGGA<br>TTCTCTGGGCGTGCGGAGCTACACCGAGGTGAAGTTCGAGAACACCATCGATAGAATCACCGCCGAGGCCA<br>ATCCTAGACAGATCGAGAGAGCCATTCGGAACTCAACATTCGACTTCGAGCTGATCTACGAGATCACTGAT<br>GAGAATGAGAACCAGGTCGAGGAAGATTTCAAGGTGATCAGAGACGGCCTGAAGCTGCTGGAACTGGACTA<br>CCTGGGCGGAAGCGGCTCCAGAGGCTACGGCAAAGTGGCTTTTGAGAACCTGAAAGCCACCACAGTGTTCG<br>GCAACTACGACGTGAAAACCCTGAACGAGCTGCTGACCGCCGAAGTGgaggggcgggggtctttgttgact<br>tgcggggatgttgaggagaacccagggccaGATTACAAAGACGATGACGATAAGATGGCGCCTAAGAAGAA<br>ACGCAAAGTGCGGGGCATGACCATCCTGACCGACGAGAACTACGTGGACATCGCCGAGAAAGCCATCCTGA<br>AGCTGGAAAGAAACACCAGAAATAGAAAGAACCCTGATGCCTTCTTCCTGACCACATCTAAGCTGCGGAAC<br>CTGCTGAGCCTGACAAGCACCCTGTTCGACGAGAGCAAGGTGAAGGAATACGACGCCCTGCTGGACAGAAT<br>CGCTTATCTGAGAGTGCAGTTCGTGTACCAGGCCGGCAGAGAGATCGCCGTGAAAGATCTGATCGAGAAGG<br>CCCAGATCCTGGAAGCTCTGAAAGAGATCAAGGACCGGGAAACCCTGCAGAGATTCTGCAGATACATGGAA<br>GCCCTGGTGGCCTACTTCAAGTTCTACGGCGGCAAGGACTGAGCTAGCGCGGCCGCATCGATAAGCTTGTC |

Table S2

| | |
|---|---|
| | GACGATATCTCCAGAGGATCATAATCAGCCATACCACATTTGTAGAGGTTTTACTTGCTTTAAAAAACCTC<br>CCACACCTCCCCCTGAACCTGAAACATAAAATGAATGCAATTGTTGTTGTTAACTTGTTTATTGCAGCTTA<br>TAATGGTTACAAATAAAGCAATAGCATCACAAATTTCACAAATAAAGCATTTTTTTCACTGCCCCGAGCTT<br>CCTCGCTCACTGACTCGCTGCGCTCGGTCGTTCGGCTGCGGCGAGCGGTATCAGCTCACTCAAAGGCGGTA<br>ATACGGTTATCCACAGAATCAGGGGATAACGCAGGAAAGAAACTAGTGAGGGCCTATTTCCCATGATTCCT<br>TCATATTTGCATATACGATACAAGGCTGTTAGAGAGATAATTAGAATTAATTTGACTGTAAACACAAAGAT<br>ATTAGTACAAAATACGTGACGTAGAAAGTAATAATTTCTTGGGTAGTTTGCAGTTTTAAAATTATGTTTTA<br>AAATGGACTATCATATGCTTACCGTAACTTGAAAGTATTTCGATTTCTTGGCTTTATATATCTTGTGGAAA<br>GGACGAAACACCGATATAAACCTAATTACCTCGAGAGGGGACGGAAACCCGTCTTCGATGAAGCGATTCAG<br>AAGACTTGATATAAACCTAATTACCTCGAGAGGGGACTTTTTTACATGTGAGCAAAAGGCCAGCAAAAGGC<br>CAGGAACCGTAAAAAGGCCGCGTTGCTGGCGTTTTTCCATAGGCTCCGCCCCCCTGACGAGCATCACAAAA<br>ATCGACGCTCAAGTCAGAGGTGGCGAAACCCGACAGGACTATAAAGATACCAGGCGTTTCCCCCTGGAAGC<br>TCCCTCGTGCGCTCTCCTGTTCCGACCCTGCCGCTTACCGGATACCTGTCCGCCTTTCTCCCTTCGGGAAG<br>CGTGGCGCTTTCTCATAGCTCACGCTGTAGGTATCTCAGTTCGGTGTAGGTCGTTCGCTCCAAGCTGGGCT<br>GTGTGCACGAACCCCCCGTTCAGCCCGACCGCTGCGCCTTATCCGGTAACTATCGTCTTGAGTCCAACCCG<br>GTAAGACACGACTTATCGCCACTGGCAGCAGCCACTGGTAACAGGATTAGCAGAGCGAGGTATGTAGGCGG<br>TGCTACAGAGTTCTTGAAGTGGTGGCCTAACTACGGCTACACTAGAAGAACAGTATTTGGTATCTGCGCTC<br>TGCTGAAGCCAGTTACCTTCGGAAAAAGAGTTGGTAGCTCTTGATCCGGCAAACAAACCACCGCTGGTAGC<br>GGTGGTTTTTTTGTTTGCAAGCAGCAGATTACGCGCAGAAAAAAAGGATCTCAAGAAGATCCTTTGATCTT<br>TTCTACGGGGTCTGACGCTCAGTGGAACGAAAACTCACGTTAAGGGATTTTGGTCATGAGATTATCAAAAA<br>GGATCTTCACCTAGATCCTTTTAAATTAAAAATGAAGTTTTAAATCAATCTAAAGTATATATGAGTAAACT<br>TGGTCTGACAGTTACCAATGCTTAATCAGTGAGGCACCTATCTCAGCGATCTGTCTATTTCGTTCATCCAT<br>AGTTGCCTGACTCCCCGTCGTGTAGATAACTACGATACGGGAGGGCTTACCATCTGGCCCCAGTGCTGCAA<br>TGATACCGCGAGACCCACGCTCACCGGCTCCAGATTTATCAGCAATAAACCAGCCAGCCGGAAGGGCCGAG<br>CGCAGAAGTGGTCCTGCAACTTTATCCGCCTCCATCCAGTCTATTAATTGTTGCCGGGAAGCTAGAGTAAG<br>TAGTTCGCCAGTTAATAGTTTGCGCAACGTTGTTGCCATTGCTACAGGCATCGTGGTGTCACGCTCGTCGT<br>TTGGTATGGCTTCATTCAGCTCCGGTTCCCAACGATCAAGGCGAGTTACATGATCCCCCATGTTGTGCAAA<br>AAAGCGGTTAGCTCCTTCGGTCCTCCGATCGTTGTCAGAAGTAAGTTGGCCGCAGTGTTATCACTCATGGT<br>TATGGCAGCACTGCATAATTCTCTTACTGTCATGCCATCCGTAAGATGCTTTTCTGTGACTGGTGAGTACT<br>CAACCAAGTCATTCTGAGAATAGTGTATGCGGCGACCGAGTTGCTCTTGCCCGGCGTCAACACGGGATAAT<br>ACCGCGCCACATAGCAGAACTTTAAAAGTGCTCATCATTGGAAAACGTTCTTCGGGGCGAAAACTCTCAAG<br>GATCTTACCGCTGTTGAGATCCAGTTCGATGTAACCCACTCGTGCACCCAACTGATCTTCAGCATCTTTTA<br>CTTTCACCAGCGTTTCTGGGTGAGCAAAAACAGGAAGGCAAAATGCCGCAAAAAAGGGAATAAGGGCGACA<br>CGGAAATGTTGAATACTCATACTCTTCCTTTTTCAATATTATTGAAGCATTTATCAGGGTTATTGTCTCAT<br>GAGCGGATACATATTTGAATGTATTTAGAAAAATAAACAAATAGGGGTTCCGCGCACATTTCCCCGAAAAG<br>TGCCACCTGACGTCGGCAGTGAAAAAAATGCTTTATTTGTGAAATTTGTGATGCTATTGCTTTATTTGTAA<br>CCATTATAAGCTGCAATAAACAAGTTAACAACAACAATTGCATTCATTTTATGTTTCAGGTTCAGGGGGAG<br>GTGTGGGAGGTTTTTTAAAGCAAGTAAAACCTCTACAAATGTGGTATGGCTGATTATGATCCTCTAGATTA<br>ATCCTTTCTGATTTCGTAGATGTACAGCAGGAGCGCGAGCTCGGCTTCCTTCCGGTCCTTATCGTTCTTGT<br>TGGTGTACCAGGAGTAGAACAGGTTCTTGAAGGTCTTGAACTTGTCCCTGTCAGTCTCCCGGGTCAGTTCT<br>TCCAGGCGAGTGAGGTAATAGGCGAGGCGTGCCATATTCATGCGGTCGTGGTTCCTCAAAAGCTCAATAAG<br>CTTATAGATGAAGTTCTTTCCCCTCTCGTCTTGATGGTTGAAGAAGTATCTAATCTGTTCCAGTTTGTCGT<br>CGTACACGTTAGTGATGAACCTATCAAACTTGAAAGTGTAGTCGCTTGAGAACAGCGAGATGGAGTCCTTT<br>TCGTTGCCCTTGGCGGCCTCTTCCAGTTCCCCGGTCTGGTGAGCCATCAGGCTAATAGGAGTCTTGTCGGC<br>GAACAACCCTATCCCCGCGGAGAGGGTGAGCTTCCCGTTGGTCCACTTGATGAAGTTTTCGCGAAGTTCCA<br>CAGTGAACGCGATGATATCCTGCCACGATCCAATGGCGAACACGGCGGCGCCGCCCGCGTAGATAATGCTC<br>AGCTTCTTGTCGGAGGCGAACTGGTTAATGTACACTTTGAAGAACAGCGACATGCTCCGGGAGAATGTGGC<br>CGATCTTGACAGAGTGGAGTATTGTCCGTTTCCCTGCTGGCTGAAACCGGCCATGAAGGCGGCGCCCAAGT<br>CATCCACGTCGAGCCGGACCACGGCCAGTCTCTTGATGCCTAGGCCGTTCTCGTTCTTGCTCAGGGCGGCG<br>TAGTTGTAGATCTCGTCGCACTGGTAATCCCCCACGAACACATGCGTAGCCTTCACGGTACCGGCCTTATA<br>GTCATTCTTCACGTAGACCCGGCTGAACGCTTCTTGGGACAGCTTTTCGAATGCCACGCCCTTTAAGCACG<br>CGTTTGGTCCAATCGGCAGCCCCTCATTTTCGGTAATGATGAAGTGGTCATGGGCAATTTCTTTCGAGAAC<br>TGGTACAGTCCCCGGCAAATGTCACAGACTTTCTGGTCGTGGTAGGACACCAGGTTCTCCACGGAGTGGCA<br>AATCTCGCACTCTCTCTCTGAGGACTTTCCACCGCGATTCAGGAGCATCAGTGTCTGGTAGTCGTATCTGG<br>AGATTTTCTTTTTGGAGATCATGCGCGAAGCCTTTTGGTACACTTGGCGGTAGGACTCGGGGCTATTCAGC<br>TCGGACATGATGTCCTTGGCCGCGAAGGAACCCCAGCCAAAGGCCACATAGAGGCGGGTCTGGAAGTTTGC<br>CAACAGGAACTGGTTGAAATCCTTCTCAAACTGCACCAGGGTTTCCACAGTCTTTTCGGTGTTGGCCAGGA<br>CGAAGTAGGCGTGTCCGCCGCCGACGTAAAGCATGTTAGCCCTGTTCAGTCCCAGCTTGTCCAGCAGGCTA<br>TCGGCGATGTACTCGGACATAAAGTCCAGGTAGAGGCTCCGGGCCTTCAGCTGCTTCGCCGCGCCGTTAGT<br>TGCGATGTTAATGTTGTAGATAAAGTCCTGGATTCCCGACAGGTCGAAGGAGGCCAGCAGGAAGGCTTCTT<br>CTTCATAGAACGCTGACACTTTGGTGAACAGGTCCTCTTTGTAGTTGTGCCGACCCTTGTCCTCCAGGTAG<br>TCGTAGATCGCCAGAGCGAAGGCAGCAGTCAAGCGGGAATGGTCGGCCAGGGAGATGTCGGCGATTTCCTT<br>AGTGTTGGTGCTTGACGGCACGAAGGAGAGAGTAGCCTCGAAAAGGTTGAGCAGGGAGTCAATCTGGACTT |

Table S2

| | |
|---|---|
| | GGTTGAACTCGAACTCGGCCAGTTCGTTCTTAATCCGGGTGGCGATGGCAGCGTAATCGCCCTTGCTAAAG<br>GGTTCGTAAGTGGCGGACGCGAAGTTGGGCTTCGACTTGAGATTCAGCACGGTAGGCTTGAAGTACCGCTT<br>ATCGGTCTGCGCTCCGAACACGTTAAAGATGTCGGCCTGGTTCGTGTAGGTGTCCCAGATCTTTGCGGAGG<br>TATCTTCGTCTGACTCTTCGTTGGATTGCCGGCGGTCGACACCGGAGGCGATGTTGTCGGCAATGTAGGTG<br>ATGTAAGCCAGGTGATCGTTGCCGAGCTTATCAGACTGGTAGTTGGCCATATGGTACCGGATCTGATCCGA<br>GATGACTTGGTTGTCGGCGATCTCGTCGAACCAGTCGGCGCCCACAAGTGCGTGTTTCTTCCGCTCTCCGG<br>TTGCTCGCTGGATGACCTTTCCGATGGCGGCCAGCAGGGCTCCGTAAAACAGATCAATCTTTTCTTTCTTC<br>ATGCCCCGCACTTTGCGTTTCTTCTTAGGCGCCATCTTATCGTCATCGTCTTTGTAATCGCCGGAATGGTG<br>ATGGTGATGGTGtggtcctggattttcttcaacatctccacaagttagcaaacttcctcttccttcGTCCT<br>TTCTTTCTTCCAGCTTGATGCCGCCCATTCCCAGGCTGGTTTTCATGCCCAGGCCGCTGTACTCGCCGAAG<br>GTCAGCAGCATCTTCACGTAGGCCTTCAGTGTCTGGGCTCCCTGCACCTTGAAGGTCAGCTTGCCCCGGAA<br>GGCGGGGATTCTCTGTCTATGCACTCTGAAGTAGCTTGTTTCCAGTCTGTAATTGGTGATGGTGCTGTGCT<br>CGCTCAGGTAGTCCAGGGTATCTTCCTCGATCTCAGGCTGGTTTTCCACCAGTCTGCCGTACTTCTGCATC<br>AGGCTCTGAAAGATCAGCCGCATGCTGGGGAAGATCACGTAGCTGCCCTGGCTTTTAAAGCCTGTAGGTGT<br>CTCGAAGATGATGCTGAAGTAGGTCTGGTTTTCCTCGCTGTTGAAGATCTCGAGCAGCTTATCAGCGCCCA<br>GCTCCTGGATTTCCACCTTTTCCACAGACAACTTAGGCAGGTCGTTCAGGGACACTTCCTTGATCTGCAGC<br>AGGGTCATGAACACCTTGTCCTCGATGTCGTCTGTCAGCAGATGCACGACCCACTGGGTGTTCTCTTTGCC<br>CTGGATCACCTTTGTGGCGTAGGGGTTGGTCTGCTGCTGGTGCAGGTAGTCCACGTAGTCGCTATCCAGCT<br>GTTCCATCAGGAAGCCGTGGAATTTCACGGCCAGGTCCTGAGCAGGGTGGTCGATCCGCTTAAAGGTGAAC<br>ACGAGCTTTTTCATGCCCCGCACTTTGCGTTTCTTCTTAGGCGCCATCTTATCGTCATCGTCTTTGTAATC<br>ggggccggggttctcctccacgtcgccgcaggtcagcaggctgcctctgccctcCTTGTACAGCTCGTCCA<br>TGCCGCCGGTGGAGTGGCGGCCCTCGGCGCGTTCGTACTGTTCCACGATGGTGTAGTCCTCGTTGTGGGAG<br>GTGATGTCCAACTTGATGTTGACGTTGTAGGCGCCGGGCAGCTGCACGGGCTTCTTGGCCTTGTAGGTGGT<br>CTTGACCTCAGCGTCGTAGTGGCCGCCGTCCTTCAGCTTCAGCCTCTGCTTGATCTCGCCCTTCAGGGCGC<br>CGTCCTCGGGGTACATCCGCTCGGAGGAGGCCTCCCAGCCCATGGTTTTCTTCTGCATTACGGGGCCGTCG<br>GAGGGGAAGTTGGTGCCGCGCAGCTTCACCTTGTAGATGAACTCGCCGTCCTGCAGGGAGGAGTCCTGGGT<br>CACGGTCACCACGCCGCCGTCCTCGAAGTTCATCACGCGCTCCCACTTGAAGCCCTCGGGGAAGGACAGCT<br>TCAAGTAGTCGGGGATGTCGGCGGGGTGCTTCACGTAGGCCTTGGAGCCGTACATGAACTGAGGGGACAGG<br>ATGTCCCAGGCGAAGGGCAGGGGGCCACCCTTGGTCACCTTCAGCTTGGCGGTCTGGGTGCCCTCGTAGGG<br>GCGGCCCTCGCCCTCGCCCTCGATCTCGAACTCGTGGCCGTTCACGGAGCCCTCCATGTGCACCTTGAAGC<br>GCATGAACTCCTTGATGATGGCCATGTTATCCTCCTCGCCCTTGCTCACCATggtggcggcACCGGTGAAT<br>TCTCCAGGCGATCTGACGGTTCACTAAACGAGCTCTGCTTATATAGGCCTCCCACCGTACACGCCACCTCG<br>ACATA |

| **Plasmid** | **pDAC742** |
|---|---|
| **Description** | Expression of Csm complex (DNase/cA mut) from separate promoters; RFP backbone |
| **Utility** | RNA KD |
| **Features** | Pcmv-FLAG-NLS-Csm1(DNase/cA mut)-pA; Pcmv-FLAG-NLS-Csm2-pA, Pcmv-FLAG-NLS-Csm3-pA; Pcmv-FLAG-NLS-Csm4-pA; Pcmv-FLAG-NLS-Csm5-pA; Pcmv-FLAG-NLS-Cas6-pA; Pu6-crRNA-pT; Pcmv-RFP-pA |
| **Sequence** | GTGATGCGGTTTTGGCAGTACATCAATGGGCGTGGATAGCGGTTTGACTCACGGGGATTTCCAAGTCTCCA<br>CCCCATTGACGTCAATGGGAGTTTGTTTTGGCACCAAAATCAACGGGACTTTCCAAAATGTCGTAACAACT<br>CCGCCCCATTGACGCAAATGGGCGGTAGGCGTGTACGGTGGGAGGTCTATATAAGCAGAGCTCGTTTAGTG<br>AACCGTCAGATCTCTAGAgccgccaccATGCACCATCACCATCACCATTCCGGCGATTACAAAGACGATGA<br>CGATAAGATGGCGCCTAAGAAGAAACGCAAAGTGCGGGGCATGAAGAAAGAAAAGATTGATCTGTTTTACG<br>GAGCCCTGCTGGCCGCCATCGGAAAGGTCATCCAGCGAGCAACCGGAGAGCGGAAGAAACACGCACTTGTG<br>GGCGCCGACTGGTTCGACGAGATCGCCGACAACCAAGTCATCTCGGATCAGATCCGGTACCATATGGCCAA<br>CTACCAGTCTGATAAGCTCGGCAACGATCACCTGGCTTACATCACCTACATTGCCGACAACATCGCCTCCG<br>GTGTCGACCGCCGGCAATCCAACGAAGAGTCAGACGAAGATACCTCCGCAAAGATCTGGGACACCTACACG<br>AACCAGGCCGACATCTTTAACGTGTTCGGAGCGCAGACCGATAAGCGGTACTTCAAGCCTACCGTGCTGAA<br>TCTCAAGTCGAAGCCCAACTTCGCGTCCGCCACTTACGAACCCTTTAGCAAGGGCGATTACGCTGCCATCG<br>CCACCCGGATTAAGAACGAACTGGCCGAGTTCGAGTTCAACCAAGTCCAGATTGACTCCCTGCTCAACCTT<br>TTCGAGGCTACTCTCTCCTTCGTGCCGTCAAGCACCAACACTAAGGAAATCGCCGACATCTCCCTGGCCGA<br>CCATTCCCGCTTGACTGCTGCCTTCGCTCTGGCGATCTACGACTACCTGGAGGACAAGGGTCGGCACAACT<br>ACAAAGAGGACCTGTTCACCAAAGTGTCAGCGTTCTATGAAGAAGAAGCCTTCCTGCTGGCCTCCTTCGAC<br>CTGTCGGGAATCCAGGACTTTATCTACAACATTAACATCGCAACTAACGGCGCGGCGAAGCAGCTGAAGGC<br>CCGGAGCCTCTACCTGGACTTTATGTCCGAGTACATCGCCGATAGCCTGCTGGACAAGCTGGGACTGAACA<br>GGGCTAACATGCTTTACGTCGGCGGCGGACACGCCTACTTCGTCCTGGCCAACACCGAAAAGACTGTGGAA<br>ACCCTGGTGCAGTTTGAGAAGGATTTCAACCAGTTCCTGTTGGCAAACTTCCAGACCCGCCTCTATGTGGC<br>CTTTGGCTGGGGTTCCTTCGCGGCCAAGGACATCATGTCCGAGCTGAATAGCCCCGAGTCCTACCGCCAAG<br>TGTACCAAAAGGCTTCGCGCATGATCTCCAAAAAGAAAATCTCCAGATACGACTACCAGACACTGATGCTC<br>CTGAATCGCGGTGGAAAGTCCTCAGAGAGAGAGTGCGAGATTTGCCACTCCGTGGAGAACCTGGTGTCCTA |

Table S2

| | |
|---|---|
| | CCACGACCAGAAAGTCTGTGACATTTGCCGGGGACTGTACCAGTTCTCGAAAGAAATTGCCCATGACCACT<br>TCATCATTACCGAAAATGAGGGGCTGCCGATTGGACCAAACGCGTGCTTAAAGGGCGTGGCATTCGAAAAG<br>CTGTCCCAAGAAGCGTTCAGCCGGGTCTACGTGAAGAATGACTATAAGGCCGGTACCGTGAAGGCTACGCA<br>TGTGTTCGTGGGGGATTACCAGTGCGACGAGATCTACAACTACGCCGCCCTGAGCAAGAACGAGAACGGCC<br>TAGGCATCAAGAGACTGGCCGTGGTCCGGCTCGACGTGGATGACTTGGGCGCCGCCTTCATGGCCGGTTTC<br>AGCCAGCAGGGAAACGGACAATACTCCACTCTGTCAAGATCGGCCACATTCTCCCGGAGCATGTCGCTGTT<br>CTTCAAAGTGTACATTAACCAGTTCGCCTCCGACAAGAAGCTGAGCATTATCTACGCGGGCGGCGCCGCCG<br>TGTTCGCCATTGGATCGTGGCAGGATATCATCGCGTTCACTGTGGAACTTCGCGAAAACTTCATCAAGTGG<br>ACCAACGGGAAGCTCACCCTCTCCGCGGGGATAGGGTTGTTCGCCGACAAGACTCCTATTAGCCTGATGGC<br>TCACCAGACCGGGGAACTGGAAGAGGCCGCCAAGGGCAACGAAAAGGACTCCATCTCGCTGTTCTCAAGCG<br>ACTACACTTTCAAGTTTGATAGGTTCATCACTAACGTGTACGACGACAAACTGGAACAGATTAGATACTTC<br>TTCAACCATCAAGACGAGAGGGGAAAGAACTTCATCTATAAGCTTATTGAGCTTTTGAGGAACCACGACCG<br>CATGAATATGGCACGCCTCGCCTATTACCTCACTCGCCTGGAAGAACTGACCCGGGAGACTGACAGGGACA<br>AGTTCAAGACCTTCAAGAACCTGTTCTACTCCTGGTACACCAACAAGAACGATAAGGACCGGAAGGAAGCC<br>GAGCTCGCGCTCCTGCTGTACATCTACGAAATCAGAAAGGATTAAcggcaataaaaagacagaataaaacg<br>cacggtgttgggtcgtttgttcGCACACATTAGCTAGCCGTCAGCACACATTGTGATGCGGTTTTGGCAGT<br>ACATCAATGGGCGTGGATAGCGGTTTGACTCACGGGGATTTCCAAGTCTCCACCCCATTGACGTCAATGGG<br>AGTTTGTTTTGGCACCAAAATCAACGGGACTTTCCAAAATGTCGTAACAACTCCGCCCCATTGACGCAAAT<br>GGGCGGTAGGCGTGTACGGTGGGAGGTCTATATAAGCAGAGCTCGTTTAGTGAACCGTCAGATCTCTAGAg<br>ccgccaccATGGATTACAAAGACGATGACGATAAGATGGCGCCTAAGAAGAAACGCAAAGTGCGGGGCATG<br>ACCATCCTGACCGACGAGAACTACGTGGACATCGCCGAGAAAGCCATCCTGAAGCTGGAAAGAAACACCAG<br>AAATAGAAAGAACCCTGATGCCTTCTTCCTGACCACATCTAAGCTGCGGAACCTGCTGAGCCTGACAAGCA<br>CCCTGTTCGACGAGAGCAAGGTGAAGGAATACGACGCCCTGCTGGACAGAATCGCTTATCTGAGAGTGCAG<br>TTCGTGTACCAGGCCGGCAGAGAGATCGCCGTGAAAGATCTGATCGAGAAGGCCCAGATCCTGGAAGCTCT<br>GAAAGAGATCAAGGACCGGGAAACCCTGCAGAGATTCTGCAGATACATGGAAGCCCTGGTGGCCTACTTCA<br>AGTTCTACGGCGGCAAGGACTGAcggcaataaaaagacagaataaaacgcacggtgttgggtcgtttgttc<br>AGTTCTTTTGCCTTACTTTCAATGCATGCGGTGATGCGGTTTTGGCAGTACATCAATGGGCGTGGATAGCG<br>GTTTGACTCACGGGGATTTCCAAGTCTCCACCCCATTGACGTCAATGGGAGTTTGTTTTGGCACCAAAATC<br>AACGGGACTTTCCAAAATGTCGTAACAACTCCGCCCCATTGACGCAAATGGGCGGTAGGCGTGTACGGTGG<br>GAGGTCTATATAAGCAGAGCTCGTTTAGTGAACCGTCAGATCTCTAGAgccgccaccATGGATTACAAAGA<br>CGATGACGATAAGATGGCGCCTAAGAAGAAACGCAAAGTGCGGGGCATGACCTTCGCCAAGATCAAATTCA<br>GCGCCCAGATCCGGCTGGAAACCGGCCTGCACATCGGAGGATCTGATGCCTTTGCCGCTATCGGCGCCATC<br>GACAGCCCTGTGATCAAGGACCCCATCACCAACCTGCCTATCATCCCCGGCTCTAGCCTGAAGGGCAAGAT<br>GAGAACACTGCTGGCCAAGGTGTACAACGAAAAGGTGGCCGAGAAGCCTAGCGACGACAGCGACATCCTGA<br>GCAGACTGTTCGGAAATAGCAAGGATAAGCGGTTCAAGATGGGCAGACTGATCTTCCGGGACGCCTTCCTG<br>AGCAACGCCGACGAGCTGGATTCTCTGGGCGTGCGGAGCTACACCGAGGTGAAGTTCGAGAACACCATCGA<br>TAGAATCACCGCCGAGGCCAATCCTAGACAGATCGAGAGAGCCATTCGGAACTCAACATTCGACTTCGAGC<br>TGATCTACGAGATCACTGATGAGAATGAGAACCAGGTCGAGGAAGATTTCAAGGTGATCAGAGACGGCCTG<br>AAGCTGCTGGAACTGGACTACCTGGGCGGAAGCGGCTCCAGAGGCTACGGCAAAGTGGCTTTTGAGAACCT<br>GAAAGCCACCACAGTGTTCGGCAACTACGACGTGAAAACCCTGAACGAGCTGCTGACCGCCGAAGTGTGAc<br>ggcaataaaaagacagaataaaacgcacggtgttgggtcgtttgttcTGGATTGCGAGAATGGACTAGTAG<br>CAAACTGTGATGCGGTTTTGGCAGTACATCAATGGGCGTGGATAGCGGTTTGACTCACGGGGATTTCCAAG<br>TCTCCACCCCATTGACGTCAATGGGAGTTTGTTTTGGCACCAAAATCAACGGGACTTTCCAAAATGTCGTA<br>ACAACTCCGCCCCATTGACGCAAATGGGCGGTAGGCGTGTACGGTGGGAGGTCTATATAAGCAGAGCTCGT<br>TTAGTGAACCGTCAGATCTCTAGAgccgccaccATGGATTACAAAGACGATGACGATAAGATGGCGCCTAA<br>GAAGAAACGCAAAGTGCGGGGCATGACTTACAAGCTCTACATTATGACCTTTCAAAACGCCCACTTCGGTT<br>CCGGCACTCTGGACTCATCGAAGCTGACCTTCTCCGCGGATAGAATCTTCTCGGCACTCGTGCTCGAGGCT<br>CTGAAGATGGGAAAGCTCGACGCCTTCTTGGCCGAGGCCAACCAGGATAAGTTCACTCTGACCGACGCGTT<br>CCCATTCCAATTCGGTCCTTTCCTGCCGAAACCGATTGGTTACCCCAAGCACGACCAGATCGACCAGTCTG<br>TGGACGTGAAGGAAGTCCGCCGCCAAGCGAAGCTGTCCAAAAAGCTCCAGTTCCTGGCTCTGGAAAACGTC<br>GACGACTACCTGAACGGAGAGCTGTTTGAGAATGAGGAACACGCCGTGATCGACACAGTGACCAAGAACCA<br>GCCCCATAAAGATGATAATCTGTACCAAGTGGCCACCACTCGGTTCTCGAACGACACCTCCCTTTACGTGA<br>TCGCCAACGAATCCGATCTGCTGAACGAACTGATGAGCAGCCTTCAGTACTCCGGGCTGGGCGGCAAAAGG<br>TCCTCAGGATTCGGCAGATTTGAGCTGGACATCCAGAACATTCCCTTGGAACTGTCCGACCGGCTGACGAA<br>GAACCACAGCGACAAGGTCATGTCACTTACCACCGCCCTCCCGGTGGACGCTGATCTCGAGGAAGCGATGG<br>AAGATGGCCATTACCTGTTGACCAAGTCGTCCGGATTCGCATTCTCCCACGCCACCAACGAAAACTATCGG<br>AAGCAGGACCTGTACAAGTTCGCCTCCGGGAGCACCTTCAGCAAGACTTTCGAGGGACAGATCGTGGACGT<br>GCGCCCTCTCGATTTCCCTCACGCCGTGCTGAACTACGCCAAGCCGCTGTTCTTTAAGCTCGAAGTCTAAc<br>ggcaataaaaagacagaataaaacgcacggtgttgggtcgtttgttcGCACATTCAAAAACAGGCAATTGG<br>ACAAGCGTGATGCGGTTTTGGCAGTACATCAATGGGCGTGGATAGCGGTTTGACTCACGGGGATTTCCAAG<br>TCTCCACCCCATTGACGTCAATGGGAGTTTGTTTTGGCACCAAAATCAACGGGACTTTCCAAAATGTCGTA<br>ACAACTCCGCCCCATTGACGCAAATGGGCGGTAGGCGTGTACGGTGGGAGGTCTATATAAGCAGAGCTCGT<br>TTAGTGAACCGTCAGATCTCTAGAgccgccaccATGGATTACAAAGACGATGACGATAAGATGGCGCCTAA |

Table S2

| | |
|---|---|
| | GAAGAAACGCAAAGTGCGGGGCATGAAAAATGACTACCGGACCTTCAAGCTGAGCCTGCTGACCCTGGCTC<br>CTATCCACATCGGCAACGGCGAGAAGTACACCAGCAGAGAATTCATCTACGAGAACAAGAAGTTCTACTTC<br>CCCGACATGGGCAAGTTCTACAACAAGATGGTGGAAAAGAGACTGGCCGAGAAGTTCGAGGCCTTCCTGAT<br>CCAGACCAGACCCAACGCCAGAAACAACCGGCTGATTTCTTTTCTGAACGACAACAGAATCGCCGAAAGAT<br>CTTTTGGCGGCTACAGCATCAGTGAAACCGGCCTGGAATCTGATAAGAACCCTAACAGCGCCGGAGCTATC<br>AACGAGGTGAACAAATTCATCCGGGACGCCTTCGGAAATCCTTACATCCCAGGCAGCAGCCTGAAGGGCGC<br>CATCCGCACCATCCTGATGAACACCACACCTAAGTGGAACAACGAGAACGCCGTGAACGACTTCGGCAGAT<br>TCCCAAAGGAAAACAAGAACCTGATCCCTTGGGGACCTAAGAAAGGCAAGGAATACGACGACCTGTTCAAC<br>GCCATCAGAGTGTCCGACAGCAAGCCCTTCGACAACAAAAGCCTGATCCTCGTGCAGAAGTGGGACTACAG<br>CGCCAAAACCAACAAGGCCAAGCCTCTGCCTCTGTACAGAGAGTCTATCAGCCCTCTGACCAAGATCGAGT<br>TCGAGATAACAACAACCACTGATGAGGCCGGCAGACTGATCGAGGAACTGGGAAAGCGGGCCCAGGCCTTT<br>TATAAGGACTACAAGGCCTTTTTCCTGTCTGAATTCCCTGATGATAAGATCCAGGCTAATCTGCAATACCC<br>CATCTACCTGGGCGCCGGCAGCGGCGCTTGGACAAAGACCCTGTTTAAGCAGGCCGACGGCATCCTGCAGC<br>GGAGATACTCCAGAATGAAAACCAAGATGGTCAAGAAGGGCGTGCTGAAGCTGACAAAGGCCCCTCTGAAA<br>ACAGTGAAGATCCCCAGCGGCAACCACAGCCTGGTGAAGAATCACGAGAGCTTCTACGAGATGGGCAAAGC<br>CAACTTCATGATCAAGGAAATCGACAAGTGAcggcaataaaaagacagaataaaacgcacggtgttgggtc<br>gtttgttcATGTGTGCCGGCCGGAGAATAAAAGTCTAAGTGATGCGGTTTTGGCAGTACATCAATGGGCGT<br>GGATAGCGGTTTGACTCACGGGGATTTCCAAGTCTCCACCCCATTGACGTCAATGGGAGTTTGTTTTGGCA<br>CCAAAATCAACGGGACTTTCCAAAATGTCGTAACAACTCCGCCCCATTGACGCAAATGGGCGGTAGGCGTG<br>TACGGTGGGAGGTCTATATAAGCAGAGCTCGTTTAGTGAACCGTCAGATCTCTAGAgccgccaccATGGAT<br>TACAAAGACGATGACGATAAGATGGCGCCTAAGAAGAAACGCAAAGTGCGGGGCATGAAAAAGCTCGTGTT<br>CACCTTTAAGCGGATCGACCACCCTGCTCAGGACCTGGCCGTGAAATTCCACGGCTTCCTGATGGAACAGC<br>TGGATAGCGACTACGTGGACTACCTGCACCAGCAGCAGACCAACCCCTACGCCACAAAGGTGATCCAGGGC<br>AAAGAGAACACCCAGTGGGTCGTGCATCTGCTGACAGACGACATCGAGGACAAGGTGTTCATGACCCTGCT<br>GCAGATCAAGGAAGTGTCCCTGAACGACCTGCCTAAGTTGTCTGTGGAAAAGGTGGAAATCCAGGAGCTGG<br>GCGCTGATAAGCTGCTCGAGATCTTCAACAGCGAGGAAAACCAGACCTACTTCAGCATCATCTTCGAGACA<br>CCTACAGGCTTTAAAAGCCAGGGCAGCTACGTGATCTTCCCCAGCATGCGGCTGATCTTTCAGAGCCTGAT<br>GCAGAAGTACGGCAGACTGGTGGAAAACCAGCCTGAGATCGAGGAAGATACCCTGGACTACCTGAGCGAGC<br>ACAGCACCATCACCAATTACAGACTGGAAACAAGCTACTTCAGAGTGCATAGACAGAGAATCCCCGCCTTC<br>CGGGGCAAGCTGACCTTCAAGGTGCAGGGAGCCCAGACACTGAAGGCCTACGTGAAGATGCTGCTGACCTT<br>CGGCGAGTACAGCGGCCTGGGCATGAAAACCAGCCTGGGAATGGGCGGCATCAAGCTGGAAGAAAGAAAGG<br>ACTGAcggcaataaaaagacagaataaaacgcacggtgttgggtcgtttgttcGTGCGATAGAGGGATCCC<br>GCATTGAATTATGTGATGCGGTTTTGGCAGTACATCAATGGGCGTGGATAGCGGTTTGACTCACGGGGATT<br>TCCAAGTCTCCACCCCATTGACGTCAATGGGAGTTTGTTTTGGCACCAAAATCAACGGGACTTTCCAAAAT<br>GTCGTAACAACTCCGCCCCATTGACGCAAATGGGCGGTAGGCGTGTACGGTGGGAGGTCTATATAAGCAGA<br>GCTCGTTTAGTGAACCGTCAGATCTCTAGAgccgccaccATGGTGAGCAAGGGCGAGGAGGATAACATGGC<br>CATCATCAAGGAGTTCATGCGCTTCAAGGTGCACATGGAGGGCTCCGTGAACGGCCACGAGTTCGAGATCG<br>AGGGCGAGGGCGAGGGCCGCCCCTACGAGGGCACCCAGACCGCCAAGCTGAAGGTGACCAAGGGTGGCCCC<br>CTGCCCTTCGCCTGGGACATCCTGTCCCCTCAGTTCATGTACGGCTCCAAGGCCTACGTGAAGCACCCCGC<br>CGACATCCCCGACTACTTGAAGCTGTCCTTCCCCGAGGGCTTCAAGTGGGAGCGCGTGATGAACTTCGAGG<br>ACGGCGGCGTGGTGACCGTGACCCAGGACTCCTCCCTGCAGGACGGCGAGTTCATCTACAAGGTGAAGCTG<br>CGCGGCACCAACTTCCCCTCCGACGGCCCCGTAATGCAGAAGAAAACCATGGGCTGGGAGGCCTCCTCCGA<br>GCGGATGTACCCCGAGGACGGCGCCCTGAAGGGCGAGATCAAGCAGAGGCTGAAGCTGAAGGACGGCGGCC<br>ACTACGACGCTGAGGTCAAGACCACCTACAAGGCCAAGAAGCCCGTGCAGCTGCCCGGCGCCTACAACGTC<br>AACATCAAGTTGGACATCACCTCCCACAACGAGGACTACACCATCGTGGAACAGTACGAACGCGCCGAGGG<br>CCGCCACTCCACCGGCGGCATGGACGAGCTGTACAAGTAAcggcaataaaaagacagaataaaacgcacgg<br>tgttgggtcgtttgttcGAGCAGATTGTACTGAGAGTGCACCGGTTGGAGGGCCTATTTCCCATGATTCCT<br>TCATATTTGCATATACGATACAAGGCTGTTAGAGAGATAATTAGAATTAATTTGACTGTAAACACAAAGAT<br>ATTAGTACAAAATACGTGACGTAGAAAGTAATAATTTCTTGGGTAGTTTGCAGTTTTAAAATTATGTTTTA<br>AAATGGACTATCATATGCTTACCGTAACTTGAAAGTATTTCGATTTCTTGGCTTTATATATCTTGTGGAAA<br>GGACGAAACACCGATATAAACCTAATTACCTCGAGAGGGGACGGAAACCCGTCTTCGATGAAGCGATTCAG<br>AAGACTTGATATAAACCTAATTACCTCGAGAGGGGACTTTTTTACATGTGTCAGAGGTTTTCACCGTCATC<br>ACCGAAACGCGCGAGACGAAAGGGCCTCGTGATACGCCTATTTTTATAGGTTAATGTCATGATAATAATGG<br>TTTCTTAGACGTCAGGTGGCACTTTTCGGGGAAATGTGCGCGGAACCCCTATTTGTTTATTTTTCTAAATA<br>CATTCAAATATGTATCCGCTCATGAGACAATAACCCTGATAAATGCTTCAATAATATTGAAAAAGGAAGAG<br>TATGAGTATTCAACATTTCCGTGTCGCCCTTATTCCCTTTTTTGCGGCATTTTGCCTTCCTGTTTTTGCTC<br>ACCCAGAAACGCTGGTGAAAGTAAAAGATGCTGAAGATCAGTTGGGTGCACGAGTGGGTTACATCGAACTG<br>GATCTCAACAGCGGTAAGATCCTTGAGAGTTTTCGCCCCGAAGAACGTTTTCCAATGATGAGCACTTTTAA<br>AGTTCTGCTATGTGGCGCGGTATTATCCCGTATTGACGCCGGGCAAGAGCAACTCGGTCGCCGCATACACT<br>ATTCTCAGAATGACTTGGTTGAGTACTCACCAGTCACAGAAAAGCATCTTACGGATGGCATGACAGTAAGA<br>GAATTATGCAGTGCTGCCATAACCATGAGTGATAACACTGCGGCCAACTTACTTCTGACAACGATCGGAGG<br>ACCGAAGGAGCTAACCGCTTTTTTGCACAACATGGGGGATCATGTAACTCGCCTTGATCGTTGGGAACCGG<br>AGCTGAATGAAGCCATACCAAACGACGAGCGTGACACCACGATGCCTGTAGCAATGGCAACAACGTTGCGC |

Table S2

| | |
|---|---|
| | AAACTATTAACTGGCGAACTACTTACTCTAGCTTCCCGGCAACAATTAATAGACTGGATGGAGGCGGATAA<br>AGTTGCAGGACCACTTCTGCGCTCGGCCCTTCCGGCTGGCTGGTTTATTGCTGATAAATCTGGAGCCGGTG<br>AGCGTGGGTCTCGCGGTATCATTGCAGCACTGGGGCCAGATGGTAAGCCCTCCCGTATCGTAGTTATCTAC<br>ACGACGGGGAGTCAGGCAACTATGGATGAACGAAATAGACAGATCGCTGAGATAGGTGCCTCACTGATTAA<br>GCATTGGTAACTGTCAGACCAAGTTTACTCATATATACTTTAGATTGATTTAAAACTTCATTTTTAATTTA<br>AAAGGATCTAGGTGAAGATCCTTTTTGATAATCTCATGACCAAAATCCCTTAACGTGAGTTTTCGTTCCAC<br>TGAGCGTCAGACCCCGTAGAAAAGATCAAAGGATCTTCTTGAGATCCTTTTTTTCTGCGCGTAATCTGCTG<br>CTTGCAAACAAAAAAACCACCGCTACCAGCGGTGGTTTGTTTGCCGGATCAAGAGCTACCAACTCTTTTTC<br>CGAAGGTAACTGGCTTCAGCAGAGCGCAGATACCAAATACTGTTCTTCTAGTGTAGCCGTAGTTAGGCCAC<br>CACTTCAAGAACTCTGTAGCACCGCCTACATACCTCGCTCTGCTAATCCTGTTACCAGTGGCTGCTGCCAG<br>TGGCGATAAGTCGTGTCTTACCGGGTTGGACTCAAGACGATAGTTACCGGATAAGGCGCAGCGGTCGGGCT<br>GAACGGGGGGTTCGTGCACACAGCCCAGCTTGGAGCGAACGACCTACACCGAACTGAGATACCTACAGCGT<br>GAGCTATGAGAAAGCGCCACGCTTCCCGAAGGGAGAAAGGCGGACAGGTATCCGGTAAGCGGCAGGGTCGG<br>AACAGGAGAGCGCACGAGGGAGCTTCCAGGGGGAAACGCCTGGTATCTTTATAGTCCTGTCGGGTTTCGCC<br>ACCTCTGACTTGAGCGTCGATTTTTGTGATGCTCGTCAGGGGGGCGGAGCCTATGGAAAAACGCCAGCAAC<br>GCGGCCTTTTTCTTAAGC |

| **Plasmid** | **pDAC760** |
|---|---|
| **Description** | Expression of Csm complex (DNase/cA mut) from separate promoters; GFP backbone |
| **Utility** | RNA KD |
| **Features** | Pcmv-FLAG-NLS-Csm1(DNase/cA mut)-pA; Pcmv-FLAG-NLS-Csm2-pA, Pcmv-FLAG-NLS-Csm3-pA; Pcmv-FLAG-NLS-Csm4-pA; Pcmv-FLAG-NLS-Csm5-pA; Pcmv-FLAG-NLS-Cas6-pA; Pu6-crRNA-pT; Pcmv-GFP-pA |
| **Sequence** | GTGATGCGGTTTTGGCAGTACATCAATGGGCGTGGATAGCGGTTTGACTCACGGGGATTTCCAAGTCTCCA<br>CCCCATTGACGTCAATGGGAGTTTGTTTTGGCACCAAAATCAACGGGACTTTCCAAAATGTCGTAACAACT<br>CCGCCCCATTGACGCAAATGGGCGGTAGGCGTGTACGGTGGGAGGTCTATATAAGCAGAGCTCGTTTAGTG<br>AACCGTCAGATCTCTAGAgccgccaccATGCACCATCACCATCACCATTCCGGCGATTACAAAGACGATGA<br>CGATAAGATGGCGCCTAAGAAGAAACGCAAAGTGCGGGGCATGAAGAAAGAAAAGATTGATCTGTTTTACG<br>GAGCCCTGCTGGCCGCCATCGGAAAGGTCATCCAGCGAGCAACCGGAGAGCGGAAGAAACACGCACTTGTG<br>GGCGCCGACTGGTTCGACGAGATCGCCGACAACCAAGTCATCTCGGATCAGATCCGGTACCATATGGCCAA<br>CTACCAGTCTGATAAGCTCGGCAACGATCACCTGGCTTACATCACCTACATTGCCGACAACATCGCCTCCG<br>GTGTCGACCGCCGGCAATCCAACGAAGAGTCAGACGAAGATACCTCCGCAAAGATCTGGGACACCTACACG<br>AACCAGGCCGACATCTTTAACGTGTTCGGAGCGCAGACCGATAAGCGGTACTTCAAGCCTACCGTGCTGAA<br>TCTCAAGTCGAAGCCCAACTTCGCGTCCGCCACTTACGAACCCTTTAGCAAGGGCGATTACGCTGCCATCG<br>CCACCCGGATTAAGAACGAACTGGCCGAGTTCGAGTTCAACCAAGTCCAGATTGACTCCCTGCTCAACCTT<br>TTCGAGGCTACTCTCTCCTTCGTGCCGTCAAGCACCAACACTAAGGAAATCGCCGACATCTCCCTGGCCGA<br>CCATTCCCGCTTGACTGCTGCCTTCGCTCTGGCGATCTACGACTACCTGGAGGACAAGGGTCGGCACAACT<br>ACAAAGAGGACCTGTTCACCAAAGTGTCAGCGTTCTATGAAGAAGAAGCCTTCCTGCTGGCCTCCTTCGAC<br>CTGTCGGGAATCCAGGACTTTATCTACAACATTAACATCGCAACTAACGGCGCGGCGAAGCAGCTGAAGGC<br>CCGGAGCCTCTACCTGGACTTTATGTCCGAGTACATCGCCGATAGCCTGCTGGACAAGCTGGGACTGAACA<br>GGGCTAACATGCTTTACGTCGGCGGCGGACACGCCTACTTCGTCCTGGCCAACACCGAAAAGACTGTGGAA<br>ACCCTGGTGCAGTTTGAGAAGGATTTCAACCAGTTCCTGTTGGCAAACTTCCAGACCCGCCTCTATGTGGC<br>CTTTGGCTGGGGTTCCTTCGCGGCCAAGGACATCATGTCCGAGCTGAATAGCCCCGAGTCCTACCGCCAAG<br>TGTACCAAAAGGCTTCGCGCATGATCTCCAAAAAGAAAATCTCCAGATACGACTACCAGACACTGATGCTC<br>CTGAATCGCGGTGGAAAGTCCTCAGAGAGAGAGTGCGAGATTTGCCACTCCGTGGAGAACCTGGTGTCCTA<br>CCACGACCAGAAAGTCTGTGACATTTGCCGGGGACTGTACCAGTTCTCGAAAGAAATTGCCCATGACCACT<br>TCATCATTACCGAAAATGAGGGGCTGCCGATTGGACCAAACGCGTGCTTAAAGGGCGTGGCATTCGAAAAG<br>CTGTCCCAAGAAGCGTTCAGCCGGGTCTACGTGAAGAATGACTATAAGGCCGGTACCGTGAAGGCTACGCA<br>TGTGTTCGTGGGGGATTACCAGTGCGACGAGATCTACAACTACGCCGCCCTGAGCAAGAACGAGAACGGCC<br>TAGGCATCAAGAGACTGGCCGTGGTCCGGCTCGACGTGGATGACTTGGGCGCCGCCTTCATGGCCGGTTTC<br>AGCCAGCAGGGAAACGGACAATACTCCACTCTGTCAAGATCGGCCACATTCTCCCGGAGCATGTCGCTGTT<br>CTTCAAAGTGTACATTAACCAGTTCGCCTCCGACAAGAAGCTGAGCATTATCTACGCGGGCGGCGCCGCCG<br>TGTTCGCCATTGGATCGTGGCAGGATATCATCGCGTTCACTGTGGAACTTCGCGAAAACTTCATCAAGTGG<br>ACCAACGGGAAGCTCACCCTCTCCGCGGGGATAGGGTTGTTCGCCGACAAGACTCCTATTAGCCTGATGGC<br>TCACCAGACCGGGGAACTGGAAGAGGCCGCCAAGGGCAACGAAAAGGACTCCATCTCGCTGTTCTCAAGCG<br>ACTACACTTTCAAGTTTGATAGGTTCATCACTAACGTGTACGACGACAAACTGGAACAGATTAGATACTTC<br>TTCAACCATCAAGACGAGAGGGGAAAGAACTTCATCTATAAGCTTATTGAGCTTTTGAGGAACCACGACCG<br>CATGAATATGGCACGCCTCGCCTATTACCTCACTCGCCTGGAAGAACTGACCCGGGAGACTGACAGGGACA<br>AGTTCAAGACCTTCAAGAACCTGTTCTACTCCTGGTACACCAACAAGAACGATAAGGACCGGAAGGAAGCC<br>GAGCTCGCGCTCCTGCTGTACATCTACGAAATCAGAAAGGATTAAcggcaataaaaagacagaataaaacg<br>cacggtgttgggtcgtttgttcGCACACATTAGCTAGCCGTCAGCACACATTGTGATGCGGTTTTGGCAGT<br>ACATCAATGGGCGTGGATAGCGGTTTGACTCACGGGGATTTCCAAGTCTCCACCCCATTGACGTCAATGGG |

Table S2

| | |
|---|---|
| | AGTTTGTTTTGGCACCAAAATCAACGGGACTTTCCAAAATGTCGTAACAACTCCGCCCCATTGACGCAAAT<br>GGGCGGTAGGCGTGTACGGTGGGAGGTCTATATAAGCAGAGCTCGTTTAGTGAACCGTCAGATCTCTAGAg<br>ccgccaccATGGATTACAAAGACGATGACGATAAGATGGCGCCTAAGAAGAAACGCAAAGTGCGGGGCATG<br>ACCATCCTGACCGACGAGAACTACGTGGACATCGCCGAGAAAGCCATCCTGAAGCTGGAAAGAAACACCAG<br>AAATAGAAAGAACCCTGATGCCTTCTTCCTGACCACATCTAAGCTGCGGAACCTGCTGAGCCTGACAAGCA<br>CCCTGTTCGACGAGAGCAAGGTGAAGGAATACGACGCCCTGCTGGACAGAATCGCTTATCTGAGAGTGCAG<br>TTCGTGTACCAGGCCGGCAGAGAGATCGCCGTGAAAGATCTGATCGAGAAGGCCCAGATCCTGGAAGCTCT<br>GAAAGAGATCAAGGACCGGGAAACCCTGCAGAGATTCTGCAGATACATGGAAGCCCTGGTGGCCTACTTCA<br>AGTTCTACGGCGGCAAGGACTGAcggcaataaaaagacagaataaaacgcacggtgttgggtcgtttgttc<br>AGTTCTTTTGCCTTACTTTCAATGCATGCGGTGATGCGGTTTTGGCAGTACATCAATGGGCGTGGATAGCG<br>GTTTGACTCACGGGGATTTCCAAGTCTCCACCCCATTGACGTCAATGGGAGTTTGTTTTGGCACCAAAATC<br>AACGGGACTTTCCAAAATGTCGTAACAACTCCGCCCCATTGACGCAAATGGGCGGTAGGCGTGTACGGTGG<br>GAGGTCTATATAAGCAGAGCTCGTTTAGTGAACCGTCAGATCTCTAGAgccgccaccATGGATTACAAAGA<br>CGATGACGATAAGATGGCGCCTAAGAAGAAACGCAAAGTGCGGGGCATGACCTTCGCCAAGATCAAATTCA<br>GCGCCCAGATCCGGCTGGAAACCGGCCTGCACATCGGAGGATCTGATGCCTTTGCCGCTATCGGCGCCATC<br>GACAGCCCTGTGATCAAGGACCCCATCACCAACCTGCCTATCATCCCCGGCTCTAGCCTGAAGGGCAAGAT<br>GAGAACACTGCTGGCCAAGGTGTACAACGAAAAGGTGGCCGAGAAGCCTAGCGACGACAGCGACATCCTGA<br>GCAGACTGTTCGGAAATAGCAAGGATAAGCGGTTCAAGATGGGCAGACTGATCTTCCGGGACGCCTTCCTG<br>AGCAACGCCGACGAGCTGGATTCTCTGGGCGTGCGGAGCTACACCGAGGTGAAGTTCGAGAACACCATCGA<br>TAGAATCACCGCCGAGGCCAATCCTAGACAGATCGAGAGAGCCATTCGGAACTCAACATTCGACTTCGAGC<br>TGATCTACGAGATCACTGATGAGAATGAGAACCAGGTCGAGGAAGATTTCAAGGTGATCAGAGACGGCCTG<br>AAGCTGCTGGAACTGGACTACCTGGGCGGAAGCGGCTCCAGAGGCTACGGCAAAGTGGCTTTTGAGAACCT<br>GAAAGCCACCACAGTGTTCGGCAACTACGACGTGAAAACCCTGAACGAGCTGCTGACCGCCGAAGTGTGAc<br>ggcaataaaaagacagaataaaacgcacggtgttgggtcgtttgttcTGGATTGCGAGAATGGACTAGTAG<br>CAAACTGTGATGCGGTTTTGGCAGTACATCAATGGGCGTGGATAGCGGTTTGACTCACGGGGATTTCCAAG<br>TCTCCACCCCATTGACGTCAATGGGAGTTTGTTTTGGCACCAAAATCAACGGGACTTTCCAAAATGTCGTA<br>ACAACTCCGCCCCATTGACGCAAATGGGCGGTAGGCGTGTACGGTGGGAGGTCTATATAAGCAGAGCTCGT<br>TTAGTGAACCGTCAGATCTCTAGAgccgccaccATGGATTACAAAGACGATGACGATAAGATGGCGCCTAA<br>GAAGAAACGCAAAGTGCGGGGCATGACTTACAAGCTCTACATTATGACCTTTCAAAACGCCCACTTCGGTT<br>CCGGCACTCTGGACTCATCGAAGCTGACCTTCTCCGCGGATAGAATCTTCTCGGCACTCGTGCTCGAGGCT<br>CTGAAGATGGGAAAGCTCGACGCCTTCTTGGCCGAGGCCAACCAGGATAAGTTCACTCTGACCGACGCGTT<br>CCCATTCCAATTCGGTCCTTTCCTGCCGAAACCGATTGGTTACCCCAAGCACGACCAGATCGACCAGTCTG<br>TGGACGTGAAGGAAGTCCGCCGCCAAGCGAAGCTGTCCAAAAAGCTCCAGTTCCTGGCTCTGGAAAACGTC<br>GACGACTACCTGAACGGAGAGCTGTTTGAGAATGAGGAACACGCCGTGATCGACACAGTGACCAAGAACCA<br>GCCCCATAAAGATGATAATCTGTACCAAGTGGCCACCACTCGGTTCTCGAACGACACCTCCCTTTACGTGA<br>TCGCCAACGAATCCGATCTGCTGAACGAACTGATGAGCAGCCTTCAGTACTCCGGGCTGGGCGGCAAAAGG<br>TCCTCAGGATTCGGCAGATTTGAGCTGGACATCCAGAACATTCCCTTGGAACTGTCCGACCGGCTGACGAA<br>GAACCACAGCGACAAGGTCATGTCACTTACCACCGCCCTCCCGGTGGACGCTGATCTCGAGGAAGCGATGG<br>AAGATGGCCATTACCTGTTGACCAAGTCGTCCGGATTCGCATTCTCCCACGCCACCAACGAAAACTATCGG<br>AAGCAGGACCTGTACAAGTTCGCCTCCGGGAGCACCTTCAGCAAGACTTTCGAGGGACAGATCGTGGACGT<br>GCGCCCTCTCGATTTCCCTCACGCCGTGCTGAACTACGCCAAGCCGCTGTTCTTTAAGCTCGAAGTCTAAc<br>ggcaataaaaagacagaataaaacgcacggtgttgggtcgtttgttcGCACATTCAAAAACAGGCAATTGG<br>ACAAGCGTGATGCGGTTTTGGCAGTACATCAATGGGCGTGGATAGCGGTTTGACTCACGGGGATTTCCAAG<br>TCTCCACCCCATTGACGTCAATGGGAGTTTGTTTTGGCACCAAAATCAACGGGACTTTCCAAAATGTCGTA<br>ACAACTCCGCCCCATTGACGCAAATGGGCGGTAGGCGTGTACGGTGGGAGGTCTATATAAGCAGAGCTCGT<br>TTAGTGAACCGTCAGATCTCTAGAgccgccaccATGGATTACAAAGACGATGACGATAAGATGGCGCCTAA<br>GAAGAAACGCAAAGTGCGGGGCATGAAAAATGACTACCGGACCTTCAAGCTGAGCCTGCTGACCCTGGCTC<br>CTATCCACATCGGCAACGGCGAGAAGTACACCAGCAGAGAATTCATCTACGAGAACAAGAAGTTCTACTTC<br>CCCGACATGGGCAAGTTCTACAACAAGATGGTGGAAAAGAGACTGGCCGAGAAGTTCGAGGCCTTCCTGAT<br>CCAGACCAGACCCAACGCCAGAAACAACCGGCTGATTTCTTTTCTGAACGACAACAGAATCGCCGAAAGAT<br>CTTTTGGCGGCTACAGCATCAGTGAAACCGGCCTGGAATCTGATAAGAACCCTAACAGCGCCGGAGCTATC<br>AACGAGGTGAACAAATTCATCCGGGACGCCTTCGGAAATCCTTACATCCCAGGCAGCAGCCTGAAGGGCGC<br>CATCCGCACCATCCTGATGAACACCACACCTAAGTGGAACAACGAGAACGCCGTGAACGACTTCGGCAGAT<br>TCCCAAAGGAAAACAAGAACCTGATCCCTTGGGGACCTAAGAAAGGCAAGGAATACGACGACCTGTTCAAC<br>GCCATCAGAGTGTCCGACAGCAAGCCCTTCGACAACAAAAGCCTGATCCTCGTGCAGAAGTGGGACTACAG<br>CGCCAAAACCAACAAGGCCAAGCCTCTGCCTCTGTACAGAGAGTCTATCAGCCCTCTGACCAAGATCGAGT<br>TCGAGATAACAACAACCACTGATGAGGCCGGCAGACTGATCGAGGAACTGGGAAAGCGGGCCCAGGCCTTT<br>TATAAGGACTACAAGGCCTTTTTCCTGTCTGAATTCCCTGATGATAAGATCCAGGCTAATCTGCAATACCC<br>CATCTACCTGGGCGCCGGCAGCGGCGCTTGGACAAAGACCCTGTTTAAGCAGGCCGACGGCATCCTGCAGC<br>GGAGATACTCCAGAATGAAAACCAAGATGGTCAAGAAGGGCGTGCTGAAGCTGACAAAGGCCCCTCTGAAA<br>ACAGTGAAGATCCCCAGCGGCAACCACAGCCTGGTGAAGAATCACGAGAGCTTCTACGAGATGGGCAAAGC<br>CAACTTCATGATCAAGGAAATCGACAAGTGAcggcaataaaaagacagaataaaacgcacggtgttgggtc<br>gtttgttcATGTGTGCCGGCCGGAGAATAAAAGTCTAAGTGATGCGGTTTTGGCAGTACATCAATGGGCGT |

Table S2

| | |
|---|---|
| | GGATAGCGGTTTGACTCACGGGGATTTCCAAGTCTCCACCCCATTGACGTCAATGGGAGTTTGTTTTGGCA<br>CCAAAATCAACGGGACTTTCCAAAATGTCGTAACAACTCCGCCCCATTGACGCAAATGGGCGGTAGGCGTG<br>TACGGTGGGAGGTCTATATAAGCAGAGCTCGTTTAGTGAACCGTCAGATCTCTAGAgccgccaccATGGAT<br>TACAAAGACGATGACGATAAGATGGCGCCTAAGAAGAAACGCAAAGTGCGGGGCATGAAAAAGCTCGTGTT<br>CACCTTTAAGCGGATCGACCACCCTGCTCAGGACCTGGCCGTGAAATTCCACGGCTTCCTGATGGAACAGC<br>TGGATAGCGACTACGTGGACTACCTGCACCAGCAGCAGACCAACCCCTACGCCACAAAGGTGATCCAGGGC<br>AAAGAGAACACCCAGTGGGTCGTGCATCTGCTGACAGACGACATCGAGGACAAGGTGTTCATGACCCTGCT<br>GCAGATCAAGGAAGTGTCCCTGAACGACCTGCCTAAGTTGTCTGTGGAAAAGGTGGAAATCCAGGAGCTGG<br>GCGCTGATAAGCTGCTCGAGATCTTCAACAGCGAGGAAAACCAGACCTACTTCAGCATCATCTTCGAGACA<br>CCTACAGGCTTTAAAAGCCAGGGCAGCTACGTGATCTTCCCCAGCATGCGGCTGATCTTTCAGAGCCTGAT<br>GCAGAAGTACGGCAGACTGGTGGAAAACCAGCCTGAGATCGAGGAAGATACCCTGGACTACCTGAGCGAGC<br>ACAGCACCATCACCAATTACAGACTGGAAACAAGCTACTTCAGAGTGCATAGACAGAGAATCCCCGCCTTC<br>CGGGGCAAGCTGACCTTCAAGGTGCAGGGAGCCCAGACACTGAAGGCCTACGTGAAGATGCTGCTGACCTT<br>CGGCGAGTACAGCGGCCTGGGCATGAAAACCAGCCTGGGAATGGGCGGCATCAAGCTGGAAGAAAGAAAGG<br>ACTGAcggcaataaaaagacagaataaaacgcacggtgttgggtcgtttgttcGTGCGATAGAGGGATCCC<br>GCATTGAATTATGTGATGCGGTTTTGGCAGTACATCAATGGGCGTGGATAGCGGTTTGACTCACGGGGATT<br>TCCAAGTCTCCACCCCATTGACGTCAATGGGAGTTTGTTTTGGCACCAAAATCAACGGGACTTTCCAAAAT<br>GTCGTAACAACTCCGCCCCATTGACGCAAATGGGCGGTAGGCGTGTACGGTGGGAGGTCTATATAAGCAGA<br>GCTCGTTTAGTGAACCGTCAGATCTCTAGAgccgccaccatggtgagcaagggcgaggagctgttcaccgg<br>ggtggtgcccatcctggtcgagctggacggcgacgtaaacggccacaagttcagcgtgtccggcgagggcg<br>agggcgatgccacctacggcaagctgaccctgaagttcatctgcaccaccggcaagctgcccgtgccctgg<br>cccaccctcgtgaccaccctgacctacggcgtgcagtgcttcagccgctaccccgaccacatgaagcagca<br>cgacttcttcaagtccgccatgcccgaaggctacgtccaggagcgcaccatcttcttcaaggacgacggca<br>actacaagacccgcgccgaggtgaagttcgagggcgacaccctggtgaaccgcatcgagctgaagggcatc<br>gacttcaaggaggacggcaacatcctggggcacaagctggagtacaactacaacagccacaacgtctatat<br>catggccgacaagcagaagaacggcatcaaggtgaacttcaagatccgccacaacatcgaggacggcagcg<br>tgcagctcgccgaccactaccagcagaacacccccatcggcgacggccccgtgctgctgcccgacaaccac<br>tacctgagcacccagtccgccctgagcaaagaccccaacgagaagcgcgatcacatggtcctgctggagtt<br>cgtgaccgccgccgggatcactctcggcatggacgagctgtacaagtgacggcaataaaaagacagaataa<br>aacgcacggtgttgggtcgtttgttcGTAGATGGCGCGCCTTTGTTGACCCGGTTGGAGGGCCTATTTCCC<br>ATGATTCCTTCATATTTGCATATACGATACAAGGCTGTTAGAGAGATAATTAGAATTAATTTGACTGTAAA<br>CACAAAGATATTAGTACAAAATACGTGACGTAGAAAGTAATAATTTCTTGGGTAGTTTGCAGTTTTAAAAT<br>TATGTTTTAAAATGGACTATCATATGCTTACCGTAACTTGAAAGTATTTCGATTTCTTGGCTTTATATATC<br>TTGTGGAAAGGACGAAACACCGATATAAACCTAATTACCTCGAGAGGGGACGGAAACCCGTCTTCGATGAA<br>GCGATTCAGAAGACTTGATATAAACCTAATTACCTCGAGAGGGGACTTTTTTACATGTGTCAGAGGTTTTC<br>ACCGTCATCACCGAAACGCGCGAGACGAAAGGGCCTCGTGATACGCCTATTTTTATAGGTTAATGTCATGA<br>TAATAATGGTTTCTTAGACGTCAGGTGGCACTTTTCGGGGAAATGTGCGCGGAACCCCTATTTGTTTATTT<br>TTCTAAATACATTCAAATATGTATCCGCTCATGAGACAATAACCCTGATAAATGCTTCAATAATATTGAAA<br>AAGGAAGAGTATGAGTATTCAACATTTCCGTGTCGCCCTTATTCCCTTTTTTGCGGCATTTTGCCTTCCTG<br>TTTTTGCTCACCCAGAAACGCTGGTGAAAGTAAAAGATGCTGAAGATCAGTTGGGTGCACGAGTGGGTTAC<br>ATCGAACTGGATCTCAACAGCGGTAAGATCCTTGAGAGTTTTCGCCCCGAAGAACGTTTTCCAATGATGAG<br>CACTTTTAAAGTTCTGCTATGTGGCGCGGTATTATCCCGTATTGACGCCGGGCAAGAGCAACTCGGTCGCC<br>GCATACACTATTCTCAGAATGACTTGGTTGAGTACTCACCAGTCACAGAAAAGCATCTTACGGATGGCATG<br>ACAGTAAGAGAATTATGCAGTGCTGCCATAACCATGAGTGATAACACTGCGGCCAACTTACTTCTGACAAC<br>GATCGGAGGACCGAAGGAGCTAACCGCTTTTTTGCACAACATGGGGGATCATGTAACTCGCCTTGATCGTT<br>GGGAACCGGAGCTGAATGAAGCCATACCAAACGACGAGCGTGACACCACGATGCCTGTAGCAATGGCAACA<br>ACGTTGCGCAAACTATTAACTGGCGAACTACTTACTCTAGCTTCCCGGCAACAATTAATAGACTGGATGGA<br>GGCGGATAAAGTTGCAGGACCACTTCTGCGCTCGGCCCTTCCGGCTGGCTGGTTTATTGCTGATAAATCTG<br>GAGCCGGTGAGCGTGGGTCTCGCGGTATCATTGCAGCACTGGGGCCAGATGGTAAGCCCTCCCGTATCGTA<br>GTTATCTACACGACGGGGAGTCAGGCAACTATGGATGAACGAAATAGACAGATCGCTGAGATAGGTGCCTC<br>ACTGATTAAGCATTGGTAACTGTCAGACCAAGTTTACTCATATATACTTTAGATTGATTTAAAACTTCATT<br>TTTAATTTAAAAGGATCTAGGTGAAGATCCTTTTTGATAATCTCATGACCAAAATCCCTTAACGTGAGTTT<br>TCGTTCCACTGAGCGTCAGACCCCGTAGAAAAGATCAAAGGATCTTCTTGAGATCCTTTTTTTCTGCGCGT<br>AATCTGCTGCTTGCAAACAAAAAAACCACCGCTACCAGCGGTGGTTTGTTTGCCGGATCAAGAGCTACCAA<br>CTCTTTTTCCGAAGGTAACTGGCTTCAGCAGAGCGCAGATACCAAATACTGTTCTTCTAGTGTAGCCGTAG<br>TTAGGCCACCACTTCAAGAACTCTGTAGCACCGCCTACATACCTCGCTCTGCTAATCCTGTTACCAGTGGC<br>TGCTGCCAGTGGCGATAAGTCGTGTCTTACCGGGTTGGACTCAAGACGATAGTTACCGGATAAGGCGCAGC<br>GGTCGGGCTGAACGGGGGGTTCGTGCACACAGCCCAGCTTGGAGCGAACGACCTACACCGAACTGAGATAC<br>CTACAGCGTGAGCTATGAGAAAGCGCCACGCTTCCCGAAGGGAGAAAGGCGGACAGGTATCCGGTAAGCGG<br>CAGGGTCGGAACAGGAGAGCGCACGAGGGAGCTTCCAGGGGGAAACGCCTGGTATCTTTATAGTCCTGTCG<br>GGTTTCGCCACCTCTGACTTGAGCGTCGATTTTTGTGATGCTCGTCAGGGGGGCGGAGCCTATGGAAAAAC<br>GCCAGCAACGCGGCCTTTTTCTTAAGC |

Table S2

| | |
|---|---|
| **Plasmid** | **pDAC761** |
| **Description** | Expression of Csm complex (DNase/cA mut) from separate promoters; Puro backbone |
| **Utility** | RNA KD |
| **Features** | Pcmv-FLAG-NLS-Csm1(DNase/cA mut)-pA; Pcmv-FLAG-NLS-Csm2-pA, Pcmv-FLAG-NLS-Csm3-pA; Pcmv-FLAG-NLS-Csm4-pA; Pcmv-FLAG-NLS-Csm5-pA; Pcmv-FLAG-NLS-Cas6-pA; Pu6-crRNA-pT; Pcmv-Puro-pA |
| **Sequence** | GTGATGCGGTTTTGGCAGTACATCAATGGGCGTGGATAGCGGTTTGACTCACGGGGATTTCCAAGTCTCCA<br>CCCCATTGACGTCAATGGGAGTTTGTTTTGGCACCAAAATCAACGGGACTTTCCAAAATGTCGTAACAACT<br>CCGCCCCATTGACGCAAATGGGCGGTAGGCGTGTACGGTGGGAGGTCTATATAAGCAGAGCTCGTTTAGTG<br>AACCGTCAGATCTCTAGAgccgccaccATGCACCATCACCATCACCATTCCGGCGATTACAAAGACGATGA<br>CGATAAGATGGCGCCTAAGAAGAAACGCAAAGTGCGGGGCATGAAGAAAGAAAAGATTGATCTGTTTTACG<br>GAGCCCTGCTGGCCGCCATCGGAAAGGTCATCCAGCGAGCAACCGGAGAGCGGAAGAAACACGCACTTGTG<br>GGCGCCGACTGGTTCGACGAGATCGCCGACAACCAAGTCATCTCGGATCAGATCCGGTACCATATGGCCAA<br>CTACCAGTCTGATAAGCTCGGCAACGATCACCTGGCTTACATCACCTACATTGCCGACAACATCGCCTCCG<br>GTGTCGACCGCCGGCAATCCAACGAAGAGTCAGACGAAGATACCTCCGCAAAGATCTGGGACACCTACACG<br>AACCAGGCCGACATCTTTAACGTGTTCGGAGCGCAGACCGATAAGCGGTACTTCAAGCCTACCGTGCTGAA<br>TCTCAAGTCGAAGCCCAACTTCGCGTCCGCCACTTACGAACCCTTTAGCAAGGGCGATTACGCTGCCATCG<br>CCACCCGGATTAAGAACGAACTGGCCGAGTTCGAGTTCAACCAAGTCCAGATTGACTCCCTGCTCAACCTT<br>TTCGAGGCTACTCTCTCCTTCGTGCCGTCAAGCACCAACACTAAGGAAATCGCCGACATCTCCCTGGCCGA<br>CCATTCCCGCTTGACTGCTGCCTTCGCTCTGGCGATCTACGACTACCTGGAGGACAAGGGTCGGCACAACT<br>ACAAAGAGGACCTGTTCACCAAAGTGTCAGCGTTCTATGAAGAAGAAGCCTTCCTGCTGGCCTCCTTCGAC<br>CTGTCGGGAATCCAGGACTTTATCTACAACATTAACATCGCAACTAACGGCGCGGCGAAGCAGCTGAAGGC<br>CCGGAGCCTCTACCTGGACTTTATGTCCGAGTACATCGCCGATAGCCTGCTGGACAAGCTGGGACTGAACA<br>GGGCTAACATGCTTTACGTCGGCGGCGGACACGCCTACTTCGTCCTGGCCAACACCGAAAAGACTGTGGAA<br>ACCCTGGTGCAGTTTGAGAAGGATTTCAACCAGTTCCTGTTGGCAAACTTCCAGACCCGCCTCTATGTGGC<br>CTTTGGCTGGGGTTCCTTCGCGGCCAAGGACATCATGTCCGAGCTGAATAGCCCCGAGTCCTACCGCCAAG<br>TGTACCAAAAGGCTTCGCGCATGATCTCCAAAAAGAAAATCTCCAGATACGACTACCAGACACTGATGCTC<br>CTGAATCGCGGTGGAAAGTCCTCAGAGAGAGAGTGCGAGATTTGCCACTCCGTGGAGAACCTGGTGTCCTA<br>CCACGACCAGAAAGTCTGTGACATTTGCCGGGGACTGTACCAGTTCTCGAAAGAAATTGCCCATGACCACT<br>TCATCATTACCGAAAATGAGGGGCTGCCGATTGGACCAAACGCGTGCTTAAAGGGCGTGGCATTCGAAAAG<br>CTGTCCCAAGAAGCGTTCAGCCGGGTCTACGTGAAGAATGACTATAAGGCCGGTACCGTGAAGGCTACGCA<br>TGTGTTCGTGGGGGATTACCAGTGCGACGAGATCTACAACTACGCCGCCCTGAGCAAGAACGAGAACGGCC<br>TAGGCATCAAGAGACTGGCCGTGGTCCGGCTCGACGTGGATGACTTGGGCGCCGCCTTCATGGCCGGTTTC<br>AGCCAGCAGGGAAACGGACAATACTCCACTCTGTCAAGATCGGCCACATTCTCCCGGAGCATGTCGCTGTT<br>CTTCAAAGTGTACATTAACCAGTTCGCCTCCGACAAGAAGCTGAGCATTATCTACGCGGGCGGCGCCGCCG<br>TGTTCGCCATTGGATCGTGGCAGGATATCATCGCGTTCACTGTGGAACTTCGCGAAAACTTCATCAAGTGG<br>ACCAACGGGAAGCTCACCCTCTCCGCGGGGATAGGGTTGTTCGCCGACAAGACTCCTATTAGCCTGATGGC<br>TCACCAGACCGGGGAACTGGAAGAGGCCGCCAAGGGCAACGAAAAGGACTCCATCTCGCTGTTCTCAAGCG<br>ACTACACTTTCAAGTTTGATAGGTTCATCACTAACGTGTACGACGACAAACTGGAACAGATTAGATACTTC<br>TTCAACCATCAAGACGAGAGGGGAAAGAACTTCATCTATAAGCTTATTGAGCTTTTGAGGAACCACGACCG<br>CATGAATATGGCACGCCTCGCCTATTACCTCACTCGCCTGGAAGAACTGACCCGGGAGACTGACAGGGACA<br>AGTTCAAGACCTTCAAGAACCTGTTCTACTCCTGGTACACCAACAAGAACGATAAGGACCGGAAGGAAGCC<br>GAGCTCGCGCTCCTGCTGTACATCTACGAAATCAGAAAGGATTAAcggcaataaaaagacagaataaaacg<br>cacggtgttgggtcgtttgttcGCACACATTAGCTAGCCGTCAGCACACATTGTGATGCGGTTTTGGCAGT<br>ACATCAATGGGCGTGGATAGCGGTTTGACTCACGGGGATTTCCAAGTCTCCACCCCATTGACGTCAATGGG<br>AGTTTGTTTTGGCACCAAAATCAACGGGACTTTCCAAAATGTCGTAACAACTCCGCCCCATTGACGCAAAT<br>GGGCGGTAGGCGTGTACGGTGGGAGGTCTATATAAGCAGAGCTCGTTTAGTGAACCGTCAGATCTCTAGAg<br>ccgccaccATGGATTACAAAGACGATGACGATAAGATGGCGCCTAAGAAGAAACGCAAAGTGCGGGGCATG<br>ACCATCCTGACCGACGAGAACTACGTGGACATCGCCGAGAAAGCCATCCTGAAGCTGGAAAGAAACACCAG<br>AAATAGAAAGAACCCTGATGCCTTCTTCCTGACCACATCTAAGCTGCGGAACCTGCTGAGCCTGACAAGCA<br>CCCTGTTCGACGAGAGCAAGGTGAAGGAATACGACGCCCTGCTGGACAGAATCGCTTATCTGAGAGTGCAG<br>TTCGTGTACCAGGCCGGCAGAGAGATCGCCGTGAAAGATCTGATCGAGAAGGCCCAGATCCTGGAAGCTCT<br>GAAAGAGATCAAGGACCGGGAAACCCTGCAGAGATTCTGCAGATACATGGAAGCCCTGGTGGCCTACTTCA<br>AGTTCTACGGCGGCAAGGACTGAcggcaataaaaagacagaataaaacgcacggtgttgggtcgtttgttc<br>AGTTCTTTTGCCTTACTTTCAATGCATGCGGTGATGCGGTTTTGGCAGTACATCAATGGGCGTGGATAGCG<br>GTTTGACTCACGGGGATTTCCAAGTCTCCACCCCATTGACGTCAATGGGAGTTTGTTTTGGCACCAAAATC<br>AACGGGACTTTCCAAAATGTCGTAACAACTCCGCCCCATTGACGCAAATGGGCGGTAGGCGTGTACGGTGG<br>GAGGTCTATATAAGCAGAGCTCGTTTAGTGAACCGTCAGATCTCTAGAgccgccaccATGGATTACAAAGA<br>CGATGACGATAAGATGGCGCCTAAGAAGAAACGCAAAGTGCGGGGCATGACCTTCGCCAAGATCAAATTCA<br>GCGCCCAGATCCGGCTGGAAACCGGCCTGCACATCGGAGGATCTGATGCCTTTGCCGCTATCGGCGCCATC<br>GACAGCCCTGTGATCAAGGACCCCATCACCAACCTGCCTATCATCCCCGGCTCTAGCCTGAAGGGCAAGAT<br>GAGAACACTGCTGGCCAAGGTGTACAACGAAAAGGTGGCCGAGAAGCCTAGCGACGACAGCGACATCCTGA |

Table S2

| | |
|---|---|
| | GCAGACTGTTCGGAAATAGCAAGGATAAGCGGTTCAAGATGGGCAGACTGATCTTCCGGGACGCCTTCCTG<br>AGCAACGCCGACGAGCTGGATTCTCTGGGCGTGCGGAGCTACACCGAGGTGAAGTTCGAGAACACCATCGA<br>TAGAATCACCGCCGAGGCCAATCCTAGACAGATCGAGAGAGCCATTCGGAACTCAACATTCGACTTCGAGC<br>TGATCTACGAGATCACTGATGAGAATGAGAACCAGGTCGAGGAAGATTTCAAGGTGATCAGAGACGGCCTG<br>AAGCTGCTGGAACTGGACTACCTGGGCGGAAGCGGCTCCAGAGGCTACGGCAAAGTGGCTTTTGAGAACCT<br>GAAAGCCACCACAGTGTTCGGCAACTACGACGTGAAAACCCTGAACGAGCTGCTGACCGCCGAAGTGTGAc<br>ggcaataaaaagacagaataaaacgcacggtgttgggtcgtttgttcTGGATTGCGAGAATGGACTAGTAG<br>CAAACTGTGATGCGGTTTTGGCAGTACATCAATGGGCGTGGATAGCGGTTTGACTCACGGGGATTTCCAAG<br>TCTCCACCCCATTGACGTCAATGGGAGTTTGTTTTGGCACCAAAATCAACGGGACTTTCCAAAATGTCGTA<br>ACAACTCCGCCCCATTGACGCAAATGGGCGGTAGGCGTGTACGGTGGGAGGTCTATATAAGCAGAGCTCGT<br>TTAGTGAACCGTCAGATCTCTAGAgccgccaccATGGATTACAAAGACGATGACGATAAGATGGCGCCTAA<br>GAAGAAACGCAAAGTGCGGGGCATGACTTACAAGCTCTACATTATGACCTTTCAAAACGCCCACTTCGGTT<br>CCGGCACTCTGGACTCATCGAAGCTGACCTTCTCCGCGGATAGAATCTTCTCGGCACTCGTGCTCGAGGCT<br>CTGAAGATGGGAAAGCTCGACGCCTTCTTGGCCGAGGCCAACCAGGATAAGTTCACTCTGACCGACGCGTT<br>CCCATTCCAATTCGGTCCTTTCCTGCCGAAACCGATTGGTTACCCCAAGCACGACCAGATCGACCAGTCTG<br>TGGACGTGAAGGAAGTCCGCCGCCAAGCGAAGCTGTCCAAAAAGCTCCAGTTCCTGGCTCTGGAAAACGTC<br>GACGACTACCTGAACGGAGAGCTGTTTGAGAATGAGGAACACGCCGTGATCGACACAGTGACCAAGAACCA<br>GCCCCATAAAGATGATAATCTGTACCAAGTGGCCACCACTCGGTTCTCGAACGACACCTCCCTTTACGTGA<br>TCGCCAACGAATCCGATCTGCTGAACGAACTGATGAGCAGCCTTCAGTACTCCGGGCTGGGCGGCAAAAGG<br>TCCTCAGGATTCGGCAGATTTGAGCTGGACATCCAGAACATTCCCTTGGAACTGTCCGACCGGCTGACGAA<br>GAACCACAGCGACAAGGTCATGTCACTTACCACCGCCCTCCCGGTGGACGCTGATCTCGAGGAAGCGATGG<br>AAGATGGCCATTACCTGTTGACCAAGTCGTCCGGATTCGCATTCTCCCACGCCACCAACGAAAACTATCGG<br>AAGCAGGACCTGTACAAGTTCGCCTCCGGGAGCACCTTCAGCAAGACTTTCGAGGGACAGATCGTGGACGT<br>GCGCCCTCTCGATTTCCCTCACGCCGTGCTGAACTACGCCAAGCCGCTGTTCTTTAAGCTCGAAGTCTAAc<br>ggcaataaaaagacagaataaaacgcacggtgttgggtcgtttgttcGCACATTCAAAAACAGGCAATTGG<br>ACAAGCGTGATGCGGTTTTGGCAGTACATCAATGGGCGTGGATAGCGGTTTGACTCACGGGGATTTCCAAG<br>TCTCCACCCCATTGACGTCAATGGGAGTTTGTTTTGGCACCAAAATCAACGGGACTTTCCAAAATGTCGTA<br>ACAACTCCGCCCCATTGACGCAAATGGGCGGTAGGCGTGTACGGTGGGAGGTCTATATAAGCAGAGCTCGT<br>TTAGTGAACCGTCAGATCTCTAGAgccgccaccATGGATTACAAAGACGATGACGATAAGATGGCGCCTAA<br>GAAGAAACGCAAAGTGCGGGGCATGAAAAATGACTACCGGACCTTCAAGCTGAGCCTGCTGACCCTGGCTC<br>CTATCCACATCGGCAACGGCGAGAAGTACACCAGCAGAGAATTCATCTACGAGAACAAGAAGTTCTACTTC<br>CCCGACATGGGCAAGTTCTACAACAAGATGGTGGAAAAGAGACTGGCCGAGAAGTTCGAGGCCTTCCTGAT<br>CCAGACCAGACCCAACGCCAGAAACAACCGGCTGATTTCTTTTCTGAACGACAACAGAATCGCCGAAAGAT<br>CTTTTGGCGGCTACAGCATCAGTGAAACCGGCCTGGAATCTGATAAGAACCCTAACAGCGCCGGAGCTATC<br>AACGAGGTGAACAAATTCATCCGGGACGCCTTCGGAAATCCTTACATCCCAGGCAGCAGCCTGAAGGGCGC<br>CATCCGCACCATCCTGATGAACACCACACCTAAGTGGAACAACGAGAACGCCGTGAACGACTTCGGCAGAT<br>TCCCAAAGGAAAACAAGAACCTGATCCCTTGGGGACCTAAGAAAGGCAAGGAATACGACGACCTGTTCAAC<br>GCCATCAGAGTGTCCGACAGCAAGCCCTTCGACAACAAAAGCCTGATCCTCGTGCAGAAGTGGGACTACAG<br>CGCCAAAACCAACAAGGCCAAGCCTCTGCCTCTGTACAGAGAGTCTATCAGCCCTCTGACCAAGATCGAGT<br>TCGAGATAACAACAACCACTGATGAGGCCGGCAGACTGATCGAGGAACTGGGAAAGCGGGCCCAGGCCTTT<br>TATAAGGACTACAAGGCCTTTTTCCTGTCTGAATTCCCTGATGATAAGATCCAGGCTAATCTGCAATACCC<br>CATCTACCTGGGCGCCGGCAGCGGCGCTTGGACAAAGACCCTGTTTAAGCAGGCCGACGGCATCCTGCAGC<br>GGAGATACTCCAGAATGAAAACCAAGATGGTCAAGAAGGGCGTGCTGAAGCTGACAAAGGCCCCTCTGAAA<br>ACAGTGAAGATCCCCAGCGGCAACCACAGCCTGGTGAAGAATCACGAGAGCTTCTACGAGATGGGCAAAGC<br>CAACTTCATGATCAAGGAAATCGACAAGTGAcggcaataaaaagacagaataaaacgcacggtgttgggtc<br>gtttgttcATGTGTGCCGGCCGGAGAATAAAAGTCTAAGTGATGCGGTTTTGGCAGTACATCAATGGGCGT<br>GGATAGCGGTTTGACTCACGGGGATTTCCAAGTCTCCACCCCATTGACGTCAATGGGAGTTTGTTTTGGCA<br>CCAAAATCAACGGGACTTTCCAAAATGTCGTAACAACTCCGCCCCATTGACGCAAATGGGCGGTAGGCGTG<br>TACGGTGGGAGGTCTATATAAGCAGAGCTCGTTTAGTGAACCGTCAGATCTCTAGAgccgccaccATGGAT<br>TACAAAGACGATGACGATAAGATGGCGCCTAAGAAGAAACGCAAAGTGCGGGGCATGAAAAAGCTCGTGTT<br>CACCTTTAAGCGGATCGACCACCCTGCTCAGGACCTGGCCGTGAAATTCCACGGCTTCCTGATGGAACAGC<br>TGGATAGCGACTACGTGGACTACCTGCACCAGCAGCAGACCAACCCCTACGCCACAAAGGTGATCCAGGGC<br>AAAGAGAACACCCAGTGGGTCGTGCATCTGCTGACAGACGACATCGAGGACAAGGTGTTCATGACCCTGCT<br>GCAGATCAAGGAAGTGTCCCTGAACGACCTGCCTAAGTTGTCTGTGGAAAAGGTGGAAATCCAGGAGCTGG<br>GCGCTGATAAGCTGCTCGAGATCTTCAACAGCGAGGAAAACCAGACCTACTTCAGCATCATCTTCGAGACA<br>CCTACAGGCTTTAAAAGCCAGGGCAGCTACGTGATCTTCCCCAGCATGCGGCTGATCTTTCAGAGCCTGAT<br>GCAGAAGTACGGCAGACTGGTGGAAAACCAGCCTGAGATCGAGGAAGATACCCTGGACTACCTGAGCGAGC<br>ACAGCACCATCACCAATTACAGACTGGAAACAAGCTACTTCAGAGTGCATAGACAGAGAATCCCCGCCTTC<br>CGGGGCAAGCTGACCTTCAAGGTGCAGGGAGCCCAGACACTGAAGGCCTACGTGAAGATGCTGCTGACCTT<br>CGGCGAGTACAGCGGCCTGGGCATGAAAACCAGCCTGGGAATGGGCGGCATCAAGCTGGAAGAAAGAAAGG<br>ACTGAcggcaataaaaagacagaataaaacgcacggtgttgggtcgtttgttcGTGCGATAGAGGGATCCC<br>GCATTGAATTATGTGATGCGGTTTTGGCAGTACATCAATGGGCGTGGATAGCGGTTTGACTCACGGGGATT<br>TCCAAGTCTCCACCCCATTGACGTCAATGGGAGTTTGTTTTGGCACCAAAATCAACGGGACTTTCCAAAAT |

Table S2

| | |
|---|---|
| | GTCGTAACAACTCCGCCCCATTGACGCAAATGGGCGGTAGGCGTGTACGGTGGGAGGTCTATATAAGCAGA<br>GCTCGTTTAGTGAACCGTCAGATCTCTAGAgccgccaccATGACCGAGTACAAGCCCACGGTGCGCCTCGC<br>CACCCGCGACGACGTCCCCAGGGCCGTACGCACCCTCGCCGCCGCGTTCGCCGACTACCCCGCCACGCGCC<br>ACACCGTCGATCCGGACCGCCACATCGAGCGGGTCACCGAGCTGCAAGAACTCTTCCTCACGCGCGTCGGG<br>CTCGACATCGGCAAGGTGTGGGTCGCGGACGACGGCGCCGCGGTGGCGGTCTGGACCACGCCGGAGAGCGT<br>CGAAGCGGGGGCGGTGTTCGCCGAGATCGGCCCGCGCATGGCCGAGTTGAGCGGTTCCCGGCTGGCCGCGC<br>AGCAACAGATGGAAGGCCTCCTGGCGCCGCACCGGCCCAAGGAGCCCGCGTGGTTCCTGGCCACCGTCGGA<br>GTCTCGCCCGACCACCAGGGCAAGGGTCTGGGCAGCGCCGTCGTGCTCCCCGGAGTGGAGGCGGCGGAGCG<br>CGCCGGGGTGCCCGCCTTCCTGGAAACCTCCGCGCCCCGCAACCTCCCCTTCTACGAGCGGCTCGGCTTCA<br>CCGTCACCGCCGACGTCGAGGTGCCCGAAGGACCGCGCACCTGGTGCATGACCCGCAAGCCCGGTGCCTGA<br>cggcaataaaaagacagaataaaacgcacggtgttgggtcgtttgttcGAGCAGATTGTACTGAGAGTGCA<br>CCGGTTGGAGGGCCTATTTCCCATGATTCCTTCATATTTGCATATACGATACAAGGCTGTTAGAGAGATAA<br>TTAGAATTAATTTGACTGTAAACACAAAGATATTAGTACAAAATACGTGACGTAGAAAGTAATAATTTCTT<br>GGGTAGTTTGCAGTTTTAAAATTATGTTTTAAAATGGACTATCATATGCTTACCGTAACTTGAAAGTATTT<br>CGATTTCTTGGCTTTATATATCTTGTGGAAAGGACGAAACACCGATATAAACCTAATTACCTCGAGAGGGG<br>ACGGAAACCCGTCTTCGATGAAGCGATTCAGAAGACTTGATATAAACCTAATTACCTCGAGAGGGGACTTT<br>TTTACATGTGTCAGAGGTTTTCACCGTCATCACCGAAACGCGCGAGACGAAAGGGCCTCGTGATACGCCTA<br>TTTTTATAGGTTAATGTCATGATAATAATGGTTTCTTAGACGTCAGGTGGCACTTTTCGGGGAAATGTGCG<br>CGGAACCCCTATTTGTTTATTTTTCTAAATACATTCAAATATGTATCCGCTCATGAGACAATAACCCTGAT<br>AAATGCTTCAATAATATTGAAAAAGGAAGAGTATGAGTATTCAACATTTCCGTGTCGCCCTTATTCCCTTT<br>TTTGCGGCATTTTGCCTTCCTGTTTTTGCTCACCCAGAAACGCTGGTGAAAGTAAAAGATGCTGAAGATCA<br>GTTGGGTGCACGAGTGGGTTACATCGAACTGGATCTCAACAGCGGTAAGATCCTTGAGAGTTTTCGCCCCG<br>AAGAACGTTTTCCAATGATGAGCACTTTTAAAGTTCTGCTATGTGGCGCGGTATTATCCCGTATTGACGCC<br>GGGCAAGAGCAACTCGGTCGCCGCATACACTATTCTCAGAATGACTTGGTTGAGTACTCACCAGTCACAGA<br>AAAGCATCTTACGGATGGCATGACAGTAAGAGAATTATGCAGTGCTGCCATAACCATGAGTGATAACACTG<br>CGGCCAACTTACTTCTGACAACGATCGGAGGACCGAAGGAGCTAACCGCTTTTTTGCACAACATGGGGGAT<br>CATGTAACTCGCCTTGATCGTTGGGAACCGGAGCTGAATGAAGCCATACCAAACGACGAGCGTGACACCAC<br>GATGCCTGTAGCAATGGCAACAACGTTGCGCAAACTATTAACTGGCGAACTACTTACTCTAGCTTCCCGGC<br>AACAATTAATAGACTGGATGGAGGCGGATAAAGTTGCAGGACCACTTCTGCGCTCGGCCCTTCCGGCTGGC<br>TGGTTTATTGCTGATAAATCTGGAGCCGGTGAGCGTGGGTCTCGCGGTATCATTGCAGCACTGGGGCCAGA<br>TGGTAAGCCCTCCCGTATCGTAGTTATCTACACGACGGGGAGTCAGGCAACTATGGATGAACGAAATAGAC<br>AGATCGCTGAGATAGGTGCCTCACTGATTAAGCATTGGTAACTGTCAGACCAAGTTTACTCATATATACTT<br>TAGATTGATTTAAAACTTCATTTTTAATTTAAAAGGATCTAGGTGAAGATCCTTTTTGATAATCTCATGAC<br>CAAAATCCCTTAACGTGAGTTTTCGTTCCACTGAGCGTCAGACCCCGTAGAAAAGATCAAAGGATCTTCTT<br>GAGATCCTTTTTTTCTGCGCGTAATCTGCTGCTTGCAAACAAAAAAACCACCGCTACCAGCGGTGGTTTGT<br>TTGCCGGATCAAGAGCTACCAACTCTTTTTCCGAAGGTAACTGGCTTCAGCAGAGCGCAGATACCAAATAC<br>TGTTCTTCTAGTGTAGCCGTAGTTAGGCCACCACTTCAAGAACTCTGTAGCACCGCCTACATACCTCGCTC<br>TGCTAATCCTGTTACCAGTGGCTGCTGCCAGTGGCGATAAGTCGTGTCTTACCGGGTTGGACTCAAGACGA<br>TAGTTACCGGATAAGGCGCAGCGGTCGGGCTGAACGGGGGGTTCGTGCACACAGCCCAGCTTGGAGCGAAC<br>GACCTACACCGAACTGAGATACCTACAGCGTGAGCTATGAGAAAGCGCCACGCTTCCCGAAGGGAGAAAGG<br>CGGACAGGTATCCGGTAAGCGGCAGGGTCGGAACAGGAGAGCGCACGAGGGAGCTTCCAGGGGGAAACGCC<br>TGGTATCTTTATAGTCCTGTCGGGTTTCGCCACCTCTGACTTGAGCGTCGATTTTTGTGATGCTCGTCAGG<br>GGGGCGGAGCCTATGGAAAAACGCCAGCAACGCGGCCTTTTTCTTAAGC |

| | |
|---|---|
| **Plasmid** | **pDAC762** |
| **Description** | Expression of Csm complex (RNase/DNase/cA mut) from separate promoters; GFP backbone |
| **Utility** | RNA binding |
| **Features** | Pcmv-FLAG-NLS-Csm1(DNase/cA mut)-pA; Pcmv-FLAG-NLS-Csm2-pA, Pcmv-FLAG-NLS-Csm3(RNase mut)-pA; Pcmv-FLAG-NLS-Csm4-pA; Pcmv-FLAG-NLS-Csm5-pA; Pcmv-FLAG-NLS-Cas6-pA; Pu6-crRNA-pT; Pcmv-GFP-pA |
| **Sequence** | GTGATGCGGTTTTGGCAGTACATCAATGGGCGTGGATAGCGGTTTGACTCACGGGGATTTCCAAGTCTCCA<br>CCCCATTGACGTCAATGGGAGTTTGTTTTGGCACCAAAATCAACGGGACTTTCCAAAATGTCGTAACAACT<br>CCGCCCCATTGACGCAAATGGGCGGTAGGCGTGTACGGTGGGAGGTCTATATAAGCAGAGCTCGTTTAGTG<br>AACCGTCAGATCTCTAGAgccgccaccATGCACCATCACCATCACCATTCCGGCGATTACAAAGACGATGA<br>CGATAAGATGGCGCCTAAGAAGAAACGCAAAGTGCGGGGCATGAAGAAAGAAAAGATTGATCTGTTTTACG<br>GAGCCCTGCTGGCCGCCATCGGAAAGGTCATCCAGCGAGCAACCGGAGAGCGGAAGAAACACGCACTTGTG<br>GGCGCCGACTGGTTCGACGAGATCGCCGACAACCAAGTCATCTCGGATCAGATCCGGTACCATATGGCCAA<br>CTACCAGTCTGATAAGCTCGGCAACGATCACCTGGCTTACATCACCTACATTGCCGACAACATCGCCTCCG<br>GTGTCGACCGCCGGCAATCCAACGAAGAGTCAGACGAAGATACCTCCGCAAAGATCTGGGACACCTACACG<br>AACCAGGCCGACATCTTTAACGTGTTCGGAGCGCAGACCGATAAGCGGTACTTCAAGCCTACCGTGCTGAA<br>TCTCAAGTCGAAGCCCAACTTCGCGTCCGCCACTTACGAACCCTTTAGCAAGGGCGATTACGCTGCCATCG<br>CCACCCGGATTAAGAACGAACTGGCCGAGTTCGAGTTCAACCAAGTCCAGATTGACTCCCTGCTCAACCTT |

Table S2

| | |
|---|---|
| | TTCGAGGCTACTCTCTCCTTCGTGCCGTCAAGCACCAACACTAAGGAAATCGCCGACATCTCCCTGGCCGA<br>CCATTCCCGCTTGACTGCTGCCTTCGCTCTGGCGATCTACGACTACCTGGAGGACAAGGGTCGGCACAACT<br>ACAAAGAGGACCTGTTCACCAAAGTGTCAGCGTTCTATGAAGAAGAAGCCTTCCTGCTGGCCTCCTTCGAC<br>CTGTCGGGAATCCAGGACTTTATCTACAACATTAACATCGCAACTAACGGCGCGGCGAAGCAGCTGAAGGC<br>CCGGAGCCTCTACCTGGACTTTATGTCCGAGTACATCGCCGATAGCCTGCTGGACAAGCTGGGACTGAACA<br>GGGCTAACATGCTTTACGTCGGCGGCGGACACGCCTACTTCGTCCTGGCCAACACCGAAAAGACTGTGGAA<br>ACCCTGGTGCAGTTTGAGAAGGATTTCAACCAGTTCCTGTTGGCAAACTTCCAGACCCGCCTCTATGTGGC<br>CTTTGGCTGGGGTTCCTTCGCGGCCAAGGACATCATGTCCGAGCTGAATAGCCCCGAGTCCTACCGCCAAG<br>TGTACCAAAAGGCTTCGCGCATGATCTCCAAAAAGAAAATCTCCAGATACGACTACCAGACACTGATGCTC<br>CTGAATCGCGGTGGAAAGTCCTCAGAGAGAGAGTGCGAGATTTGCCACTCCGTGGAGAACCTGGTGTCCTA<br>CCACGACCAGAAAGTCTGTGACATTTGCCGGGGACTGTACCAGTTCTCGAAAGAAATTGCCCATGACCACT<br>TCATCATTACCGAAAATGAGGGGCTGCCGATTGGACCAAACGCGTGCTTAAAGGGCGTGGCATTCGAAAAG<br>CTGTCCCAAGAAGCGTTCAGCCGGGTCTACGTGAAGAATGACTATAAGGCCGGTACCGTGAAGGCTACGCA<br>TGTGTTCGTGGGGGATTACCAGTGCGACGAGATCTACAACTACGCCGCCCTGAGCAAGAACGAGAACGGCC<br>TAGGCATCAAGAGACTGGCCGTGGTCCGGCTCGACGTGGATGACTTGGGCGCCGCCTTCATGGCCGGTTTC<br>AGCCAGCAGGGAAACGGACAATACTCCACTCTGTCAAGATCGGCCACATTCTCCCGGAGCATGTCGCTGTT<br>CTTCAAAGTGTACATTAACCAGTTCGCCTCCGACAAGAAGCTGAGCATTATCTACGCGGGCGGCGCCGCCG<br>TGTTCGCCATTGGATCGTGGCAGGATATCATCGCGTTCACTGTGGAACTTCGCGAAAACTTCATCAAGTGG<br>ACCAACGGGAAGCTCACCCTCTCCGCGGGGATAGGGTTGTTCGCCGACAAGACTCCTATTAGCCTGATGGC<br>TCACCAGACCGGGGAACTGGAAGAGGCCGCCAAGGGCAACGAAAAGGACTCCATCTCGCTGTTCTCAAGCG<br>ACTACACTTTCAAGTTTGATAGGTTCATCACTAACGTGTACGACGACAAACTGGAACAGATTAGATACTTC<br>TTCAACCATCAAGACGAGAGGGGAAAGAACTTCATCTATAAGCTTATTGAGCTTTTGAGGAACCACGACCG<br>CATGAATATGGCACGCCTCGCCTATTACCTCACTCGCCTGGAAGAACTGACCCGGGAGACTGACAGGGACA<br>AGTTCAAGACCTTCAAGAACCTGTTCTACTCCTGGTACACCAACAAGAACGATAAGGACCGGAAGGAAGCC<br>GAGCTCGCGCTCCTGCTGTACATCTACGAAATCAGAAAGGATTAAcggcaataaaaagacagaataaaacg<br>cacggtgttgggtcgtttgttcGCACACATTAGCTAGCCGTCAGCACACATTGTGATGCGGTTTTGGCAGT<br>ACATCAATGGGCGTGGATAGCGGTTTGACTCACGGGGATTTCCAAGTCTCCACCCCATTGACGTCAATGGG<br>AGTTTGTTTTGGCACCAAAATCAACGGGACTTTCCAAAATGTCGTAACAACTCCGCCCCATTGACGCAAAT<br>GGGCGGTAGGCGTGTACGGTGGGAGGTCTATATAAGCAGAGCTCGTTTAGTGAACCGTCAGATCTCTAGAg<br>ccgccaccATGGATTACAAAGACGATGACGATAAGATGGCGCCTAAGAAGAAACGCAAAGTGCGGGGCATG<br>ACCATCCTGACCGACGAGAACTACGTGGACATCGCCGAGAAAGCCATCCTGAAGCTGGAAAGAAACACCAG<br>AAATAGAAAGAACCCTGATGCCTTCTTCCTGACCACATCTAAGCTGCGGAACCTGCTGAGCCTGACAAGCA<br>CCCTGTTCGACGAGAGCAAGGTGAAGGAATACGACGCCCTGCTGGACAGAATCGCTTATCTGAGAGTGCAG<br>TTCGTGTACCAGGCCGGCAGAGAGATCGCCGTGAAAGATCTGATCGAGAAGGCCCAGATCCTGGAAGCTCT<br>GAAAGAGATCAAGGACCGGGAAACCCTGCAGAGATTCTGCAGATACATGGAAGCCCTGGTGGCCTACTTCA<br>AGTTCTACGGCGGCAAGGACTGAcggcaataaaaagacagaataaaacgcacggtgttgggtcgtttgttc<br>AGTTCTTTTGCCTTACTTTCAATGCATGCGGTGATGCGGTTTTGGCAGTACATCAATGGGCGTGGATAGCG<br>GTTTGACTCACGGGGATTTCCAAGTCTCCACCCCATTGACGTCAATGGGAGTTTGTTTTGGCACCAAAATC<br>AACGGGACTTTCCAAAATGTCGTAACAACTCCGCCCCATTGACGCAAATGGGCGGTAGGCGTGTACGGTGG<br>GAGGTCTATATAAGCAGAGCTCGTTTAGTGAACCGTCAGATCTCTAGAgccgccaccATGGATTACAAAGA<br>CGATGACGATAAGATGGCGCCTAAGAAGAAACGCAAAGTGCGGGGCATGACCTTCGCCAAGATCAAATTCA<br>GCGCCCAGATCCGGCTGGAAACCGGCCTGCACATCGGAGGATCTGATGCCTTTGCCGCTATCGGCGCCATC<br>GCCAGCCCTGTGATCAAGGACCCCATCACCAACCTGCCTATCATCCCCGGCTCTAGCCTGAAGGGCAAGAT<br>GAGAACACTGCTGGCCAAGGTGTACAACGAAAAGGTGGCCGAGAAGCCTAGCGACGACAGCGACATCCTGA<br>GCAGACTGTTCGGAAATAGCAAGGATAAGCGGTTCAAGATGGGCAGACTGATCTTCCGGGACGCCTTCCTG<br>AGCAACGCCGACGAGCTGGATTCTCTGGGCGTGCGGAGCTACACCGAGGTGAAGTTCGAGAACACCATCGA<br>TAGAATCACCGCCGAGGCCAATCCTAGACAGATCGAGAGAGCCATTCGGAACTCAACATTCGACTTCGAGC<br>TGATCTACGAGATCACTGATGAGAATGAGAACCAGGTCGAGGAAGATTTCAAGGTGATCAGAGACGGCCTG<br>AAGCTGCTGGAACTGGACTACCTGGGCGGAAGCGGCTCCAGAGGCTACGGCAAAGTGGCTTTTGAGAACCT<br>GAAAGCCACCACAGTGTTCGGCAACTACGACGTGAAAACCCTGAACGAGCTGCTGACCGCCGAAGTGTGAc<br>ggcaataaaaagacagaataaaacgcacggtgttgggtcgtttgttcTGGATTGCGAGAATGGACTAGTAG<br>CAAACTGTGATGCGGTTTTGGCAGTACATCAATGGGCGTGGATAGCGGTTTGACTCACGGGGATTTCCAAG<br>TCTCCACCCCATTGACGTCAATGGGAGTTTGTTTTGGCACCAAAATCAACGGGACTTTCCAAAATGTCGTA<br>ACAACTCCGCCCCATTGACGCAAATGGGCGGTAGGCGTGTACGGTGGGAGGTCTATATAAGCAGAGCTCGT<br>TTAGTGAACCGTCAGATCTCTAGAgccgccaccATGGATTACAAAGACGATGACGATAAGATGGCGCCTAA<br>GAAGAAACGCAAAGTGCGGGGCATGACTTACAAGCTCTACATTATGACCTTTCAAAACGCCCACTTCGGTT<br>CCGGCACTCTGGACTCATCGAAGCTGACCTTCTCCGCGGATAGAATCTTCTCGGCACTCGTGCTCGAGGCT<br>CTGAAGATGGGAAAGCTCGACGCCTTCTTGGCCGAGGCCAACCAGGATAAGTTCACTCTGACCGACGCGTT<br>CCCATTCCAATTCGGTCCTTTCCTGCCGAAACCGATTGGTTACCCCAAGCACGACCAGATCGACCAGTCTG<br>TGGACGTGAAGGAAGTCCGCCGCCAAGCGAAGCTGTCCAAAAAGCTCCAGTTCCTGGCTCTGGAAAACGTC<br>GACGACTACCTGAACGGAGAGCTGTTTGAGAATGAGGAACACGCCGTGATCGACACAGTGACCAAGAACCA<br>GCCCCATAAAGATGATAATCTGTACCAAGTGGCCACCACTCGGTTCTCGAACGACACCTCCCTTTACGTGA<br>TCGCCAACGAATCCGATCTGCTGAACGAACTGATGAGCAGCCTTCAGTACTCCGGGCTGGGCGGCAAAAGG |

Table S2

| | |
|---|---|
| | TCCTCAGGATTCGGCAGATTTGAGCTGGACATCCAGAACATTCCCTTGGAACTGTCCGACCGGCTGACGAA<br>GAACCACAGCGACAAGGTCATGTCACTTACCACCGCCCTCCCGGTGGACGCTGATCTCGAGGAAGCGATGG<br>AAGATGGCCATTACCTGTTGACCAAGTCGTCCGGATTCGCATTCTCCCACGCCACCAACGAAAACTATCGG<br>AAGCAGGACCTGTACAAGTTCGCCTCCGGGAGCACCTTCAGCAAGACTTTCGAGGGACAGATCGTGGACGT<br>GCGCCCTCTCGATTTCCCTCACGCCGTGCTGAACTACGCCAAGCCGCTGTTCTTTAAGCTCGAAGTCTAAc<br>ggcaataaaaagacagaataaaacgcacggtgttgggtcgtttgttcGCACATTCAAAAACAGGCAATTGG<br>ACAAGCGTGATGCGGTTTTGGCAGTACATCAATGGGCGTGGATAGCGGTTTGACTCACGGGGATTTCCAAG<br>TCTCCACCCCATTGACGTCAATGGGAGTTTGTTTTGGCACCAAAATCAACGGGACTTTCCAAAATGTCGTA<br>ACAACTCCGCCCCATTGACGCAAATGGGCGGTAGGCGTGTACGGTGGGAGGTCTATATAAGCAGAGCTCGT<br>TTAGTGAACCGTCAGATCTCTAGAgccgccaccATGGATTACAAAGACGATGACGATAAGATGGCGCCTAA<br>GAAGAAACGCAAAGTGCGGGGCATGAAAAATGACTACCGGACCTTCAAGCTGAGCCTGCTGACCCTGGCTC<br>CTATCCACATCGGCAACGGCGAGAAGTACACCAGCAGAGAATTCATCTACGAGAACAAGAAGTTCTACTTC<br>CCCGACATGGGCAAGTTCTACAACAAGATGGTGGAAAAGAGACTGGCCGAGAAGTTCGAGGCCTTCCTGAT<br>CCAGACCAGACCCAACGCCAGAAACAACCGGCTGATTTCTTTTCTGAACGACAACAGAATCGCCGAAAGAT<br>CTTTTGGCGGCTACAGCATCAGTGAAACCGGCCTGGAATCTGATAAGAACCCTAACAGCGCCGGAGCTATC<br>AACGAGGTGAACAAATTCATCCGGGACGCCTTCGGAAATCCTTACATCCCAGGCAGCAGCCTGAAGGGCGC<br>CATCCGCACCATCCTGATGAACACCACACCTAAGTGGAACAACGAGAACGCCGTGAACGACTTCGGCAGAT<br>TCCCAAAGGAAAACAAGAACCTGATCCCTTGGGGACCTAAGAAAGGCAAGGAATACGACGACCTGTTCAAC<br>GCCATCAGAGTGTCCGACAGCAAGCCCTTCGACAACAAAAGCCTGATCCTCGTGCAGAAGTGGGACTACAG<br>CGCCAAAACCAACAAGGCCAAGCCTCTGCCTCTGTACAGAGAGTCTATCAGCCCTCTGACCAAGATCGAGT<br>TCGAGATAACAACAACCACTGATGAGGCCGGCAGACTGATCGAGGAACTGGGAAAGCGGGCCCAGGCCTTT<br>TATAAGGACTACAAGGCCTTTTTCCTGTCTGAATTCCCTGATGATAAGATCCAGGCTAATCTGCAATACCC<br>CATCTACCTGGGCGCCGGCAGCGGCGCTTGGACAAAGACCCTGTTTAAGCAGGCCGACGGCATCCTGCAGC<br>GGAGATACTCCAGAATGAAAACCAAGATGGTCAAGAAGGGCGTGCTGAAGCTGACAAAGGCCCCTCTGAAA<br>ACAGTGAAGATCCCCAGCGGCAACCACAGCCTGGTGAAGAATCACGAGAGCTTCTACGAGATGGGCAAAGC<br>CAACTTCATGATCAAGGAAATCGACAAGTGAcggcaataaaaagacagaataaaacgcacggtgttgggtc<br>gtttgttcATGTGTGCCGGCCGGAGAATAAAAGTCTAAGTGATGCGGTTTTGGCAGTACATCAATGGGCGT<br>GGATAGCGGTTTGACTCACGGGGATTTCCAAGTCTCCACCCCATTGACGTCAATGGGAGTTTGTTTTGGCA<br>CCAAAATCAACGGGACTTTCCAAAATGTCGTAACAACTCCGCCCCATTGACGCAAATGGGCGGTAGGCGTG<br>TACGGTGGGAGGTCTATATAAGCAGAGCTCGTTTAGTGAACCGTCAGATCTCTAGAgccgccaccATGGAT<br>TACAAAGACGATGACGATAAGATGGCGCCTAAGAAGAAACGCAAAGTGCGGGGCATGAAAAAGCTCGTGTT<br>CACCTTTAAGCGGATCGACCACCCTGCTCAGGACCTGGCCGTGAAATTCCACGGCTTCCTGATGGAACAGC<br>TGGATAGCGACTACGTGGACTACCTGCACCAGCAGCAGACCAACCCCTACGCCACAAAGGTGATCCAGGGC<br>AAAGAGAACACCCAGTGGGTCGTGCATCTGCTGACAGACGACATCGAGGACAAGGTGTTCATGACCCTGCT<br>GCAGATCAAGGAAGTGTCCCTGAACGACCTGCCTAAGTTGTCTGTGGAAAAGGTGGAAATCCAGGAGCTGG<br>GCGCTGATAAGCTGCTCGAGATCTTCAACAGCGAGGAAAACCAGACCTACTTCAGCATCATCTTCGAGACA<br>CCTACAGGCTTTAAAAGCCAGGGCAGCTACGTGATCTTCCCCAGCATGCGGCTGATCTTTCAGAGCCTGAT<br>GCAGAAGTACGGCAGACTGGTGGAAAACCAGCCTGAGATCGAGGAAGATACCCTGGACTACCTGAGCGAGC<br>ACAGCACCATCACCAATTACAGACTGGAAACAAGCTACTTCAGAGTGCATAGACAGAGAATCCCCGCCTTC<br>CGGGGCAAGCTGACCTTCAAGGTGCAGGGAGCCCAGACACTGAAGGCCTACGTGAAGATGCTGCTGACCTT<br>CGGCGAGTACAGCGGCCTGGGCATGAAAACCAGCCTGGGAATGGGCGGCATCAAGCTGGAAGAAAGAAAGG<br>ACTGAcggcaataaaaagacagaataaaacgcacggtgttgggtcgtttgttcGTGCGATAGAGGGATCCC<br>GCATTGAATTATGTGATGCGGTTTTGGCAGTACATCAATGGGCGTGGATAGCGGTTTGACTCACGGGGATT<br>TCCAAGTCTCCACCCCATTGACGTCAATGGGAGTTTGTTTTGGCACCAAAATCAACGGGACTTTCCAAAAT<br>GTCGTAACAACTCCGCCCCATTGACGCAAATGGGCGGTAGGCGTGTACGGTGGGAGGTCTATATAAGCAGA<br>GCTCGTTTAGTGAACCGTCAGATCTCTAGAgccgccaccatggtgagcaagggcgaggagctgttcaccgg<br>ggtggtgcccatcctggtcgagctggacggcgacgtaaacggccacaagttcagcgtgtccggcgagggcg<br>agggcgatgccacctacggcaagctgaccctgaagttcatctgcaccaccggcaagctgcccgtgccctgg<br>cccaccctcgtgaccaccctgacctacggcgtgcagtgcttcagccgctaccccgaccacatgaagcagca<br>cgacttcttcaagtccgccatgcccgaaggctacgtccaggagcgcaccatcttcttcaaggacgacggca<br>actacaagacccgcgccgaggtgaagttcgagggcgacaccctggtgaaccgcatcgagctgaagggcatc<br>gacttcaaggaggacggcaacatcctggggcacaagctggagtacaactacaacagccacaacgtctatat<br>catggccgacaagcagaagaacggcatcaaggtgaacttcaagatccgccacaacatcgaggacggcagcg<br>tgcagctcgccgaccactaccagcagaacacccccatcggcgacggccccgtgctgctgcccgacaaccac<br>tacctgagcacccagtccgccctgagcaaagaccccaacgagaagcgcgatcacatggtcctgctggagtt<br>cgtgaccgccgccgggatcactctcggcatggacgagctgtacaagtgacggcaataaaaagacagaataa<br>aacgcacggtgttgggtcgtttgttcGTAGATGGCGCGCCTTTGTTGACCCGGTTGGAGGGCCTATTTCCC<br>ATGATTCCTTCATATTTGCATATACGATACAAGGCTGTTAGAGAGATAATTAGAATTAATTTGACTGTAAA<br>CACAAAGATATTAGTACAAAATACGTGACGTAGAAAGTAATAATTTCTTGGGTAGTTTGCAGTTTTAAAAT<br>TATGTTTTAAAATGGACTATCATATGCTTACCGTAACTTGAAAGTATTTCGATTTCTTGGCTTTATATATC<br>TTGTGGAAAGGACGAAACACCGATATAAACCTAATTACCTCGAGAGGGGACGGAAACCCGTCTTCGATGAA<br>GCGATTCAGAAGACTTGATATAAACCTAATTACCTCGAGAGGGGACTTTTTACATGTGTCAGAGGTTTTC<br>ACCGTCATCACCGAAACGCGCGAGACGAAAGGGCCTCGTGATACGCCTATTTTTATAGGTTAATGTCATGA |

Table S2

| | |
|---|---|
| | TAATAATGGTTTCTTAGACGTCAGGTGGCACTTTTCGGGGAAATGTGCGCGGAACCCCTATTTGTTTATTT<br>TTCTAAATACATTCAAATATGTATCCGCTCATGAGACAATAACCCTGATAAATGCTTCAATAATATTGAAA<br>AAGGAAGAGTATGAGTATTCAACATTTCCGTGTCGCCCTTATTCCCTTTTTTGCGGCATTTTGCCTTCCTG<br>TTTTTGCTCACCCAGAAACGCTGGTGAAAGTAAAAGATGCTGAAGATCAGTTGGGTGCACGAGTGGGTTAC<br>ATCGAACTGGATCTCAACAGCGGTAAGATCCTTGAGAGTTTTCGCCCCGAAGAACGTTTTCCAATGATGAG<br>CACTTTTAAAGTTCTGCTATGTGGCGCGGTATTATCCCGTATTGACGCCGGGCAAGAGCAACTCGGTCGCC<br>GCATACACTATTCTCAGAATGACTTGGTTGAGTACTCACCAGTCACAGAAAAGCATCTTACGGATGGCATG<br>ACAGTAAGAGAATTATGCAGTGCTGCCATAACCATGAGTGATAACACTGCGGCCAACTTACTTCTGACAAC<br>GATCGGAGGACCGAAGGAGCTAACCGCTTTTTTGCACAACATGGGGGATCATGTAACTCGCCTTGATCGTT<br>GGGAACCGGAGCTGAATGAAGCCATACCAAACGACGAGCGTGACACCACGATGCCTGTAGCAATGGCAACA<br>ACGTTGCGCAAACTATTAACTGGCGAACTACTTACTCTAGCTTCCCGGCAACAATTAATAGACTGGATGGA<br>GGCGGATAAAGTTGCAGGACCACTTCTGCGCTCGGCCCTTCCGGCTGGCTGGTTTATTGCTGATAAATCTG<br>GAGCCGGTGAGCGTGGGTCTCGCGGTATCATTGCAGCACTGGGGCCAGATGGTAAGCCCTCCCGTATCGTA<br>GTTATCTACACGACGGGGAGTCAGGCAACTATGGATGAACGAAATAGACAGATCGCTGAGATAGGTGCCTC<br>ACTGATTAAGCATTGGTAACTGTCAGACCAAGTTTACTCATATATACTTTAGATTGATTTAAAACTTCATT<br>TTTAATTTAAAAGGATCTAGGTGAAGATCCTTTTTGATAATCTCATGACCAAAATCCCTTAACGTGAGTTT<br>TCGTTCCACTGAGCGTCAGACCCCGTAGAAAAGATCAAAGGATCTTCTTGAGATCCTTTTTTTCTGCGCGT<br>AATCTGCTGCTTGCAAACAAAAAAACCACCGCTACCAGCGGTGGTTTGTTTGCCGGATCAAGAGCTACCAA<br>CTCTTTTTCCGAAGGTAACTGGCTTCAGCAGAGCGCAGATACCAAATACTGTTCTTCTAGTGTAGCCGTAG<br>TTAGGCCACCACTTCAAGAACTCTGTAGCACCGCCTACATACCTCGCTCTGCTAATCCTGTTACCAGTGGC<br>TGCTGCCAGTGGCGATAAGTCGTGTCTTACCGGGTTGGACTCAAGACGATAGTTACCGGATAAGGCGCAGC<br>GGTCGGGCTGAACGGGGGGTTCGTGCACACAGCCCAGCTTGGAGCGAACGACCTACACCGAACTGAGATAC<br>CTACAGCGTGAGCTATGAGAAAGCGCCACGCTTCCCGAAGGGAGAAAGGCGGACAGGTATCCGGTAAGCGG<br>CAGGGTCGGAACAGGAGAGCGCACGAGGGAGCTTCCAGGGGGAAACGCCTGGTATCTTTATAGTCCTGTCG<br>GGTTTCGCCACCTCTGACTTGAGCGTCGATTTTTGTGATGCTCGTCAGGGGGGCGGAGCCTATGGAAAAAC<br>GCCAGCAACGCGGCCTTTTTCTTAAGC |

| **Plasmid** | **pDAC763** |
|---|---|
| **Description** | Expression of Csm-GFP complex (RNase/DNase/cA mut) from separate promoters |
| **Utility** | RNA imaging |
| **Features** | Pcmv-FLAG-NLS-Csm1(DNase/cA mut)-pA; Pcmv-FLAG-NLS-Csm2-pA, Pcmv-FLAG-NLS-Csm3(RNase mut)-GFP-pA; Pcmv-FLAG-NLS-Csm4-pA; Pcmv-FLAG-NLS-Csm5-pA; Pcmv-FLAG-NLS-Cas6-pA; Pu6-crRNA-pT |
| **Sequence** | GTGATGCGGTTTTGGCAGTACATCAATGGGCGTGGATAGCGGTTTGACTCACGGGGATTTCCAAGTCTCCA<br>CCCCATTGACGTCAATGGGAGTTTGTTTTGGCACCAAAATCAACGGGACTTTCCAAAATGTCGTAACAACT<br>CCGCCCCATTGACGCAAATGGGCGGTAGGCGTGTACGGTGGGAGGTCTATATAAGCAGAGCTCGTTTAGTG<br>AACCGTCAGATCTCTAGAgccgccaccATGCACCATCACCATCACCATTCCGGCGATTACAAAGACGATGA<br>CGATAAGATGGCGCCTAAGAAGAAACGCAAAGTGCGGGGCATGAAGAAAGAAAAGATTGATCTGTTTTACG<br>GAGCCCTGCTGGCCGCCATCGGAAAGGTCATCCAGCGAGCAACCGGAGAGCGGAAGAAACACGCACTTGTG<br>GGCGCCGACTGGTTCGACGAGATCGCCGACAACCAAGTCATCTCGGATCAGATCCGGTACCATATGGCCAA<br>CTACCAGTCTGATAAGCTCGGCAACGATCACCTGGCTTACATCACCTACATTGCCGACAACATCGCCTCCG<br>GTGTCGACCGCCGGCAATCCAACGAAGAGTCAGACGAAGATACCTCCGCAAAGATCTGGGACACCTACACG<br>AACCAGGCCGACATCTTTAACGTGTTCGGAGCGCAGACCGATAAGCGGTACTTCAAGCCTACCGTGCTGAA<br>TCTCAAGTCGAAGCCCAACTTCGCGTCCGCCACTTACGAACCCTTTAGCAAGGGCGATTACGCTGCCATCG<br>CCACCCGGATTAAGAACGAACTGGCCGAGTTCGAGTTCAACCAAGTCCAGATTGACTCCCTGCTCAACCTT<br>TTCGAGGCTACTCTCTCCTTCGTGCCGTCAAGCACCAACACTAAGGAAATCGCCGACATCTCCCTGGCCGA<br>CCATTCCCGCTTGACTGCTGCCTTCGCTCTGGCGATCTACGACTACCTGGAGGACAAGGGTCGGCACAACT<br>ACAAAGAGGACCTGTTCACCAAAGTGTCAGCGTTCTATGAAGAAGAAGCCTTCCTGCTGGCCTCCTTCGAC<br>CTGTCGGGAATCCAGGACTTTATCTACAACATTAACATCGCAACTAACGGCGCGGCGAAGCAGCTGAAGGC<br>CCGGAGCCTCTACCTGGACTTTATGTCCGAGTACATCGCCGATAGCCTGCTGGACAAGCTGGGACTGAACA<br>GGGCTAACATGCTTTACGTCGGCGGCGGACACGCCTACTTCGTCCTGGCCAACACCGAAAAGACTGTGGAA<br>ACCCTGGTGCAGTTTGAGAAGGATTTCAACCAGTTCCTGTTGGCAAACTTCCAGACCCGCCTCTATGTGGC<br>CTTTGGCTGGGGTTCCTTCGCGGCCAAGGACATCATGTCCGAGCTGAATAGCCCCGAGTCCTACCGCCAAG<br>TGTACCAAAAGGCTTCGCGCATGATCTCCAAAAAGAAAATCTCCAGATACGACTACCAGACACTGATGCTC<br>CTGAATCGCGGTGGAAAGTCCTCAGAGAGAGAGTGCGAGATTTGCCACTCCGTGGAGAACCTGGTGTCCTA<br>CCACGACCAGAAAGTCTGTGACATTTGCCGGGGACTGTACCAGTTCTCGAAAGAAATTGCCCATGACCACT<br>TCATCATTACCGAAAATGAGGGGCTGCCGATTGGACCAAACGCGTGCTTAAAGGGCGTGGCATTCGAAAAG<br>CTGTCCCAAGAAGCGTTCAGCCGGGTCTACGTGAAGAATGACTATAAGGCCGGTACCGTGAAGGCTACGCA<br>TGTGTTCGTGGGGGATTACCAGTGCGACGAGATCTACAACTACGCCGCCCTGAGCAAGAACGAGAACGGCC<br>TAGGCATCAAGAGACTGGCCGTGGTCCGGCTCGACGTGGATGACTTGGGCGCCGCCTTCATGGCCGGTTTC<br>AGCCAGCAGGGAAACGGACAATACTCCACTCTGTCAAGATCGGCCACATTCTCCCGGAGCATGTCGCTGTT<br>CTTCAAAGTGTACATTAACCAGTTCGCCTCCGACAAGAAGCTGAGCATTATCTACGCGGGCGGCGCCGCCG |

Table S2

| | |
|---|---|
| | TGTTCGCCATTGGATCGTGGCAGGATATCATCGCGTTCACTGTGGAACTTCGCGAAAACTTCATCAAGTGG<br>ACCAACGGGAAGCTCACCCTCTCCGCGGGGATAGGGTTGTTCGCCGACAAGACTCCTATTAGCCTGATGGC<br>TCACCAGACCGGGGAACTGGAAGAGGCCGCCAAGGGCAACGAAAAGGACTCCATCTCGCTGTTCTCAAGCG<br>ACTACACTTTCAAGTTTGATAGGTTCATCACTAACGTGTACGACGACAAACTGGAACAGATTAGATACTTC<br>TTCAACCATCAAGACGAGAGGGGAAAGAACTTCATCTATAAGCTTATTGAGCTTTTGAGGAACCACGACCG<br>CATGAATATGGCACGCCTCGCCTATTACCTCACTCGCCTGGAAGAACTGACCCGGGAGACTGACAGGGACA<br>AGTTCAAGACCTTCAAGAACCTGTTCTACTCCTGGTACACCAACAAGAACGATAAGGACCGGAAGGAAGCC<br>GAGCTCGCGCTCCTGCTGTACATCTACGAAATCAGAAAGGATTAAcggcaataaaaagacagaataaaacg<br>cacggtgttgggtcgtttgttcGCACACATTAGCTAGCCGTCAGCACACATTGTGATGCGGTTTTGGCAGT<br>ACATCAATGGGCGTGGATAGCGGTTTGACTCACGGGGATTTCCAAGTCTCCACCCCATTGACGTCAATGGG<br>AGTTTGTTTTGGCACCAAAATCAACGGGACTTTCCAAAATGTCGTAACAACTCCGCCCCATTGACGCAAAT<br>GGGCGGTAGGCGTGTACGGTGGGAGGTCTATATAAGCAGAGCTCGTTTAGTGAACCGTCAGATCTCTAGAg<br>ccgccaccATGGATTACAAAGACGATGACGATAAGATGGCGCCTAAGAAGAAACGCAAAGTGCGGGGCATG<br>ACCATCCTGACCGACGAGAACTACGTGGACATCGCCGAGAAAGCCATCCTGAAGCTGGAAAGAAACACCAG<br>AAATAGAAAGAACCCTGATGCCTTCTTCCTGACCACATCTAAGCTGCGGAACCTGCTGAGCCTGACAAGCA<br>CCCTGTTCGACGAGAGCAAGGTGAAGGAATACGACGCCCTGCTGGACAGAATCGCTTATCTGAGAGTGCAG<br>TTCGTGTACCAGGCCGGCAGAGAGATCGCCGTGAAAGATCTGATCGAGAAGGCCCAGATCCTGGAAGCTCT<br>GAAAGAGATCAAGGACCGGGAAACCCTGCAGAGATTCTGCAGATACATGGAAGCCCTGGTGGCCTACTTCA<br>AGTTCTACGGCGGCAAGGACTGAcggcaataaaaagacagaataaaacgcacggtgttgggtcgtttgttc<br>AGTTCTTTTGCCTTACTTTCAATGCATGCGGTGATGCGGTTTTGGCAGTACATCAATGGGCGTGGATAGCG<br>GTTTGACTCACGGGGATTTCCAAGTCTCCACCCCATTGACGTCAATGGGAGTTTGTTTTGGCACCAAAATC<br>AACGGGACTTTCCAAAATGTCGTAACAACTCCGCCCCATTGACGCAAATGGGCGGTAGGCGTGTACGGTGG<br>GAGGTCTATATAAGCAGAGCTCGTTTAGTGAACCGTCAGATCTCTAGAgccgccaccATGGATTACAAAGA<br>CGATGACGATAAGATGGCGCCTAAGAAGAAACGCAAAGTGCGGGGCATGACCTTCGCCAAGATCAAATTCA<br>GCGCCCAGATCCGGCTGGAAACCGGCCTGCACATCGGAGGATCTGATGCCTTTGCCGCTATCGGCGCCATC<br>GCCAGCCCTGTGATCAAGGACCCCATCACCAACCTGCCTATCATCCCCGGCTCTAGCCTGAAGGGCAAGAT<br>GAGAACACTGCTGGCCAAGGTGTACAACGAAAAGGTGGCCGAGAAGCCTAGCGACGACAGCGACATCCTGA<br>GCAGACTGTTCGGAAATAGCAAGGATAAGCGGTTCAAGATGGGCAGACTGATCTTCCGGGACGCCTTCCTG<br>AGCAACGCCGACGAGCTGGATTCTCTGGGCGTGCGGAGCTACACCGAGGTGAAGTTCGAGAACACCATCGA<br>TAGAATCACCGCCGAGGCCAATCCTAGACAGATCGAGAGAGCCATTCGGAACTCAACATTCGACTTCGAGC<br>TGATCTACGAGATCACTGATGAGAATGAGAACCAGGTCGAGGAAGATTTCAAGGTGATCAGAGACGGCCTG<br>AAGCTGCTGGAACTGGACTACCTGGGCGGAAGCGGCTCCAGAGGCTACGGCAAAGTGGCTTTTGAGAACCT<br>GAAAGCCACCACAGTGTTCGGCAACTACGACGTGAAAACCCTGAACGAGCTGCTGACCGCCGAAGTGGGCG<br>GAGGTGCTGCCGCAAtggtgagcaagggcgaggagctgttcaccggggtggtgcccatcctggtcgagctg<br>gacggcgacgtaaacggccacaagttcagcgtgtccggcgagggcgagggcgatgccacctacggcaagct<br>gaccctgaagttcatctgcaccaccggcaagctgcccgtgccctggcccaccctcgtgaccaccctgacct<br>acggcgtgcagtgcttcagccgctaccccgaccacatgaagcagcacgacttcttcaagtccgccatgccc<br>gaaggctacgtccaggagcgcaccatcttcttcaaggacgacggcaactacaagacccgcgccgaggtgaa<br>gttcgagggcgacaccctggtgaaccgcatcgagctgaagggcatcgacttcaaggaggacggcaacatcc<br>tggggcacaagctggagtacaactacaacagccacaacgtctatatcatggccgacaagcagaagaacggc<br>atcaaggtgaacttcaagatccgccacaacatcgaggacggcagcgtgcagctcgccgaccactaccagca<br>gaacacccccatcggcgacggccccgtgctgctgcccgacaaccactacctgagcacccagtccgccctga<br>gcaaagaccccaacgagaagcgcgatcacatggtcctgctggagttcgtgaccgccgccgggatcactctc<br>ggcatggacgagctgtacaagtgacggcaataaaaagacagaataaaacgcacggtgttgggtcgtttgtt<br>cTGGATTGCGAGAATGGACTAGTAGCAAACTGTGATGCGGTTTTGGCAGTACATCAATGGGCGTGGATAGC<br>GGTTTGACTCACGGGGATTTCCAAGTCTCCACCCCATTGACGTCAATGGGAGTTTGTTTTGGCACCAAAAT<br>CAACGGGACTTTCCAAAATGTCGTAACAACTCCGCCCCATTGACGCAAATGGGCGGTAGGCGTGTACGGTG<br>GGAGGTCTATATAAGCAGAGCTCGTTTAGTGAACCGTCAGATCTCTAGAgccgccaccATGGATTACAAAG<br>ACGATGACGATAAGATGGCGCCTAAGAAGAAACGCAAAGTGCGGGGCATGACTTACAAGCTCTACATTATG<br>ACCTTTCAAAACGCCCACTTCGGTTCCGGCACTCTGGACTCATCGAAGCTGACCTTCTCCGCGGATAGAAT<br>CTTCTCGGCACTCGTGCTCGAGGCTCTGAAGATGGGAAAGCTCGACGCCTTCTTGGCCGAGGCCAACCAGG<br>ATAAGTTCACTCTGACCGACGCGTTCCCATTCCAATTCGGTCCTTTCCTGCCGAAACCGATTGGTTACCCC<br>AAGCACGACCAGATCGACCAGTCTGTGGACGTGAAGGAAGTCCGCCGCCAAGCGAAGCTGTCCAAAAAGCT<br>CCAGTTCCTGGCTCTGGAAAACGTCGACGACTACCTGAACGGAGAGCTGTTTGAGAATGAGGAACACGCCG<br>TGATCGACACAGTGACCAAGAACCAGCCCCATAAAGATGATAATCTGTACCAAGTGGCCACCACTCGGTTC<br>TCGAACGACACCTCCCTTTACGTGATCGCCAACGAATCCGATCTGCTGAACGAACTGATGAGCAGCCTTCA<br>GTACTCCGGGCTGGGCGGCAAAAGGTCCTCAGGATTCGGCAGATTTGAGCTGGACATCCAGAACATTCCCT<br>TGGAACTGTCCGACCGGCTGACGAAGAACCACAGCGACAAGGTCATGTCACTTACCACCGCCCTCCCGGTG<br>GACGCTGATCTCGAGGAAGCGATGGAAGATGGCCATTACCTGTTGACCAAGTCGTCCGGATTCGCATTCTC<br>CCACGCCACCAACGAAAACTATCGGAAGCAGGACCTGTACAAGTTCGCCTCCGGGAGCACCTTCAGCAAGA<br>CTTTCGAGGGACAGATCGTGGACGTGCGCCCTCTCGATTTCCCTCACGCCGTGCTGAACTACGCCAAGCCG<br>CTGTTCTTTAAGCTCGAAGTCTAAcggcaataaaaagacagaataaaacgcacggtgttgggtcgtttgtt<br>cGCACATTCAAAAACAGGCAATTGGACAAGCGTGATGCGGTTTTGGCAGTACATCAATGGGCGTGGATAGC |

Table S2

| | |
|---|---|
| | GGTTTGACTCACGGGGATTTCCAAGTCTCCACCCCATTGACGTCAATGGGAGTTTGTTTTGGCACCAAAAT<br>CAACGGGACTTTCCAAAATGTCGTAACAACTCCGCCCCATTGACGCAAATGGGCGGTAGGCGTGTACGGTG<br>GGAGGTCTATATAAGCAGAGCTCGTTTAGTGAACCGTCAGATCTCTAGAgccgccaccATGGATTACAAAG<br>ACGATGACGATAAGATGGCGCCTAAGAAGAAACGCAAAGTGCGGGGCATGAAAAATGACTACCGGACCTTC<br>AAGCTGAGCCTGCTGACCCTGGCTCCTATCCACATCGGCAACGGCGAGAAGTACACCAGCAGAGAATTCAT<br>CTACGAGAACAAGAAGTTCTACTTCCCCGACATGGGCAAGTTCTACAACAAGATGGTGGAAAAGAGACTGG<br>CCGAGAAGTTCGAGGCCTTCCTGATCCAGACCAGACCCAACGCCAGAAACAACCGGCTGATTTCTTTTCTG<br>AACGACAACAGAATCGCCGAAAGATCTTTTGGCGGCTACAGCATCAGTGAAACCGGCCTGGAATCTGATAA<br>GAACCCTAACAGCGCCGGAGCTATCAACGAGGTGAACAAATTCATCCGGGACGCCTTCGGAAATCCTTACA<br>TCCCAGGCAGCAGCCTGAAGGGCGCCATCCGCACCATCCTGATGAACACCACACCTAAGTGGAACAACGAG<br>AACGCCGTGAACGACTTCGGCAGATTCCCAAAGGAAAACAAGAACCTGATCCCTTGGGGACCTAAGAAAGG<br>CAAGGAATACGACGACCTGTTCAACGCCATCAGAGTGTCCGACAGCAAGCCCTTCGACAACAAAAGCCTGA<br>TCCTCGTGCAGAAGTGGGACTACAGCGCCAAAACCAACAAGGCCAAGCCTCTGCCTCTGTACAGAGAGTCT<br>ATCAGCCCTCTGACCAAGATCGAGTTCGAGATAACAACAACCACTGATGAGGCCGGCAGACTGATCGAGGA<br>ACTGGGAAAGCGGGCCCAGGCCTTTTATAAGGACTACAAGGCCTTTTTCCTGTCTGAATTCCCTGATGATA<br>AGATCCAGGCTAATCTGCAATACCCCATCTACCTGGGCGCCGGCAGCGGCGCTTGGACAAAGACCCTGTTT<br>AAGCAGGCCGACGGCATCCTGCAGCGGAGATACTCCAGAATGAAAACCAAGATGGTCAAGAAGGGCGTGCT<br>GAAGCTGACAAAGGCCCCTCTGAAAACAGTGAAGATCCCCAGCGGCAACCACAGCCTGGTGAAGAATCACG<br>AGAGCTTCTACGAGATGGGCAAAGCCAACTTCATGATCAAGGAAATCGACAAGTGAcggcaataaaaagac<br>agaataaaacgcacggtgttgggtcgtttgttcATGTGTGCCGGCCGGAGAATAAAAGTCTAAGTGATGCG<br>GTTTTGGCAGTACATCAATGGGCGTGGATAGCGGTTTGACTCACGGGGATTTCCAAGTCTCCACCCCATTG<br>ACGTCAATGGGAGTTTGTTTTGGCACCAAAATCAACGGGACTTTCCAAAATGTCGTAACAACTCCGCCCCA<br>TTGACGCAAATGGGCGGTAGGCGTGTACGGTGGGAGGTCTATATAAGCAGAGCTCGTTTAGTGAACCGTCA<br>GATCTCTAGAgccgccaccATGGATTACAAAGACGATGACGATAAGATGGCGCCTAAGAAGAAACGCAAAG<br>TGCGGGGCATGAAAAAGCTCGTGTTCACCTTTAAGCGGATCGACCACCCTGCTCAGGACCTGGCCGTGAAA<br>TTCCACGGCTTCCTGATGGAACAGCTGGATAGCGACTACGTGGACTACCTGCACCAGCAGCAGACCAACCC<br>CTACGCCACAAAGGTGATCCAGGGCAAAGAGAACACCCAGTGGGTCGTGCATCTGCTGACAGACGACATCG<br>AGGACAAGGTGTTCATGACCCTGCTGCAGATCAAGGAAGTGTCCCTGAACGACCTGCCTAAGTTGTCTGTG<br>GAAAAGGTGGAAATCCAGGAGCTGGGCGCTGATAAGCTGCTCGAGATCTTCAACAGCGAGGAAAACCAGAC<br>CTACTTCAGCATCATCTTCGAGACACCTACAGGCTTTAAAAGCCAGGGCAGCTACGTGATCTTCCCCAGCA<br>TGCGGCTGATCTTTCAGAGCCTGATGCAGAAGTACGGCAGACTGGTGGAAAACCAGCCTGAGATCGAGGAA<br>GATACCCTGGACTACCTGAGCGAGCACAGCACCATCACCAATTACAGACTGGAAACAAGCTACTTCAGAGT<br>GCATAGACAGAGAATCCCCGCCTTCCGGGGCAAGCTGACCTTCAAGGTGCAGGGAGCCCAGACACTGAAGG<br>CCTACGTGAAGATGCTGCTGACCTTCGGCGAGTACAGCGGCCTGGGCATGAAAACCAGCCTGGGAATGGGC<br>GGCATCAAGCTGGAAGAAAGAAAGGACTGAcggcaataaaaagacagaataaaacgcacggtgttgggtcg<br>tttgttcGTGCGATAGAGGGATCCAGTGCACCGGTTGGAGGGCCTATTTCCCATGATTCCTTCATATTTGC<br>ATATACGATACAAGGCTGTTAGAGAGATAATTAGAATTAATTTGACTGTAAACACAAAGATATTAGTACAA<br>AATACGTGACGTAGAAAGTAATAATTTCTTGGGTAGTTTGCAGTTTTAAAATTATGTTTTAAAATGGACTA<br>TCATATGCTTACCGTAACTTGAAAGTATTTCGATTTCTTGGCTTTATATATCTTGTGGAAAGGACGAAACA<br>CCGATATAAACCTAATTACCTCGAGAGGGGACGGAAACCCGTCTTCGATGAAGCGATTCAGAAGACTTGAT<br>ATAAACCTAATTACCTCGAGAGGGGACTTTTTTACATGTGTCAGAGGTTTTCACCGTCATCACCGAAACGC<br>GCGAGACGAAAGGGCCTCGTGATACGCCTATTTTTATAGGTTAATGTCATGATAATAATGGTTTCTTAGAC<br>GTCAGGTGGCACTTTTCGGGGAAATGTGCGCGGAACCCCTATTTGTTTATTTTTCTAAATACATTCAAATA<br>TGTATCCGCTCATGAGACAATAACCCTGATAAATGCTTCAATAATATTGAAAAAGGAAGAGTATGAGTATT<br>CAACATTTCCGTGTCGCCCTTATTCCCTTTTTTGCGGCATTTTGCCTTCCTGTTTTTGCTCACCCAGAAAC<br>GCTGGTGAAAGTAAAAGATGCTGAAGATCAGTTGGGTGCACGAGTGGGTTACATCGAACTGGATCTCAACA<br>GCGGTAAGATCCTTGAGAGTTTTCGCCCCGAAGAACGTTTTCCAATGATGAGCACTTTTAAAGTTCTGCTA<br>TGTGGCGCGGTATTATCCCGTATTGACGCCGGGCAAGAGCAACTCGGTCGCCGCATACACTATTCTCAGAA<br>TGACTTGGTTGAGTACTCACCAGTCACAGAAAAGCATCTTACGGATGGCATGACAGTAAGAGAATTATGCA<br>GTGCTGCCATAACCATGAGTGATAACACTGCGGCCAACTTACTTCTGACAACGATCGGAGGACCGAAGGAG<br>CTAACCGCTTTTTTGCACAACATGGGGGATCATGTAACTCGCCTTGATCGTTGGGAACCGGAGCTGAATGA<br>AGCCATACCAAACGACGAGCGTGACACCACGATGCCTGTAGCAATGGCAACAACGTTGCGCAAACTATTAA<br>CTGGCGAACTACTTACTCTAGCTTCCCGGCAACAATTAATAGACTGGATGGAGGCGGATAAAGTTGCAGGA<br>CCACTTCTGCGCTCGGCCCTTCCGGCTGGCTGGTTTATTGCTGATAAATCTGGAGCCGGTGAGCGTGGGTC<br>TCGCGGTATCATTGCAGCACTGGGGCCAGATGGTAAGCCCTCCCGTATCGTAGTTATCTACACGACGGGGA<br>GTCAGGCAACTATGGATGAACGAAATAGACAGATCGCTGAGATAGGTGCCTCACTGATTAAGCATTGGTAA<br>CTGTCAGACCAAGTTTACTCATATATACTTTAGATTGATTTAAAACTTCATTTTTAATTTAAAAGGATCTA<br>GGTGAAGATCCTTTTTGATAATCTCATGACCAAAATCCCTTAACGTGAGTTTTCGTTCCACTGAGCGTCAG<br>ACCCCGTAGAAAAGATCAAAGGATCTTCTTGAGATCCTTTTTTTCTGCGCGTAATCTGCTGCTTGCAAACA<br>AAAAAACCACCGCTACCAGCGGTGGTTTGTTTGCCGGATCAAGAGCTACCAACTCTTTTTCCGAAGGTAAC<br>TGGCTTCAGCAGAGCGCAGATACCAAATACTGTTCTTCTAGTGTAGCCGTAGTTAGGCCACCACTTCAAGA<br>ACTCTGTAGCACCGCCTACATACCTCGCTCTGCTAATCCTGTTACCAGTGGCTGCTGCCAGTGGCGATAAG<br>TCGTGTCTTACCGGGTTGGACTCAAGACGATAGTTACCGGATAAGGCGCAGCGGTCGGGCTGAACGGGGGG |

Table S2

| | |
|---|---|
| | TTCGTGCACACAGCCCAGCTTGGAGCGAACGACCTACACCGAACTGAGATACCTACAGCGTGAGCTATGAG<br>AAAGCGCCACGCTTCCCGAAGGGAGAAAGGCGGACAGGTATCCGGTAAGCGGCAGGGTCGGAACAGGAGAG<br>CGCACGAGGGAGCTTCCAGGGGGAAACGCCTGGTATCTTTATAGTCCTGTCGGGTTTCGCCACCTCTGACT<br>TGAGCGTCGATTTTTGTGATGCTCGTCAGGGGGGCGGAGCCTATGGAAAAACGCCAGCAACGCGGCCTTTT<br>TCTTAAGC |

| **Plasmid** | **pDAC689** |
|---|---|
| **Description** | Expression of Cas13; RFP backbone |
| **Utility** | RNA KD |
| **Features** | Pcmv-NLS-Cas13-NLS-pA; Pu6-crRNA-pT; Pcmv-RFP-pA |
| **Sequence** | GTGATGCGGTTTTGGCAGTACATCAATGGGCGTGGATAGCGGTTTGACTCACGGGGATTTCCAAGTCTCCA<br>CCCCATTGACGTCAATGGGAGTTTGTTTTGGCACCAAAATCAACGGGACTTTCCAAAATGTCGTAACAACT<br>CCGCCCCATTGACGCAAATGGGCGGTAGGCGTGTACGGTGGGAGGTCTATATAAGCAGAGCTCGTTTAGTG<br>AACCGTCAGATCTCTAGAgccgccaccATGcccaagaagaagagaaaggtggaggccagcatcgagaagaa<br>gaagagcttcgccaagggcatgggagtgaagagcaccctggtgtccggctctaaggtgtacatgaccacat<br>ttgctgagggaagcgacgccaggctggagaagatcgtggagggcgatagcatcagatccgtgaacgaggga<br>gaggctttcagcgccgagatggctgacaagaacgctggctacaagatcggaaacgccaagttttcccaccc<br>aaagggctacgccgtggtggctaacaacccactgtacaccggaccagtgcagcaggacatgctgggactga<br>aggagacactggagaagaggtacttcggcgagtccgccgacggaaacgataacatctgcatccaggtcatc<br>cacaacatcctggatatcgagaagatcctggctgagtacatcacaaacgccgcttacgccgtgaacaacat<br>ctccggcctggacaaggatatcatcggcttcggaaagttttctaccgtgtacacatacgacgagttcaagg<br>atccagagcaccaccggccgcttttaacaacaacgacaagctgatcaacgccatcaaggctcagtacgac<br>gagttcgataactttctggataaccccaggctgggctacttcggacaggctttcttttctaaggagggcag<br>aaactacatcatcaactacggaaacgagtgttacgatatcctggcgctgctgagcggactgaggcactggg<br>tggtgcacaacaacgaggaggagtctcggatctctcgcacctggctgtacaacctggacaagaacctggat<br>aacgagtacatctccacactgaactacctgtacgacaggatcaccaacgagctgacaaacagcttctccaa<br>gaactctgccgctaacgtgaactacatcgctgagaccctgggcatcaacccagctgagttcgctgagcagt<br>acttcagattttccatcatgaaggagcagaagaacctgggcttcaacatcacaaagctgagagaagtgatg<br>ctggacagaaaggatatgtccgagatcaggaagaaccacaaggtgttcgattctatcagaaccaaggtgta<br>cacaatgatggactttgtgatctacaggtactacatcgaggaggatgccaaggtggccgctgccaacaaga<br>gcctgcccgacaacgagaagtctctgagcgagaaggatatcttcgtgatcaacctgagaggctcctttaac<br>gacgatcagaaggacgctctgtactacgatgaggccaacaggatctggagaaagctggagaacatcatgca<br>caacatcaaggagttccggggaaacaagacccgcgagtacaagaagaaggacgctccaaggctgcctagga<br>tcctgcctgctggaagggacgtgagcgccttcagcaagctgatgtacgccctgacaatgtttctggacgga<br>aaggagatcaacgatctgctgaccacactgatcaacaagttcgacaacatccagtcttttctgaaagtgat<br>gcctctgatcggcgtgaacgctaagttcgtggaggagtacgccttctttaaggacagcgccaagatcgctg<br>atgagctgcggctgatcaagtcctttgccaggatgggagagccaatcgctgacgctaggagagctatgtac<br>atcgatgccatccggatcctgggaaccaacctgtcttacgacgagctgaaggctctggccgacaccttcag<br>cctggatgagaacggcaacaagctgaagaagggcaagcacggaatgcgcaacttcatcatcaacaacgtga<br>tcagcaacaagcggtttcactacctgatcagatacggcgacccagctcacctgcacgagatcgctaagaac<br>gaggccgtggtgaagttcgtgctgggacggatcgccgatatccagaagaagcagggccagaacggaaagaa<br>ccagatcgaccgctactacgagacctgcatcggcaaggataagggaaagtccgtgtctgagaaggtggacg<br>ctctgaccaagatcatcacaggcatgaactacgaccagttcgataagaagagatctgtgatcgaggacacc<br>ggaagggagaacgccgagagagagaagtttaagaagatcatcagcctgtacctgacagtgatctaccacat<br>cctgaagaacatcgtgaacatcaacgctagatacgtgatcggcttccactgcgtggagcgcgatgcccagc<br>tgtacaaggagaagggatacgacatcaacctgaagaagctggaggagaagggctttagctccgtgaccaag<br>ctgtgcgctggaatcgacgagacagcccccgacaagaggaaggatgtggagaaggagatggccgagagagc<br>taaggagagcatcgactccctggagtctgctaaccctaagctgtacgccaactacatcaagtactccgatg<br>agaagaaggccgaggagtttaccaggcagatcaacagagagaaggccaagaccgctctgaacgcctacctg<br>aggaacacaaagtggaacgtgatcatccgggaggacctgctgcgcatcgataacaagacctgtacactgtt<br>ccggaacaaggctgtgcacctggaggtggctcgctacgtgcacgcctacatcaacgacatcgccgaggtga<br>actcctactttcagctgtaccactacatcatgcagaggatcatcatgaacgagagatacgagaagtctagc<br>ggcaaggtgtctgagtacttcgacgccgtgaacgatgagaagaagtacaacgatagactgctgaagctgct<br>gtgcgtgcctttcggatactgtatcccacggtttaagaacctgagcatcgaggccctgttcgaccgcaacg<br>aggctgccaagtttgataaggagaagaagaaggtgagcggcaactccggatccggacctaagaaaaagagg<br>aaggtgtgacggcaataaaaagacagaataaaacgcacggtgttgggtcgtttgttcGTGCGATAGAGGGA<br>TCCCGCATTGAATTATGTGATGCGGTTTTGGCAGTACATCAATGGGCGTGGATAGCGGTTTGACTCACGGG<br>GATTTCCAAGTCTCCACCCCATTGACGTCAATGGGAGTTTGTTTTGGCACCAAAATCAACGGGACTTTCCA<br>AAATGTCGTAACAACTCCGCCCCATTGACGCAAATGGGCGGTAGGCGTGTACGGTGGGAGGTCTATATAAG<br>CAGAGCTCGTTTAGTGAACCGTCAGATCTCTAGAgccgccaccATGGTGAGCAAGGGCGAGGAGGATAACA<br>TGGCCATCATCAAGGAGTTCATGCGCTTCAAGGTGCACATGGAGGGCTCCGTGAACGGCCACGAGTTCGAG<br>ATCGAGGGCGAGGGCGAGGGCCGCCCCTACGAGGGCACCCAGACCGCCAAGCTGAAGGTGACCAAGGGTGG<br>CCCCCTGCCCTTCGCCTGGGACATCCTGTCCCCTCAGTTCATGTACGGCTCCAAGGCCTACGTGAAGCACC |

Table S2

| | |
|---|---|
| | CCGCCGACATCCCCGACTACTTGAAGCTGTCCTTCCCCGAGGGCTTCAAGTGGGAGCGCGTGATGAACTTC<br>GAGGACGGCGGCGTGGTGACCGTGACCCAGGACTCCTCCCTGCAGGACGGCGAGTTCATCTACAAGGTGAA<br>GCTGCGCGGCACCAACTTCCCCTCCGACGGCCCCGTAATGCAGAAGAAAACCATGGGCTGGGAGGCCTCCT<br>CCGAGCGGATGTACCCCGAGGACGGCGCCCTGAAGGGCGAGATCAAGCAGAGGCTGAAGCTGAAGGACGGC<br>GGCCACTACGACGCTGAGGTCAAGACCACCTACAAGGCCAAGAAGCCCGTGCAGCTGCCCGGCGCCTACAA<br>CGTCAACATCAAGTTGGACATCACCTCCCACAACGAGGACTACACCATCGTGGAACAGTACGAACGCGCCG<br>AGGGCCGCCACTCCACCGGCGGCATGGACGAGCTGTACAAGTAAcggcaataaaaagacagaataaaacgc<br>acggtgttgggtcgtttgttcGAGCAGATTGTACTGAGAGTGCACCGGTTGGAGGGCCTATTTCCCATGAT<br>TCCTTCATATTTGCATATACGATACAAGGCTGTTAGAGAGATAATTAGAATTAATTTGACTGTAAACACAA<br>AGATATTAGTACAAAATACGTGACGTAGAAAGTAATAATTTCTTGGGTAGTTTGCAGTTTTAAAATTATGT<br>TTTAAAATGGACTATCATATGCTTACCGTAACTTGAAAGTATTTCGATTTCTTGGCTTTATATATCTTGTG<br>GAAAGGACGAAACACCGcaagtaaacccctaccaactggtcggggtttgaaacgggtcttcgagaagaccc<br>TTTTTTACATGTGTCAGAGGTTTTCACCGTCATCACCGAAACGCGCGAGACGAAAGGGCCTCGTGATACGC<br>CTATTTTTATAGGTTAATGTCATGATAATAATGGTTTCTTAGACGTCAGGTGGCACTTTTCGGGGAAATGT<br>GCGCGGAACCCCTATTTGTTTATTTTTCTAAATACATTCAAATATGTATCCGCTCATGAGACAATAACCCT<br>GATAAATGCTTCAATAATATTGAAAAAGGAAGAGTATGAGTATTCAACATTTCCGTGTCGCCCTTATTCCC<br>TTTTTTGCGGCATTTTGCCTTCCTGTTTTTGCTCACCCAGAAACGCTGGTGAAAGTAAAAGATGCTGAAGA<br>TCAGTTGGGTGCACGAGTGGGTTACATCGAACTGGATCTCAACAGCGGTAAGATCCTTGAGAGTTTTCGCC<br>CCGAAGAACGTTTTCCAATGATGAGCACTTTTAAAGTTCTGCTATGTGGCGCGGTATTATCCCGTATTGAC<br>GCCGGGCAAGAGCAACTCGGTCGCCGCATACACTATTCTCAGAATGACTTGGTTGAGTACTCACCAGTCAC<br>AGAAAAGCATCTTACGGATGGCATGACAGTAAGAGAATTATGCAGTGCTGCCATAACCATGAGTGATAACA<br>CTGCGGCCAACTTACTTCTGACAACGATCGGAGGACCGAAGGAGCTAACCGCTTTTTTGCACAACATGGGG<br>GATCATGTAACTCGCCTTGATCGTTGGGAACCGGAGCTGAATGAAGCCATACCAAACGACGAGCGTGACAC<br>CACGATGCCTGTAGCAATGGCAACAACGTTGCGCAAACTATTAACTGGCGAACTACTTACTCTAGCTTCCC<br>GGCAACAATTAATAGACTGGATGGAGGCGGATAAAGTTGCAGGACCACTTCTGCGCTCGGCCCTTCCGGCT<br>GGCTGGTTTATTGCTGATAAATCTGGAGCCGGTGAGCGTGGGTCTCGCGGTATCATTGCAGCACTGGGGCC<br>AGATGGTAAGCCCTCCCGTATCGTAGTTATCTACACGACGGGGAGTCAGGCAACTATGGATGAACGAAATA<br>GACAGATCGCTGAGATAGGTGCCTCACTGATTAAGCATTGGTAACTGTCAGACCAAGTTTACTCATATATA<br>CTTTAGATTGATTTAAAACTTCATTTTTAATTTAAAAGGATCTAGGTGAAGATCCTTTTTGATAATCTCAT<br>GACCAAAATCCCTTAACGTGAGTTTTCGTTCCACTGAGCGTCAGACCCCGTAGAAAAGATCAAAGGATCTT<br>CTTGAGATCCTTTTTTTCTGCGCGTAATCTGCTGCTTGCAAACAAAAAAACCACCGCTACCAGCGGTGGTT<br>TGTTTGCCGGATCAAGAGCTACCAACTCTTTTTCCGAAGGTAACTGGCTTCAGCAGAGCGCAGATACCAAA<br>TACTGTTCTTCTAGTGTAGCCGTAGTTAGGCCACCACTTCAAGAACTCTGTAGCACCGCCTACATACCTCG<br>CTCTGCTAATCCTGTTACCAGTGGCTGCTGCCAGTGGCGATAAGTCGTGTCTTACCGGGTTGGACTCAAGA<br>CGATAGTTACCGGATAAGGCGCAGCGGTCGGGCTGAACGGGGGGTTCGTGCACACAGCCCAGCTTGGAGCG<br>AACGACCTACACCGAACTGAGATACCTACAGCGTGAGCTATGAGAAAGCGCCACGCTTCCCGAAGGGAGAA<br>AGGCGGACAGGTATCCGGTAAGCGGCAGGGTCGGAACAGGAGAGCGCACGAGGGAGCTTCCAGGGGGAAAC<br>GCCTGGTATCTTTATAGTCCTGTCGGGTTTCGCCACCTCTGACTTGAGCGTCGATTTTTGTGATGCTCGTC<br>AGGGGGGCGGAGCCTATGGAAAAACGCCAGCAACGCGGCCTTTTTCTTAAGC |

| **Plasmid** | **pDAC690** |
|---|---|
| **Description** | Expression of shRNA; RFP backbone |
| **Utility** | RNA KD |
| **Features** | Pu6-shRNA-pT; Pcmv-RFP-pA |
| **Sequence** | GTGATGCGGTTTTGGCAGTACATCAATGGGCGTGGATAGCGGTTTGACTCACGGGGATTTCCAAGTCTCCA<br>CCCCATTGACGTCAATGGGAGTTTGTTTTGGCACCAAAATCAACGGGACTTTCCAAAATGTCGTAACAACT<br>CCGCCCCATTGACGCAAATGGGCGGTAGGCGTGTACGGTGGGAGGTCTATATAAGCAGAGCTCGTTTAGTG<br>AACCGTCAGATCTCTAGAgccgccaccATGGTGAGCAAGGGCGAGGAGGATAACATGGCCATCATCAAGGA<br>GTTCATGCGCTTCAAGGTGCACATGGAGGGCTCCGTGAACGGCCACGAGTTCGAGATCGAGGGCGAGGGCG<br>AGGGCCGCCCCTACGAGGGCACCCAGACCGCCAAGCTGAAGGTGACCAAGGGTGGCCCCCTGCCCTTCGCC<br>TGGGACATCCTGTCCCCTCAGTTCATGTACGGCTCCAAGGCCTACGTGAAGCACCCCGCCGACATCCCCGA<br>CTACTTGAAGCTGTCCTTCCCCGAGGGCTTCAAGTGGGAGCGCGTGATGAACTTCGAGGACGGCGGCGTGG<br>TGACCGTGACCCAGGACTCCTCCCTGCAGGACGGCGAGTTCATCTACAAGGTGAAGCTGCGCGGCACCAAC<br>TTCCCCTCCGACGGCCCCGTAATGCAGAAGAAAACCATGGGCTGGGAGGCCTCCTCCGAGCGGATGTACCC<br>CGAGGACGGCGCCCTGAAGGGCGAGATCAAGCAGAGGCTGAAGCTGAAGGACGGCGGCCACTACGACGCTG<br>AGGTCAAGACCACCTACAAGGCCAAGAAGCCCGTGCAGCTGCCCGGCGCCTACAACGTCAACATCAAGTTG<br>GACATCACCTCCCACAACGAGGACTACACCATCGTGGAACAGTACGAACGCGCCGAGGGCCGCCACTCCAC<br>CGGCGGCATGGACGAGCTGTACAAGTAAcggcaataaaaagacagaataaaacgcacggtgttgggtcgtt<br>tgttcGAGCAGATTGTACTGAGAGTGCACCGGTTGGAGGGCCTATTTCCCATGATTCCTTCATATTTGCAT<br>ATACGATACAAGGCTGTTAGAGAGATAATTAGAATTAATTTGACTGTAAACACAAAGATATTAGTACAAAA<br>TACGTGACGTAGAAAGTAATAATTTCTTGGGTAGTTTGCAGTTTTAAAATTATGTTTTAAAATGGACTATC<br>ATATGCTTACCGTAACTTGAAAGTATTTCGATTTCTTGGCTTTATATATCTTGTGGAAAGGACGAAACACC<br>GcgtcttcCCTGACCCAgaagacgCTTTTTTACATGTGTCAGAGGTTTTCACCGTCATCACCGAAACGCGC |

Table S2

| | |
|---|---|
| | GAGACGAAAGGGCCTCGTGATACGCCTATTTTTATAGGTTAATGTCATGATAATAATGGTTTCTTAGACGT<br>CAGGTGGCACTTTTCGGGGAAATGTGCGCGGAACCCCTATTTGTTTATTTTTCTAAATACATTCAAATATG<br>TATCCGCTCATGAGACAATAACCCTGATAAATGCTTCAATAATATTGAAAAAGGAAGAGTATGAGTATTCA<br>ACATTTCCGTGTCGCCCTTATTCCCTTTTTTGCGGCATTTTGCCTTCCTGTTTTTGCTCACCCAGAAACGC<br>TGGTGAAAGTAAAAGATGCTGAAGATCAGTTGGGTGCACGAGTGGGTTACATCGAACTGGATCTCAACAGC<br>GGTAAGATCCTTGAGAGTTTTCGCCCCGAAGAACGTTTTCCAATGATGAGCACTTTTAAAGTTCTGCTATG<br>TGGCGCGGTATTATCCCGTATTGACGCCGGGCAAGAGCAACTCGGTCGCCGCATACACTATTCTCAGAATG<br>ACTTGGTTGAGTACTCACCAGTCACAGAAAAGCATCTTACGGATGGCATGACAGTAAGAGAATTATGCAGT<br>GCTGCCATAACCATGAGTGATAACACTGCGGCCAACTTACTTCTGACAACGATCGGAGGACCGAAGGAGCT<br>AACCGCTTTTTTGCACAACATGGGGGATCATGTAACTCGCCTTGATCGTTGGGAACCGGAGCTGAATGAAG<br>CCATACCAAACGACGAGCGTGACACCACGATGCCTGTAGCAATGGCAACAACGTTGCGCAAACTATTAACT<br>GGCGAACTACTTACTCTAGCTTCCCGGCAACAATTAATAGACTGGATGGAGGCGGATAAAGTTGCAGGACC<br>ACTTCTGCGCTCGGCCCTTCCGGCTGGCTGGTTTATTGCTGATAAATCTGGAGCCGGTGAGCGTGGGTCTC<br>GCGGTATCATTGCAGCACTGGGGCCAGATGGTAAGCCCTCCCGTATCGTAGTTATCTACACGACGGGGAGT<br>CAGGCAACTATGGATGAACGAAATAGACAGATCGCTGAGATAGGTGCCTCACTGATTAAGCATTGGTAACT<br>GTCAGACCAAGTTTACTCATATATACTTTAGATTGATTTAAAACTTCATTTTTAATTTAAAAGGATCTAGG<br>TGAAGATCCTTTTTGATAATCTCATGACCAAAATCCCTTAACGTGAGTTTTCGTTCCACTGAGCGTCAGAC<br>CCCGTAGAAAAGATCAAAGGATCTTCTTGAGATCCTTTTTTTCTGCGCGTAATCTGCTGCTTGCAAACAAA<br>AAAACCACCGCTACCAGCGGTGGTTTGTTTGCCGGATCAAGAGCTACCAACTCTTTTTCCGAAGGTAACTG<br>GCTTCAGCAGAGCGCAGATACCAAATACTGTTCTTCTAGTGTAGCCGTAGTTAGGCCACCACTTCAAGAAC<br>TCTGTAGCACCGCCTACATACCTCGCTCTGCTAATCCTGTTACCAGTGGCTGCTGCCAGTGGCGATAAGTC<br>GTGTCTTACCGGGTTGGACTCAAGACGATAGTTACCGGATAAGGCGCAGCGGTCGGGCTGAACGGGGGGTT<br>CGTGCACACAGCCCAGCTTGGAGCGAACGACCTACACCGAACTGAGATACCTACAGCGTGAGCTATGAGAA<br>AGCGCCACGCTTCCCGAAGGGAGAAAGGCGGACAGGTATCCGGTAAGCGGCAGGGTCGGAACAGGAGAGCG<br>CACGAGGGAGCTTCCAGGGGGAAACGCCTGGTATCTTTATAGTCCTGTCGGGTTTCGCCACCTCTGACTTG<br>AGCGTCGATTTTTGTGATGCTCGTCAGGGGGGCGGAGCCTATGGAAAAACGCCAGCAACGCGGCCTTTTTC<br>TTAAGC |
